# Zwitterionic hydrogel designs for conducting polymers enable bioelectronics with suppressed foreign body responses

**DOI:** 10.1101/2025.10.31.683260

**Authors:** Shinya Wai, Seounghun Kang, Nan Li, Yahao Dai, Tera Lavoie, Joseph Strzalka, Sean Sutyak, Maximilian Weires, Tianda Fu, Jin Wang, Kaden C. Stevens, Ruojia Li, Jeffrey A. Hubbell, Matthew V. Tirrell, Sihong Wang

## Abstract

For long-term, continuous operation of implantable biosensors and electrophysiological devices, the foreign body response (FBR) is a major obstacle that needs to be overcome. As the FBR progresses, any implanted device will become damaged and isolated from its physiological environment, due to encapsulation by fibrotic tissue and inflammatory immune cells. To achieve more compatible and low-impedance biointerfaces, conducting polymers, such as PEDOT:PSS, have been extensively explored as ideal materials. However, FBR on such conducting polymers remains an unmet challenge. We report a zwitteronic-hydrogel-based double-network design for PEDOT:PSS that can significantly suppress the FBR by 64%, in addition to improving conductivity by more than one order of magnitude. Surprisingly, the FBR level of this design is even lower than that of the parent zwitteronic hydrogel by 53%. Our further immunological investigations at the histological, cellular, and transcriptomic levels give deeper insights into the unique effects that come from the chemical heterogeneity. Furthermore, chronic electrocardiographic recording in mice demonstrate the benefit of this material design to long-term, implanted electrophysiology, which provides indications for the future development of immunocompatible electronic polymers.

## Introduction

Implantable medical devices (IMDs), such as pacemakers, cochlear implants, and glucose monitors, are increasing in societal importance due to an aging population and rising chronic disease rates^1^. However, the foreign body response (FBR) has been a major barrier to their long-term function, which involves inflammation and fibrosis that can degrade and isolate devices through fibrotic encapsulation^2,3^. Fibrotic encapsulation is especially harmful for biosensors and electrophysiological devices, as it impedes analyte and ion transport, limiting their effectiveness and longevity in physiological environments^4^. The FBR is initiated by the adsorption of proteins on the implant surface, a process commonly known as fouling. This triggers an inflammatory response that leads to the recruitment of macrophages, which become a continuous, long-term source of degradative enzymes and chemicals such as reactive oxygen species. In addition, macrophages and other immune cells recruit fibroblasts that facilitate the formation of fibrotic tissue^2,3^. Even for IMDs that have been approved by the Food and Drug Administration (FDA) and in clinical use, e.g., pacemakers, the FBR is commonly still an issue that limits the longevity of the device function and causes additional side effects^5–7^. A number of approaches have been and are being pursued to reduce the FBR, yielding FBR-suppressing, i.e., immunocompatible properties^2–4,8^.

Strategies to suppress the foreign body response (FBR) in implantable medical devices (IMDs) generally fall into three categories: (i) mechanical matching to tissue^9,10^; (ii) controlled release of immunomodulatory drugs like dexamethasone^8^; and (iii) surface coatings and modifications to provide antibiofouling ^8,11,12^ or tissue adhesive effects^13,14^. While each has shown some success, they also face key limitations. Mechanical matching, even at tissue-level stiffness at a few kPa, can still trigger fibrosis^9,10^. Drug release strategies may cause systemic side effects^15,16^ and complicate fabrication, as many drugs are sensitive to heat and UV^17,18^. Surface coatings often suffer from long-term instability due to mechanical wear or chemical degradation and may hinder the transport of chemical species. Among coatings, zwitterionic polymers and hydrogels are particularly promising due to their super-hydrophilic, antifouling nature^19–21^ and established clinical use in various IMDs^22^. Studies have shown that zwitterionic hydrogels can nearly eliminate the FBR for up to three months in mice, due to their antifouling properties^20^. However, these coatings are electronically inactive, which may hinder the transport of analytes and ions for biosensors and electrophysiological devices. In such applications ^5–7,23^, having conductive or semiconductive materials directly interfacing with tissue is often more desirable to preserve device function.

As electronic materials for IMDs, conjugated polymers are promising candidates due to their low mechanical moduli, broad chemical design space, and lower interfacial impedance^24^. Specifically, poly(3,4-ethylenedioxythiophene):poly(styrene sulfonate) (PEDOT:PSS) has attracted the most attention because it is chemically stable, highly conductive, highly processable, and non-cytotoxic^25,26^. PEDOT:PSS has demonstrated its usefulness in applications ranging from biosensors to electrophysiological recording and stimulation^27–32^. Although PEDOT:PSS has been shown to be biocompatible in terms of cytotoxicity, there is still a lack of understanding of the FBR against PEDOT:PSS and PEDOT derivatives. Most prior studies assessing the FBR to PEDOT:PSS have relied on immunofluorescence (IF) to measure the infiltration of immune cells (e.g., macrophages)^28,32–36^. However, the specific ways that IF was used in these studies may have limitations in giving complete and accurate information about the FBR for the following reasons. First, IF has been used to detect a limited number of markers, leading to an incomplete picture of the FBR. Second, using IF to compare the number of infiltrating immune cells without information about specific phenotypes (e.g., M1 and M2 for macrophage, etc.) does not necessarily reflect FBR severity^37–39^. For example, it has been shown that tissue-resident macrophages, rather than recruited macrophages, are responsible for FBR-related fibrosis^40^.

In fact, negatively charged polymers have been reported to lead to poor FBR outcomes^3,41–44^. As such, it is reasonable to speculate that the negatively charged PSS in PEDOT:PSS follows this observation. In this work, by combining multiple immunological assays, including histology, immunofluorescence, flow cytometry, cytokine analysis, and transcriptomics, we show that PEDOT:PSS is indeed associated with fibrosis (Fig. 1a,c). Therefore, design strategies for improving the intrinsic immunocompatibility of PEDOT:PSS are highly desirable, yet remain a major challenge. To date, the realization of immunocompatible properties in polymers has almost exclusively relied on the design and synthesis of new chemical structures incorporating immune-compatible components^11,12,21,45^, including a recent report on immune-compatible semiconducting polymers^46^. However, because PEDOT:PSS is typically formulated as an aqueous colloidal dispersion, this conventional molecular design strategy—based on synthesis in organic solutions— is not technically feasible. As a result, identifying a fundamentally different approach remains a major challenge.

**Fig. 1.**
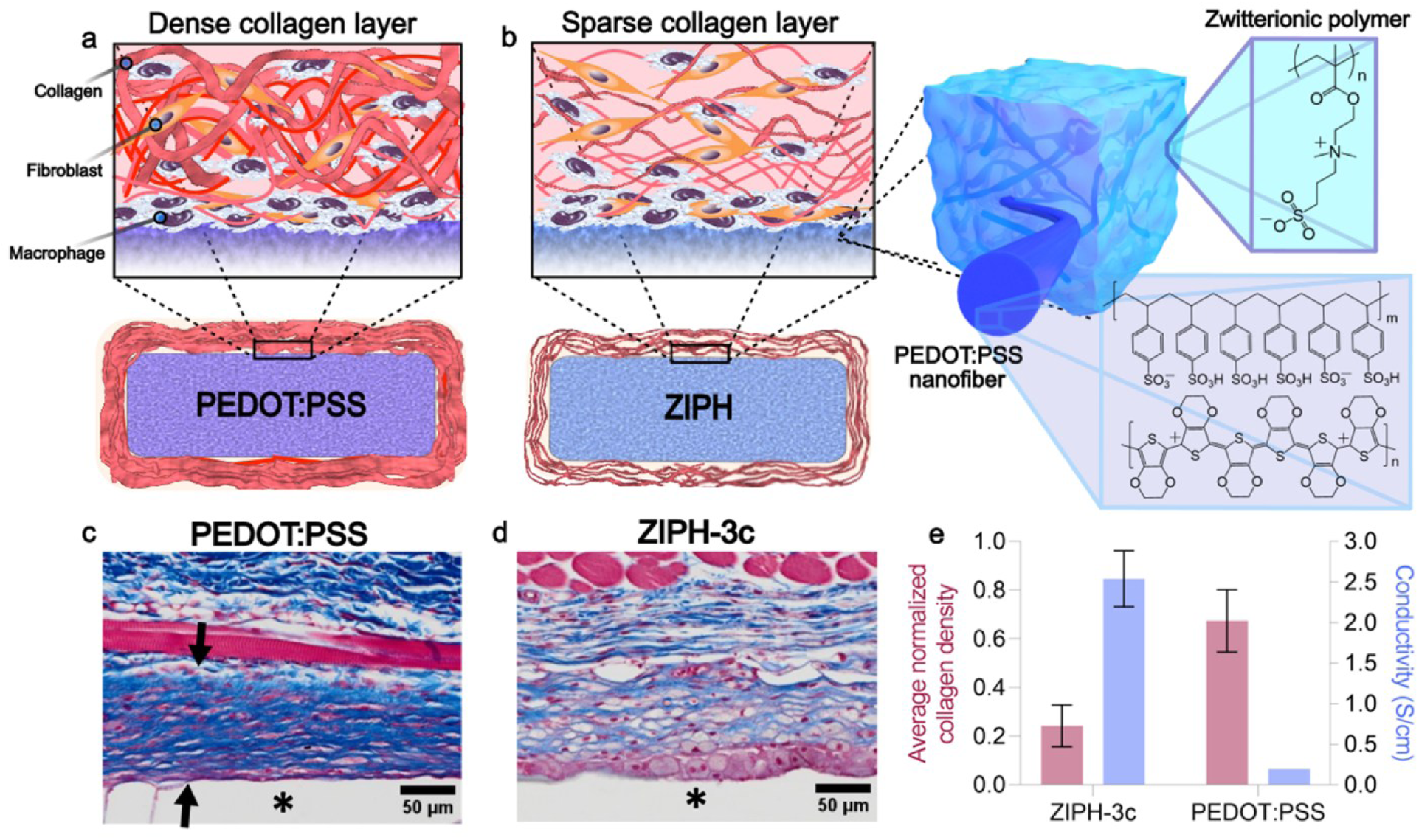
Design of zwitterionic PEDOT:PSS hydrogel (ZIPH) achieves superior immunocompatibility and conductivity compared to PEDOT:PSS. **a,** Schematic illustration of the FBR against PEDOT:PSS. **b,** Schematic illustration of the FBR against ZIPH and its nanoscale morphology, which has much less collagen deposition compared to PEDOT:PSS. **c,** Representative MTS-stained tissue sections of explanted PEDOT:PSS. The collagen layer is marked with arrows, and the locations of implants appear clear and are indicated by asterisks. **d,** Representative MTS-stained tissue sections of explanted ZIPH-3c, which shows a much less dense collagen layer. **e,** Average normalized collagen density and conductivity value of ZIPH-3c compared to PEDOT:PSS.

We hypothesize that creating double-network morphologies between PEDOT:PSS and an immune-compatible polymer, with a sufficiently small phase separation length scale, could represent a viable yet unexplored strategy to impart immune-compatible properties to PEDOT:PSS. To this end, we introduce a design and processing method that promotes PEDOT:PSS to form nanofiber networks embedded in a zwitterionic hydrogel matrix (Fig. 1b). Compared to unmodified PEDOT:PSS hydrogels, our zwitterionic PEDOT:PSS hydrogel (ZIPH) exhibits a reduction of FBR-associated fibrotic severity by 64% (Fig. 1d,e), in addition to an increase in electrical conductivity by more than one order of magnitude (Fig. 1e). Surprisingly, the FBR-associated fibrotic severity is even markedly lower than that of the zwitterionic hydrogel (by 53%) utilized in this design, which could come from certain previously unknown effects from the phase separation morphology. Improved chronic electrophysiological recordings obtained with ZIPH electrodes have further validated the benefits provided by the decreased fibrotic severity for practical applications and demonstrates that such material designs can be stable under harsh *in vivo* conditions. To our knowledge, no studies for implantable biomaterials have explored double-network morphologies as a design strategy for suppressing FBR-mediated fibrosis. As such, the design strategy presented in this work could provide unique insights for other types of biomaterials to achieve suppressed FBR.

### Design of immunocompatible, conductive hydrogels

For combining PEDOT:PSS and zwitterionic polymers for implantable electronics, there are three major criteria that need to be satisfied. First, the double-network morphology should have the zwitterionic matrix dominate the overall immunocompatibility. For this, nanoscale phase separation is preferred so that PEDOT:PSS phases are smaller than the length scales (tens to hundreds of nm) that can be recognized by cells^47^. Second, to achieve high conductivity, the PEDOT:PSS domain needs to be interconnected to facilitate charge transport^48^. Third, to keep stable electrical performance in implantable environments, the design must not have severe swelling in water.

Based on the aforementioned criteria, poly(sulfobetaine methacrylate) (PSB) was chosen as the zwitterionic component. Sulfobetaine-based small molecules have been reported to increase the conductivity of PEDOT:PSS films^49^. Furthermore, due to the strong intra- and inter-chain electrostatic attraction, PSB hydrogels swell the least compared to other zwitterionic hydrogels.^20,50^ We use it to form double-network ZIPH with PEDOT:PSS by firstly dissolving SB monomer, crosslinker (*N,N’-*methylenebisacrylamide), initiator (ammonium persulfate or 2-hydroxy-2-methylpropiophenone), and 4-ethylbenzenesulfonic acid (EBSA) as an additive in the aqueous PEDOT:PSS dispersion, and then polymerizing the mixture in bulk (thermal initiation) or thin-film (UV initiation) states. In particular, the addition of EBSA is hypothesized to facilitate the formation of rod-like structures (Supplementary Fig. 1) for the PEDOT phase, through the destabilization of the micellar structure of PEDOT:PSS.

To further promote the aggregation and interconnectivity of the PEDOT phase in ZIPH, we devised a novel solvent annealing method that involves alternatively exposing ZIPH to ethanol and 2,2,2-trifluoroethanol (TFE) with a certain number of cycles (ZIPH-*n*c for *n* cycles). Because TFE is a good solvent for uncrosslinked PSB while having poor chemical compatibility with PEDOT:PSS, we hypothesize that TFE could penetrate into the PSB hydrogel matrix to further destabilize the PEDOT:PSS phase, thereby promoting the formation of larger PEDOT:PSS aggregates in the PSB hydrogel matrix (Supplementary Fig. 2). Before each TFE treatment, ZIPH is treated with ethanol in each cycle to deswell the hydrogel, thereby preemptively counteracting the excessive swelling of ZIPH by TFE. In addition, the deswelling by ethanol causes the PEDOT:PSS phases to be in closer proximity to each other, which increases the likelihood of adjacent PEDOT:PSS phases to connect (Fig. 2a).

**Fig. 2.**
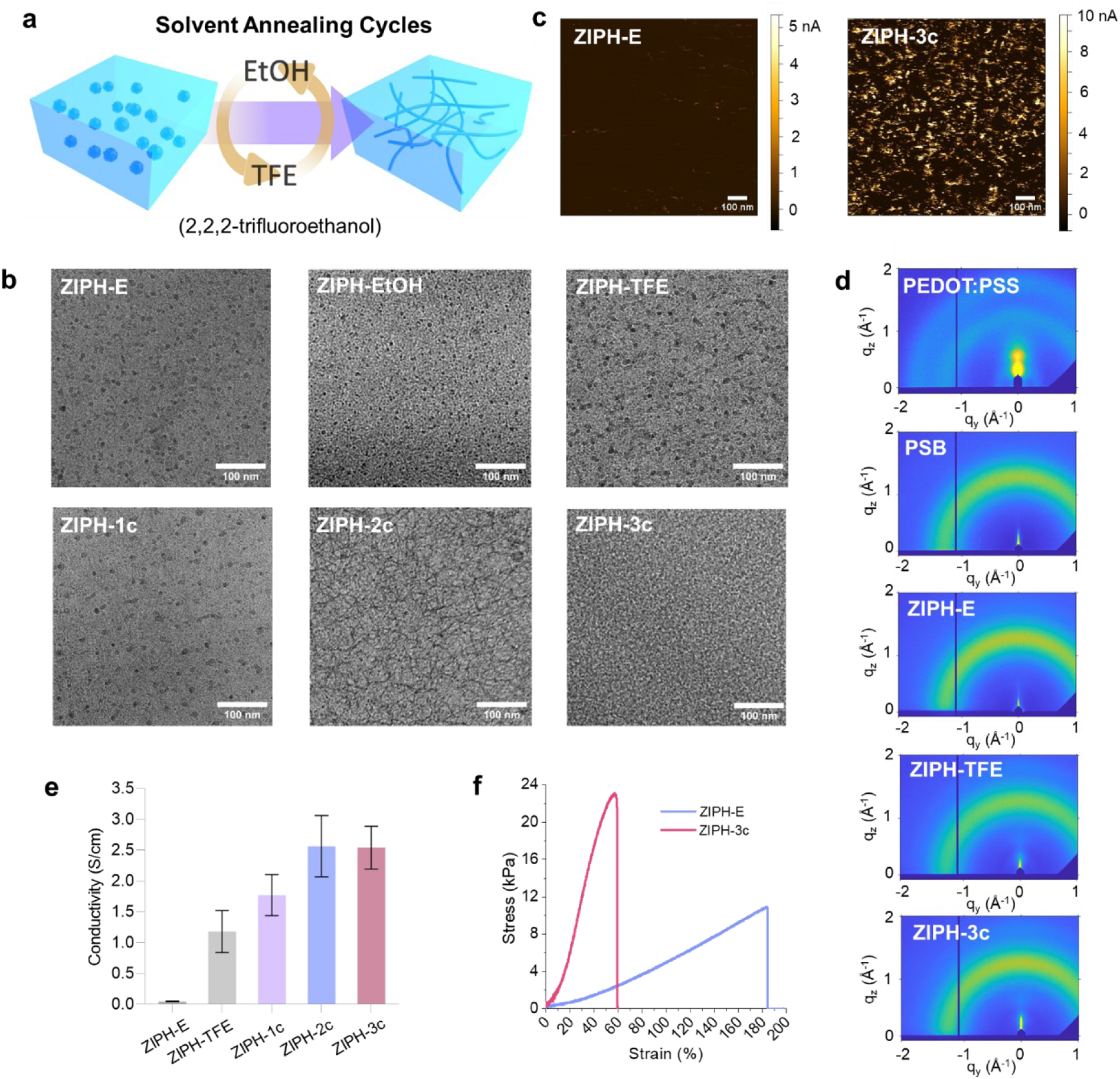
Solvent annealing treatment on ZIPH facilitate the formation of percolated PEDOT:PSS networks and higher conductivity. **a**, Schematic illustration of the morphological changes that occur to ZIPH after ethanol-TFE annealing cycles. **b**, Cryo-TEM images of ZIPH subjected to various solvent annealing conditions. With each ethanol-TFE cycle, the PEDOT:PSS particles transition to a more interconnected fibrillar morphology. **c**, C-AFM images of ZIPH before (ZIPH-E) and after (ZIPH-3c) 3 cycles of ethanol-TFE annealing cycles. **d**, GIXD diffraction patterns of PSB hydrogel and ZIPH subjected to various solvent annealing conditions. **e**, Conductivity values of non-implanted ZIPH samples prepared with various solvent annealing conditions obtained with a standard four-point measurement. All hydrogels were pre-equilibrated in PBS and autoclaved before being measured in a state fully hydrated with PBS. **f**, Tensile curves of ZIPH before (ZIPH-E) and after (ZIPH-3c) 3 cycles of ethanol-TFE annealing cycles.

### Morphology and conductivity evolution of ZIPH during solvent annealing

The hypothesized effect of the solvent annealing process in facilitating the formation of PEDOT nanofibrous morphology is validated by cryogenic transmission electron microscopy (cryo-TEM) on ZIPH samples that went through different solvent annealing processes (Fig. 2b). For ZIPH blended with EBSA for promoting aggregation but without any solvent annealing steps (referred as ZIPH-E), dispersed PEDOT:PSS nanoparticles of 5-15 nm in diameter were observed. This shows that the favorable chemical compatibility between PSB and PEDOT:PSS does not provide enough driving force for PEDOT:PSS to grow nanoparticles into larger aggregates. After the solvent treatments in only ethanol or TFE, or 1 cycle of ethanol-TFE, there is minimal change in the nanoparticle morphology for PEDOT:PSS. After 2 to 3 solvent annealing cycles, the interconnected nanofibers started to emerge. The formation of the PEDOT:PSS fibrous structures from the solvent treatment processes can also be observed using conductive atomic force microscopy (C-AFM), as shown by the comparison of ZIPH-E and ZIPH-3c in Fig. 2c.

Grazing-incidence X-ray diffraction (GIXD) revealed that in contrast to the crystalline structure of pristine PEDOT:PSS, ZIPH suppresses the long-range crystallization of PEDOT (Fig. 2d). The solvent treatment processes have little effect in changing such amorphous structure. The diffraction pattern was nearly identical to that of PSB hydrogel (Fig. 2d). The PSS to PEDOT ratio also remained largely unchanged (Supplementary Discussion, Supplementary Fig. 3).

As a result of the solvent annealing-induced formation of the interconnected nanofibrous morphology of PEDOT:PSS, the conductivity of non-implanted, hydrated ZIPH samples improved substantially from ∼10^−4^ S/cm for ZIPH-E to ∼2.5 S/cm for ZIPH-2c and ZIPH-3c (Fig. 2e and Supplementary Fig. 4a & 5), which is much higher increase than other reported treatment methods utilized on ZIPH samples (Supplementary Fig. 6). For dry, thin film samples, the conductivity of ZIPH-3c reached 80 S/cm (Supplementary Fig. 7 & 8). The effective capacitance of ZIPH-3c was similar to that of PEDOT:PSS (Supplementary Fig. 4b & 9). For TFE-only or 1 cycle of ethanol-TFE annealing, even though little changes were observed in the morphology, increases in conductivity have already started to take effect. The solvent treatment process also led to the increase in Young’s modulus, from 1.4 kPa for ZIPH-E to 9.9 kPa for ZIPH-3c (Fig. 2f and Supplementary Fig. 10). This is also due to the formation of an interconnected network by the more rigid phase of PEDOT:PSS.

### ZIPH-3c is associated with the lowest collagen density among non-drug-eluting implants

Next, we studied the FBR behaviors against ZIPH by subcutaneously implanting disc-shaped samples into mice (Supplementary Fig. 11-13). After pre-determined timepoints, the mice were sacrificed at 1-, 2-, 4-, 8-, 12-, and 24-weeks post-implantation and histological analyses of explanted tissues around the implants were conducted (Fig. 3a & b and Supplementary Fig. 15). Using MTS, the density of the collagen layer formed as a result of the FBR was quantified. Young’s moduli for all implants tested were kept at around 10 kPa except for PEDOT:PSS to account for mechanical effects (Supplementary Discussion, Supplementary Table 1, Supplementary Fig. 14), and the surface roughness for all implants were kept at the nanometer scale by using glass and plastic molds with a surface roughness on the order of a few nanometers to account for topographical effects. As shown by the average collagen density of the fibrotic capsule over time (Fig. 3c & d and Supplementary Fig. 16 & 17), ZIPH-3c indeed had substantially decreased collagen density than the PEDOT:PSS hydrogel at all timepoints, which validated that the incorporation of the zwitterionic hydrogel PSB effectively suppresses the FBR. Surprisingly, ZIPH-3c had a much lower collagen density than the PSB hydrogel at all timepoints as well (Fig. 3b & c and Supplementary Fig. 16). For the thickness of the fibrotic capsule, ZIPH-3c was associated with the second thickest, behind only PEDOT:PSS, at most timepoints (Supplementary Fig. 18). This indicates that the double-network morphology might be having some unique effect in helping to decrease the collagen density of FBR-associated fibrosis without necessarily decreasing the thickness. Along this line, the higher collagen density of ZIPH without the addition of EBSA or solvent treatment (ZIPH-u) compared to ZIPH-3c implies that the interconnected nanofibrous morphology of PEDOT:PSS in ZIPH-3c is also favorable for suppressing the FBR (Supplementary Fig. 19). To the best of our knowledge, such effects of double-network morphology on the FBR have never been reported before in other biomaterials studies.

**Fig. 3.**
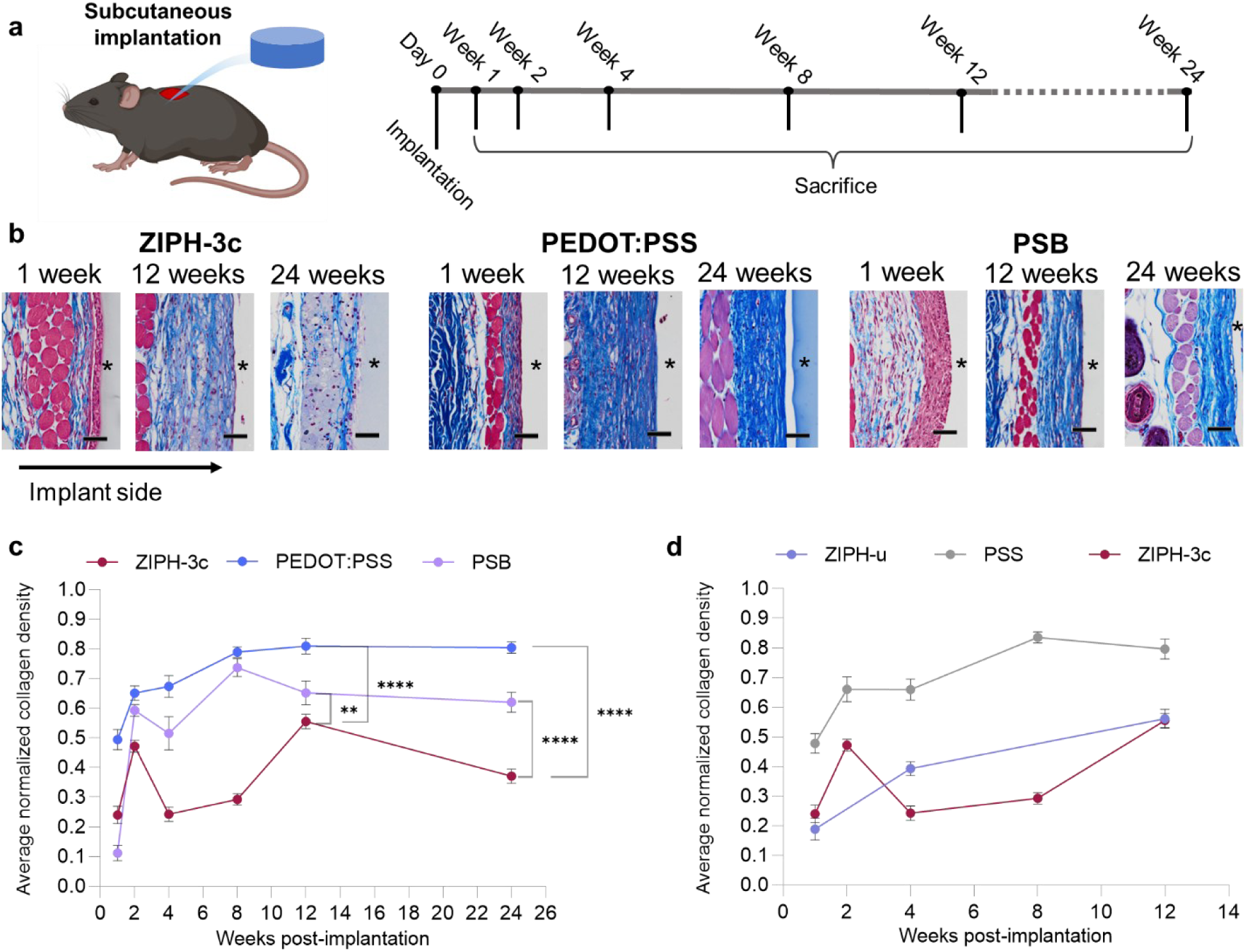
Measurement of FBR-induced collagen deposition shows that ZIPH-3c has one of the lowest collagen density. **a**, Schematic illustration of the subcutaneous implantation timeline. **b**, Representative MTS-stained tissues from 3 types of implants 1-, 4-, 12-, and 24-weeks post-implantation. Locations of implants appear clear and are indicated by asterisks. **c-d**, Average collagen density (blue pixel density) of the fibrotic capsule over time, determined from MTS-stained tissue. Scale bars, 50 μm. Error bars, mean ± s.d. N=5-6 mice per treatment. One-way ANOVA was used to compare collagen densities between treatments. **, P < 0.001; ****, P < 0.0001.

For PEDOT:PSS hydrogel, the high FBR-induced fibrosis could largely come from the negatively charged PSS^3,41–44^. To test this, we also prepared and tested PSS hydrogels, which, indeed, gave the highest collagen density at the surface of the implant (Fig. 3d and Supplementary Fig. 17).

Although the immunocompatible properties of zwitterionic polymers mostly come from the antifouling properties, protein adsorption assays indicate that this may not be the case for the ZIPH designs. Among all the tested samples (Supplementary Figs. 20-22), both ZIPH-3c and ZIPH-u have some of the highest levels of protein adsorption. This suggests that the embedment of PEDOT:PSS in the PSB matrix may not be enough to negate the high protein adsorptive behavior of PEDOT:PSS. Our result also shows that PSS is indeed responsible for the high protein adsorptive behavior of PEDOT:PSS (Supplementary Fig. 20-22), which is consistent with the *in vivo* collagen density results. Overall, the trend from these protein adsorption results does not fully agree with the *in vivo* histology results discussed above, which indicates that in our designs, protein adsorption only has a loose correlation with the fibrosis associated with each implant type.

### Immune response for ZIPH-3c diverges from its parent materials

In the FBR, macrophages are known to play a central role in the final outcome^2,3,37^. To gain a deeper understanding of the different immune responses induced by different hydrogel designs, we used both flow cytometry (FC) and immunofluorescence (IF) to analyze the population size and phenotypes of macrophages surrounding each implant at 1-, 4-, and 12-weeks post-implantation (Supplementary Fig. 23-27). FC data (Fig. 4a and Supplementary Fig. 23-25) indicated that ZIPH-3c was associated with the greatest number of Ly-6C^-/lo^ F4/80^hi^ macrophages compared to other samples including PEDOT:PSS, PSB, PSS, and sham for all timepoints. Furthermore, the polarization states of Ly-6C^-/lo^ F4/80^hi^ macrophages, M1 and M2 macrophages, were analyzed. M1 and M2 macrophages are thought to be pro-and anti-inflammatory, respectively, but studies have shown that an imbalance in macrophage polarization in either direction compared to sham can be a primary driver for poor FBR and fibrosis outcomes^37,51^. Macrophage polarization states exist on a continuum, and the M1/M2 classification can vary significantly depending on the markers used^37,51^. To gain a clear understanding on the average polarization states of macrophages, iNOS (M1) / Arg-1 (M2) and CD86 (M1) / CD206 (M2) single positive ratios were quantified using FC at 4 weeks. Both macrophage polarization classification methods indicated that the immunological milieu of ZIPH-3c was biased towards M1 polarization (Fig. 4b & c and Supplementary Fig. 24 & 25). Immunofluorescence (IF) microscopy confirmed that the Ly-6C^-/lo^ F4/80^hi^ macrophages analyzed using FC were localized at the surface of the implants (Fig. 4d and Supplementary Fig. 28-31). Since an M1 bias suggests a polarization to type 1 immunity, T cell phenotypes were also assessed, and the fraction of cells expressing T-bet, a master regulator of Type 1 T helper cells (T_h_1) and a marker for type 1 immunity, among CD3^+^ T cells was found to be the highest for ZIPH-3c (Fig. 4e), consistent with the observed M1 bias. To check if markers for type 1 immunity necessarily imply an inflammatory response, FoxP3^+^ T regulatory cells (T_reg_s) were also quantified. It was found that ZIPH-3c was associated with the highest fraction of T_reg_s among CD4^+^ T helper cells (T_h_ cells), indicating the co-existence of an anti-inflammatory regulatory response (Supplementary Discussion). Consistent with previous results for zwitterionic hydrogels^21^, PSB exhibited a preference towards M2 polarization and type 2 immunity (Fig. 4b,c,e and Supplementary Fig. 24), an indication that the immune response to ZIPH-3c has significantly diverged from PSB.

**Fig. 4.**
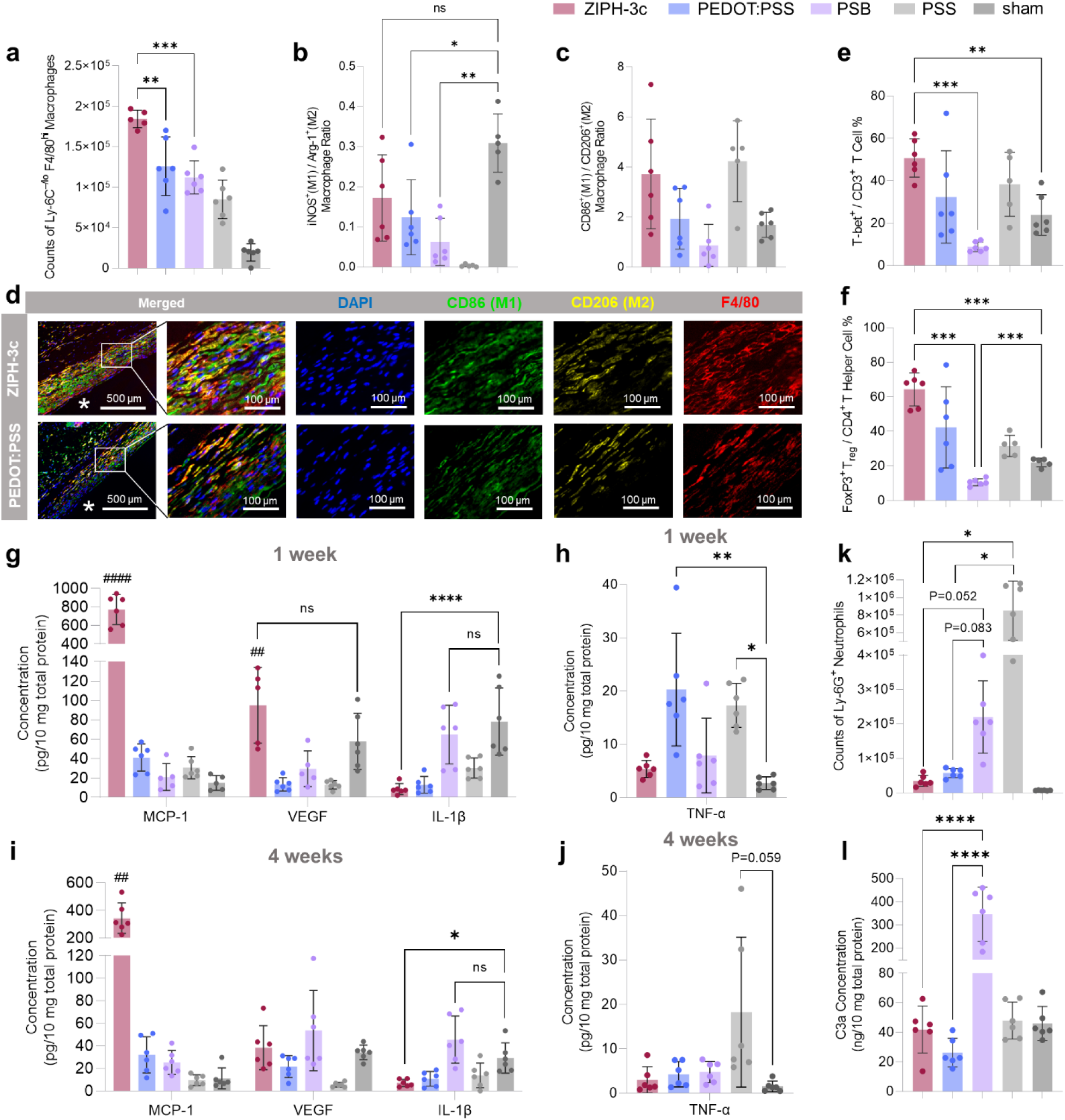
Comparisons of key immune cell populations and cytokine concentration from different implants show that ZIPH-3c induces a distinct immune response compared to its parent materials. **a**, Total counts of macrophages associated with each type of implant, determined by flow cytometry and 4-week post-implantation. Macrophages were defined as F4/80^+^ cells. ZIPH-3c is consistently associated with the greatest number of macrophages. **b**, iNOS^+^ Arg-1^‒^ / iNOS^‒^ Arg-1^+^ macrophage ratios associated with each implant, determined by flow cytometry at 4 weeks post-implantation. iNOS^+^ Arg-1^‒^ macrophages are considered to be M1-like and iNOS^‒^ Arg-1^+^ macrophages are considered to be M2-like. ZIPH-3c is associated with the most M1-like macrophages. **c**, CD86^+^ CD206^‒^ / CD86^‒^ CD206^+^ macrophage ratios associated with each implant, determined by flow cytometry at 4 weeks post-implantation. CD86^+^ CD206^‒^ macrophages are considered to be M1-like and CD86^‒^ CD206^+^ macrophages are considered to be M2-like. ZIPH-3c is associated with the most M1-like macrophages. **d**, Representative immunofluorescence microscopy images of tissues stained with F4/80 (macrophage), CD86 (M1), and CD206 (M2). Locations of implants appear clear and are marked by asterisks. **e**, Percentage of CD3^+^ T cells that are T-bet^+^. T cells associated with ZIPH-3c have a high fraction of T-bet^+^ cells. **f**, Percentage of CD4^+^ T helper cells that are FoxP3^+^ T_reg_s. CD4+ T helper cells associated with ZIPH-3c have a high fraction of FoxP3^+^ T_reg_s. **g-h**, Key cytokines associated with each implant, determined via LegendPlex assay at 1-week (g-h) and 4-week (i-j) post-implantation. MCP-1 is consistently upregulated for both ZIPH-3c and ZIPH-u. **k**, Total counts of neutrophils associated with each implant, determined by flow cytometry 1-week post-implantation. Neutrophils were defined as Ly-6G^+^ cells. **l**, Complement C3a concentration associated with each implant, determined by ELISA 1-week post-implantation. The high number of neutrophils is correlated with the generation of C3a. PSB has a distinctly higher neutrophil number and C3a concentration than the rest of the hydrogels that contain PSS. Error bars, mean ± s.d. N = 5-6 mice per treatment. One-way ANOVA was used for statistical comparison of multiple means. *, P < 0.05; **, P < 0.01; ****, P < 0.0001; ##, P < 0.01 compared to rest; ####, P < 0.0001 compared to rest.

Cytokines secreted during the FBR also provide in-depth information on the different immune responses. We used a 14-cytokine LegendPlex panel to quantify cytokine concentrations at 1-, 2-, 4-, 8-, and 12-weeks post-implantation (Fig. 4g-j and Supplementary Fig. 32-36). IFN-γ, a crucial cytokine for type 1 immunity, was found to be expressed at similar levels for all implants, including ZIPH-3c, suggesting that T_reg_s may be counteracting the inflammatory effects of type 1 immunity (Supplementary Fig. 32-36). However, IL-10, a T_reg_-associated cytokine, was not significantly upregulated (Supplementary Fig. 32-36), suggesting an IL-10-independent regulatory mechanism (Supplementary Discussion). In addition, other cytokine expression profiles of ZIPH-3c and ZIPH-u are found to significantly diverge from their parent materials, PEDOT:PSS and PSB. In particular, ZIPH-3c and ZIPH-u were found to be associated with significantly higher concentrations of MCP-1 at all timepoints tested and higher concentrations of VEGF at the 1-, 2-, and 4-week timepoints compared to the rest of the implants (Fig. 4g & i and Supplementary Fig. 32-36). The significant upregulation of MCP-1 is a potential explanation for why ZIPH-3c is associated with the greatest number of macrophages (Supplementary Discussion). Interestingly, even though MCP-1 and VEGF are known to be powerful arteriogenic and angiogenic cytokines, respectively^52,53^, vascularization in the surrounding areas did not match the high levels of MCP-1 and VEGF at 4-weeks post-implantation (Supplementary Fig. 37). Another important cytokine is IL-1β, which is one of the first cytokines released during tissue injury^54^. All the hydrogels except for PSB had suppressed IL-1β levels, and ZIPH-3c had one of the lowest IL-1β levels for all time points (Fig. 4g & i and Supplementary Fig. 38a). This difference between ZIPH-3c and PSB may explain the lower fibrotic tendency of ZIPH-3c. For PEDOT:PSS and PSS hydrogels, significant upregulation of TNF-α was found after 1 week, suggesting an inflammatory response (Fig. 4h). Such upregulation persisted at 4 weeks for PSS (Fig. 4j and Supplementary Fig. 32-34). Because TNF-α is a powerful pro-inflammatory cytokine leading to fibrosis, this partially explains the fibrotic response against PEDOT:PSS and PSS. Cytokine concentration evolved significantly over time, and some key cytokines were observed to be in sync with the evolution of collagen density and thickness over time (Fig. 3c and Supplementary Fig. 38b).

Another important type of cell in the FBR is neutrophils, which act as the first responders to the implants. We used flow cytometry to quantify Ly-6G^+^ neutrophils. At 1-week post-implantation, the number of Ly-6G^+^ neutrophils associated with PSB and PSS was found to be greater compared to rest (Fig. 4k and Supplementary Fig. 23, 39, 40). It is known that the recruitment of neutrophils can be facilitated by the activation of the complement system^3^. Therefore, we further compared complement activation of different samples by quantifying protein C3a, which is generated as a result of complement activation. At the 1-week timepoint, PSB had substantially higher C3a concentrations in the peri-implant tissue than the rest (Fig. 4l and Supplementary Fig. 41), suggesting that PSB is activating the complement cascade. This finding is consistent with a previous study showing that a zwitterionic polymer, poly(2-metharyloyloxyethyl phosphorylcholine), facilitates complement activation^55^. For PSS, neutrophil recruitment may primarily be facilitated by TNF-α. For the ZIPH samples, despite having PSB as a significant fraction, the lower trafficking of neutrophils and complement activation (Fig 4k & l and Supplementary Figs. 23 & 41) could come from PSS, which has been shown to suppress complement activation^56,57^. This difference between ZIPH and PSB could also be part of the reason for the lower collagen density from ZIPH.

### ZIPH-3c is associated with a unique gene expression pattern that suppresses fibrosis

To gain further insight into the mechanisms of the FBR against ZIPH-3c, the tissues associated with the implant materials were analyzed with a 50-gene NanoString panel 4-and 12-weeks post-implantation (Supplementary Fig. 42-46). The most difference in gene expression occurred at 4-weeks post-implantation. Principal component analysis (PCA) was used to analyze the data. When the data are plotted on a vector space defined by the first (PC1) and third (PC3) principal components, there is a loose separation between the mildly fibrotic implants, consisting of ZIPH-3c and ZIPH-u, and the more severe fibrotic implants, consisting of PEDOT:PSS and PSB (Fig. 5a).

**Fig. 5.**
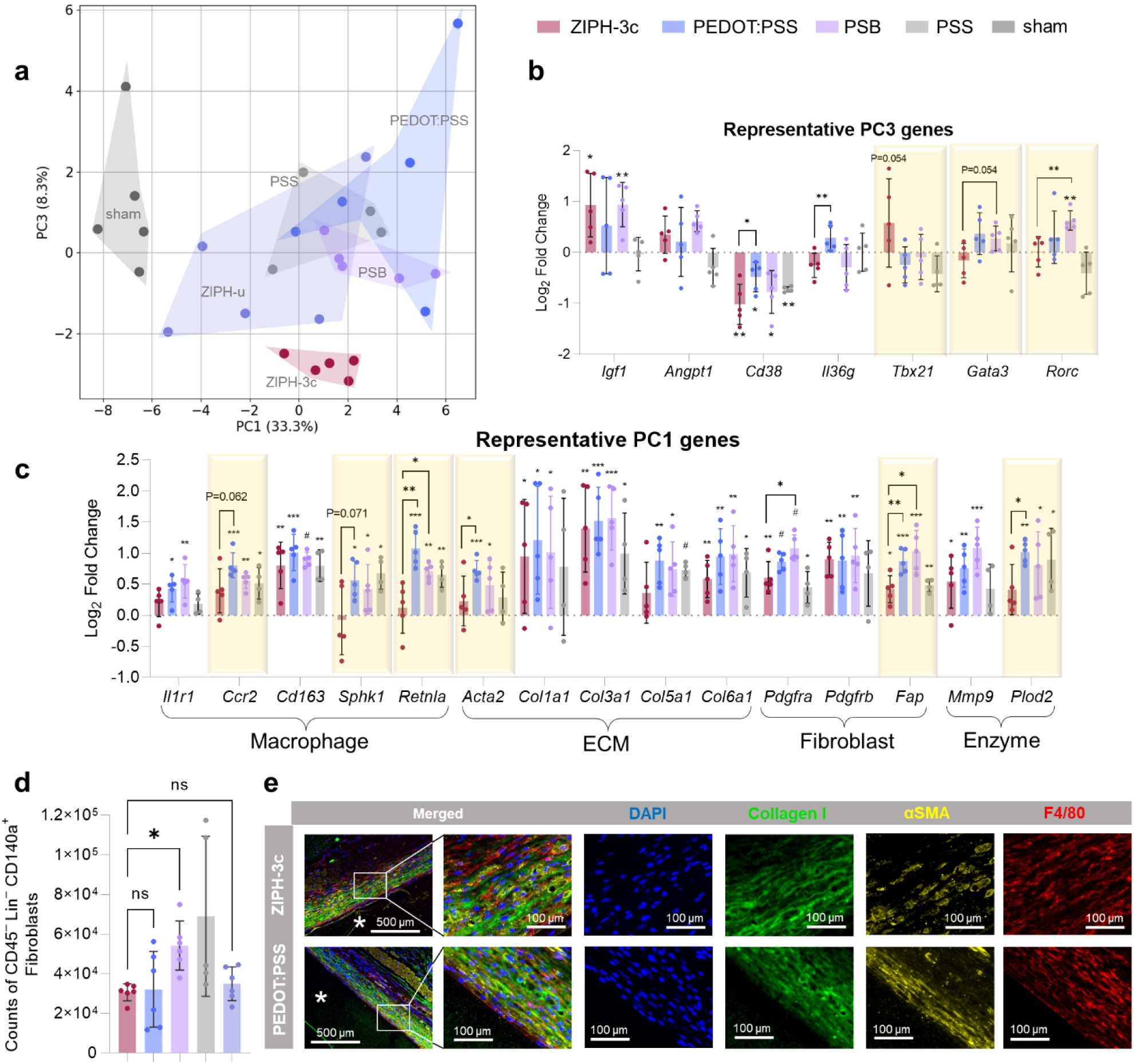
Gene expression analysis coupled with characterization of fibroblast and myofibroblast populations show that ZIPH-3c is associated with a unique, mildly fibrotic gene expression pattern. **a,** Principal component analysis (PCA) of a 50-gene panel from 4-week post-implantation samples plotted against each other. **b,** Genes associated with the negative direction of the third principal component (PC3). The expression of *Tbx21* (T-bet) makes ZIPH-3c unique among the samples in the experiment. T helper cell-related genes are highlighted. **c,** Differentially expressed genes associated with the positive direction of the first principal component (PC1). All the genes have some connection to fibrotic diseases. Genes that are statistically significantly different from sham control and have a strong connection with fibrosis are highlighted. **d,** Total counts of fibroblasts associated with each implant, determined by flow cytometry 4 weeks post-implantation. Fibroblasts are defined as CD45^‒^ Lin^‒^ CD140a^+^, in which Lin consists of CD31, Ep-CAM, Tie-2, and Ter-119. **e,** Representative immunofluorescence microscopy images of tissues stained with collagen I, αSMA (myofibroblast), and F4/80 (macrophage). Locations of implants are marked by asterisks. Error bars, mean ± s.d. N=5-6 mice per treatment. One-way ANOVA was used for statistical comparison of multiple means. *, P < 0.05; **, P < 0.01; ***, P < 0.001; #, P < 0.00001.

Compared to the other implants, the gene expression pattern for ZIPH-3c is unique. ZIPH-3c forms a tight, unique cluster on the PC1-PC3 vector space and is one of the main drivers of variance for PC3 (Fig. 5a). Among all the representative PC3 genes, the gene with the most distinct expression level for ZIPH-3c is *Tbx21*, a gene for T-bet. Although not statistically significant (P = 0.054), *Tbx21* was upregulated for ZIPH-3c, consistent with FC results. T-bet is the master regulator for Type 1 T helper cells (T_h_1), which, under certain circumstances, is implicated in the suppression of fibrosis^58–60^. But for T_h_2 and T_h_17, which are implicated in the induction^61–63^, their master regulators, *Gata3* (T_h_2) and *Rorc* (T_h_17), from ZIPH-3c were found to be indistinguishable from the baseline sham control (Fig. 5b and Supplementary Fig. 44b). On the other hand, PC1 is composed of genes that have the biggest correlation with fibrosis^64–68^, including *Ccr2*, *Sphk1*, *Retnla, Acta2*, *Fap,* and *Plod2*. Strong upregulations of these fibrotic genes are consistently observed for PEDOT:PSS, PSB, and ZI-SAM-PH, but not for ZIPH-3c or ZIPH-u (Fig. 5c and Supplementary Fig. 44a). Not surprisingly, collagen genes also contribute strongly to PC1. Their expression patterns overall follow the trend of collagen deposition on different implants (Fig. 5c and Supplementary Fig. 44a). However, given that the differences between implants are not statistically significant, there should be other genes and pathways important for the FBR outcome.

Fibroblasts and myofibroblasts are responsible for collagen deposition, with the increasing presence of myofibroblasts associated with increasing FBR severity^69^. *Acta2*, a gene expressed by myofibroblasts, was found to be upregulated with statistical significance for PEDOT:PSS and PSB implants, while ZIPH-3c maintained a similar level with sham (Fig. 5c). This agrees with IF imaging using αSMA (*Acta2*), which shows concentrated myofibroblasts at the surface of PEDOT:PSS but not for ZIPH-3c (Fig. 5e). In addition, CD45^‒^ Lin^‒^ CD140a^+^ cells, which encompass both fibroblasts and myofibroblasts, were quantified via FC. ZIPH-3c, was associated with the one of the least number of CD45^‒^ Lin^‒^ CD140a^+^ cells across all timepoints (Fig. 5d and Supplementary Fig. 23-25). IF imaging also confirmed that collagen I, αSMA, and F4/80 were all co-localized at the implant surface, confirming that the fibrotic response was directed against the implants and both F4/80^+^ macrophages and CD45^‒^ Lin^‒^ CD140a^+^ cells are part of the same immunological milieu (Fig. 5e and Supplementary Fig. 47-49). In contrast to the FC result for CD45^‒^ Lin^‒^ CD140a^+^ cells, the αSMA expression on ZIPH-3c indicates a low number of myofibroblasts, which is consistent with the current understanding of the FBR^69^.

### ZIPH-3c benefits electrophysiological recordings by decreasing signal loss over time

To demonstrate that the lower collagen density of the fibrotic capsule associated with ZIPH-3c benefits electrophysiological recordings, we further utilized ZIPH-3c to fabricate passive electrocardiography (ECG) recording devices (Fig. 6a, Supplementary Fig. 50, Supplementary Discussion) and implanted them in the subcutaneous space of mice over the course of 12 weeks. Modified vascular access buttons were installed transdermally in mice and connected with two separate ECG electrodes, and the external ports were used to connect to peripheral electronics and to enable chronic recording (Fig. 6b and Supplementary Fig. 51). ECG recordings from ZIPH-3c electrodes were compared with those from pristine PEDOT:PSS electrodes, as well as the PEDOT:PSS electrodes (PEDOT:PSS-Dex) incorporating a dexamethasone elution layer as the gold standard for suppressing FBR (Fig. 6c, d). Because of the lower collagen density associated with ZIPH-3c, we expect it to have less signal decay compared to PEDOT:PSS, which is associated with a dense collagen layer. For PEDOT:PSS-Dex, the controlled release of dexamethasone from the Dex-PDMS layer should eliminate the growth of a fibrotic capsule, also leading to less signal decay (Fig. 6c). These are indeed supported by the statistical results shown in Fig. 6e, with 3-4 device replicates in each group. As expected, both ZIPH-3c and PEDOT:PSS-Dex retained more of the initial signal than PEDOT:PSS by week 4. By week 12, the signal retentions of ZIPH-3c (66%) and PEDOT:PSS-Dex (68%) were twice that of PEDOT:PSS (33%) (Fig. 6e and Supplementary Discussion). These results show that ZIPH-3c can decrease FBR-induced electrophysiological signal decay just as well as dexamethasone.

**Figure 6.**
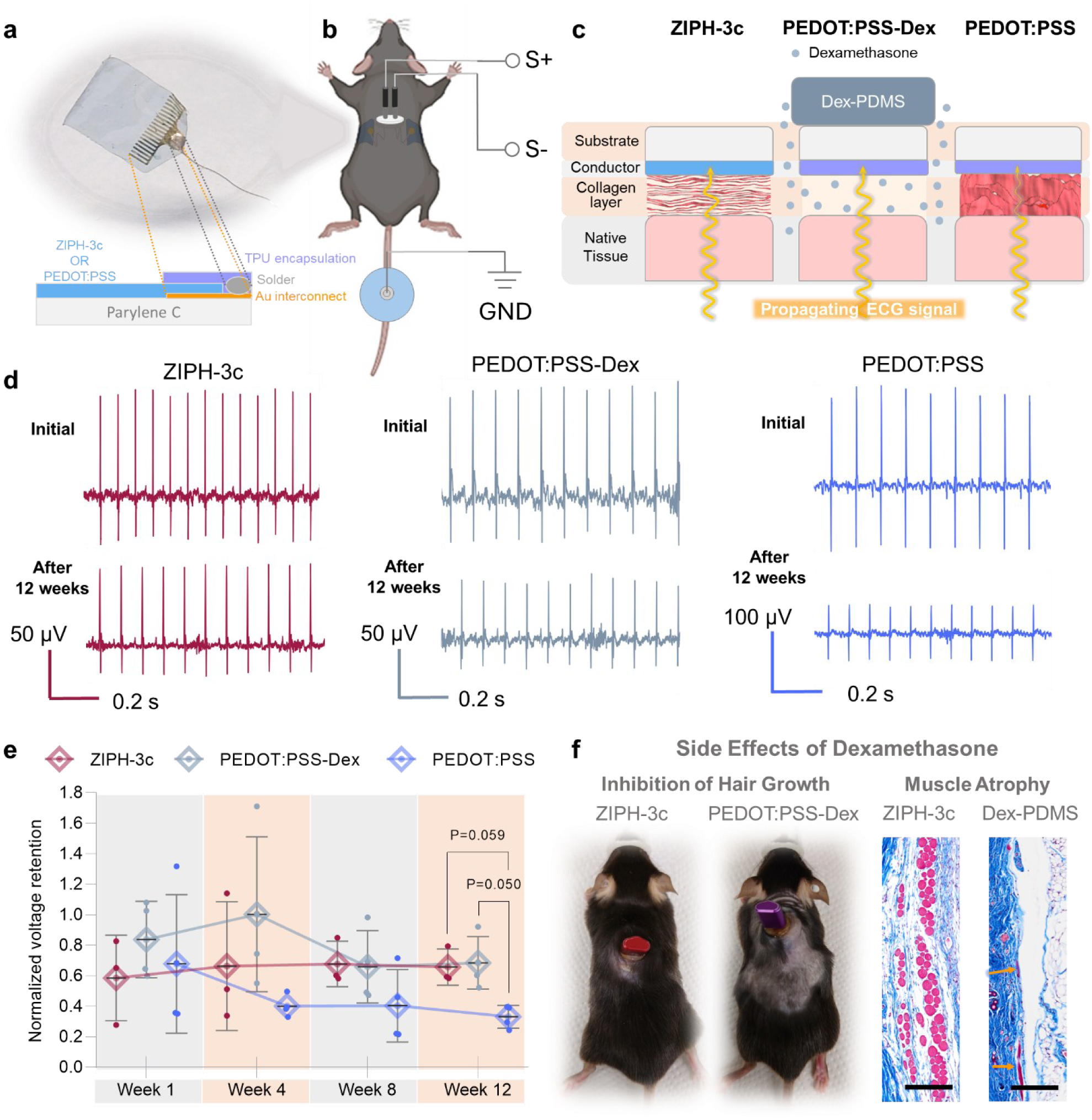
ZIPH-3c electrocardiographic electrodes experience less signal loss compared to PEDOT:PSS electrodes. **a**, Device structure of ECG electrodes used to test the effect of the FBR on electrophysiological recordings. **b**, Electrical connection on anesthetized mouse for ECG recording. Created with BioRender.com. **c**, Schematic illustration of the effects of the fibrotic capsule on ECG signal propagation for different electrode types. **d**, Examples of ECG traces immediately after surgery (top) and 12-weeks post-implantation (bottom). Schematic illustration of the effects of the fibrotic capsule on ECG signal propagation. **e**, Peak R-wave voltages at various timepoints normalized to peak R-wave voltages immediately after surgery. **f**, Side effects of dexamethasone 12-weeks post-implantation, as demonstrated by photographs showing the inhibition of hair growth in mice implanted with PEDOT:PSS-Dex electrodes (left) and MTS-stained tissues showing the atrophy of the panniculus carnosus in mice implanted with Dex-PDMS discs (right). Orange arrows indicate the locations of the remaining panniculus carnosus for the Dex-PDMS-implanted mouse. Scale bar, 200 µm. Error bars, mean ± s.d. N = 3-4 mice per treatment. One-way ANOVA was used for statistical comparison of multiple means.

Notably, the mice with PEDOT:PSS-Dex electrode implants did not grow back the fur shaved for surgery even after 12 weeks, as opposed to the mice implanted with the other electrodes pairs, which mostly grew their fur back after 12 weeks (Fig. 6f and Supplementary Fig. 52). In combination with the muscle atrophy caused by dexamethasone (Fig. 6f and Supplementary Fig. 53), a side effect that has been reported previously^15,16^, this shows that ZIPH-3c electrodes can perform just as well as dexamethasone-eluting electrodes for chronic, electrophysiological implantation without causing the side effects associated with dexamethasone.

## Discussions

To suppress the FBR against PEDOT:PSS by imparting intrinsic immunocompatibility, we report a double-network design of *in situ* formed PEDOT:PSS nanofibers embedded in a zwitterionic hydrogel matrix. By tuning the phase separation morphology, this design not only achieved higher conductivity, but also substantially reduced the FBR by 64% (as measured by the collagen density) compared to pristine PEDOT:PSS hydrogels (Supplementary Table 2). Most interestingly, its FBR severity is markedly lower than its parent zwitterionic polymer, PSB, which suggests some unique synergistic effects between PSB and PEDOT:PSS.

Along with these functional and morphological findings, our systematic immunological experiments combining histology, immunofluorescence, flow cytometry, RNA profiling, cytokine assays, and in vitro protein adsorption experiments have provided deeper insights. First, the poor immunocompatibility of PEDOT:PSS was found to be mainly attributed to the high protein adsorptive properties of negatively charged PSS. Second, the higher immunocompatibility of ZIPH-3c compared to PSB may arise from four possible underlying reasons: a bias towards type 1 immunity while maintaining a high T_reg_ population, inhibition of complement activation by PSS, and suppressed fibrotic gene expression levels. These effects could come from the interplay between a unique combination of polymers and the resulting morphological structure.

Overall, this work suggests that chemical heterogeneity, which has been rarely explored, could be another design parameter for enabling facile “mix-and-match” approaches for imparting immunocompatibility to functional materials without changing the chemical structure. Also, for deepening the immunological understanding of the FBR mechanism, creating model systems with controlled chemical heterogeneity could be a useful tool. For the future uses of PEDOT:PSS in electrically biointerfaced devices, as exemplified by our chronic, in-vivo ECG tests, this work provides the first thoroughly tested solution for addressing the most challenging problem of the foreign-body response.

## Methods

### Materials

PEDOT:PSS (PH1000) was purchased from Clevios and was filtered through a 0.7 μm glass fiber filter (Tish Scientific) before use. The following chemicals were purchased from Sigma-Aldrich: [2-(Methacryloyloxy)ethyl]dimethyl-(3-sulfopropyl)ammonium hydroxide (SBMA, 95%), *N*,*N*′-Methylenebisacrylamide (MBAA, ≥98%), ammonium persulfate (APS, ≥98%), *N,N,N’,N’*-tetramethylethylenediamine (TEMED, 99%), 2-hydroxy-2-methylpropiophenone (HMPP, 97%), 4-ethylbenzenesulfonic acid (EBSA, 95%), sodium styrene sulfonate (NaSS, Sigma-Aldrich, ≥90%), divinylbenzene (DVB, 80%), (3-aminopropyl)triethoxysilane (APTES, 99%), acrylic acid (AAc, 99%), 4,4′-azobis(4-cyanovaleric acid) (ACVA, ≥98%), (vinylbenzyl)trimethyl ammonium chloride (VBTMAC, 99%), sodium dodecylbenzene sulfonate (SDS, ≥98.5%), 4-cyano-4-(phenylcarbonothioylthio) pentanoic acid (CPhPA), and 2-cyano-2-propyl benzodithioate (CPBD, >97%). 2,2,2-trifluoroethanol (TFE, ≥99%) was purchased from Sigma-Aldrich and Tokyo Chemical Industries. 2,2′-azobis[2-(2-imidazolin-2-yl)propane]dihydrochloride (VA-044) was purchased from Wako Chemicals (USA). Poly(allylamine) (PAH, 15% aq. soln.) was purchased from Polysciences. PDMS Sylgard 184 was purchased from Dow-Corning. The acetate buffer was prepared with 0.1 M acetic acid (glacial, Sigma-Aldrich, ≥99.85%), 0.1 M sodium acetate trihydrate (Sigma-Aldrich, >99%), and 0.5 M NaCl at pH 5.2. Hydromed D4 Hydrophilic Thermoplastic Polyurethane (HPU) was purchased from AdvanSource Biomaterials. Elastollan 1185A10 Polyether Type Polyurethane (TPU) was purchased from BASF

### Preparation of hydrogels

For preparing ZIPH-E hydrogels, 250 mg of [2- (Methacryloyloxy)ethyl]dimethyl-(3-sulfopropyl)ammonium hydroxide (SBMA, Sigma-Aldrich, 95%) and 1.4 mg (1 mol%) of *N*,*N*′-Methylenebisacrylamide (MBAA, Sigma-Aldrich, ≥98%) were dissolved in 1.0 mL of PEDOT:PSS dispersion. Then, for the preparation of thermally initiated hydrogels, 10.21 μL (0.1 mol%) of APS (20 mg/mL) and 10.41 μL (0.2 mol%) of TEMED (20 mg/mL) aqueous solutions were added dropwise to the PEDOT:PSS mixture under continuous agitation. For the preparation of UV initiated hydrogels, 7.5 μL (0.1 mol%) of HMPP (20 mg/mL) aqueous solution was added. This was followed by the dropwise addition of 62.5 μL of EBSA aqueous solution (20 wt%) while the mixture is continuously agitated and vortexed for an additional minute. For thermal polymerization, the final mixture was placed under vacuum for 15 minutes, injected into a sandwich sheet mold, and placed in an 80°C oven for 1 hour. For UV polymerization, the final mixture was sparged with nitrogen for 10 minutes, spin coated and exposed to UV irradiation (365 nm wavelength, 4 mW/cm^2^) for 5 minutes. To prepare ZIPH-u hydrogels, the above procedures were carried out without the addition of EBSA. To prepare annealing cycle treated bulk ZIPH hydrogels, ZIPH-E hydrogels were briefly rinsed in ethanol. Subsequently, the hydrogels were immersed in a volume of ethanol three times the volume of the ZIPH sample. Thereafter, the ethanol was discarded, and the hydrogels were immersed in a volume of TFE three times the volume of the ZIPH sample. The TFE was discarded, and the hydrogels were briefly rinsed in water. The whole annealing process was repeated two more times to obtain ZIPH-3c. To prepare annealing cycle treated thin film ZIPH hydrogels, the films were immersed in ethanol for 1 minute and immersed in TFE for 1 minute.

To prepare PSB hydrogels, 1.5 g of SBMA (55 wt%), 4.2 mg of MBAA (0.5 mol%), and 0.882 mg of HMPP (0.1 mol%) were dissolved in PBS. The solution was vortexed for 1 minute and sparged with nitrogen for 10 minutes. The sparged solution was injected into a sandwich sheet mold and irradiated with UV light (365 nm wavelength, 4 mW/cm^2^) for 30 minutes per side.

To prepare PSS hydrogels, 700 mg of NaSS (28 wt%) was dissolved in 1694 μL of DMSO by stirring overnight. To the NaSS solution, 22.6 μL of DVB (4 mol%), 43.4 μL (0.1 mol%) of APS DMSO solution (20 mg/mL), and 44.2 μL (0.2 mol%) of TEMED DMSO solution (20 mg/mL) were added and vortexed for 1 minute. Thereafter, the solution was placed under vacuum for 15 minutes, injected into a sandwich sheet mold, and placed in an 80°C oven for 3 hours. The resulting PSS gel was rinsed twice in PBS.

For the preparation of PEDOT:PSS bulk hydrogels, 16% v/v of DMSO was added to PEDOT:PSS. Then, 10 mg/mL of EMIM OTf was added dropwise while the solution was continuously agitated. The resultant mixture was vortexed for 1 minute, placed under vacuum for 15 minutes, and injected into a sandwich sheet mold with silicone lining on top. The mold was placed on a hotplate, heated to 90 °C, for 90 minutes with the silicone lining side up. Then, the top piece with the silicone lining was removed, and the PEDOT:PSS gel was exposed to air. The PEDOT:PSS gel was dried at 40 °C for 16 hours, followed by another round of drying at 50 °C for 16 hours, at which point the thickness and lateral dimensions of the gel was approximately a third and a half of the original dimensions, respectively. The resulting PEDOT:PSS gel was rinsed twice in PBS.

All gels were washed in PBS for three days with changes of fresh PBS every 12 hours. All hydrogels used in *in vivo* experiments were autoclaved at 121 °C for 30 minutes in PBS. To ensure that most of the EMIM OTf was fully washed out during the PBS washing steps, a freshly made PEDOT:PSS hydrogel was soaked in D_2_O-PBS at 30 times the volume of the hydrogel. 3 mg/mL of 1,3,5-trioxane was dissolved in D_2_O-PBS to serve as an internal standard for calculating the residual concentration of EMIM OTf in the discarded wash. Fresh changes of D_2_O were made every 12 hours, and the discarded wash was analyzed via ^1^H NMR (Supplementary Fig. 11 & 12).

### Preparation of PDMS

To prepare PDMS discs, 45 parts of base was mixed with 1 part of curing agent and poured in a petri dish. To prepare 1 mm thick samples, the uncured PDMS was leveled in the petri dish and cured at 80 °C for 3 hours. To prepare Dex-PDMS, 2% w/v of dexamethasone was dissolved in THF. 1 volume of the resultant THF solution was mixed with 1 volume of PDMS base, and the THF was slowly removed in a rotary evaporator. The resultant mixture was placed in vacuum overnight to completely remove the remaing THF and to obtain a PDMS base mixture with 1% w/w dexamethasone. To prepare 1 mm thick samples, 44 parts of the dexamethasone-PDMS base mixture was mixed with 1 part curing agent and poured into a petri dish. The uncured mixture was leveled in the petri dish and cured at 80 °C for 3 hours. To prepare ∼100 μm thick films, dextran (10% in DI H_2_O, 0.05% w/v Tween-20) was spin coated at 2000 RPM on O_2_ plasma treated glass and baked at 80 °C for 1 minute. Then, a mixture of 10 parts of dexamethasone-PDMS base and 1 part of curing agent was spin coated at 1000 RPM for 2 minutes and baked at 80 °C for 30 minutes.

All *in vivo* PDMS samples were sterilized with ethylene oxide.

### Preparation of tissue adhesive

Dextran (10% in DI H_2_O, 0.05% w/v Tween-20) was spin coated at 2000 RPM on O_2_ plasma treated glass and baked at 80 °C for 1 minute. PDMS (10:1 base:curing agent, 50% in heptane) was spin coated at 2000 RPM for 2 minutes and baked at 120 °C for 10 minutes. The spin coated PDMS was treated with O_2_ plasma and incubated in APTES (2% in ethanol) for 1 hour. Then, the tissue adhesive precursor solution was prepared by dissolving 5 wt% HPU, 30 wt% AAc, 1 wt% AAc-NHS ester, 0.3 wt% MBAA, and 0.2 mol% (vs. AAc) HMPP in 95/5 ethanol/DI H_2_O and sparged in nitrogen for 10 minutes. The sparged precursor solution was sandwiched between a piece of glass with APTES-treated PDMS and a piece of glass with PTFE coating with a ∼130 μm PET spacer. The precursor solution was cured under UV (365 nm wavelength, 4 mW/cm^2^) for 30 minutes. After curing, the mold was opened, rinsed with DI H_2_O, and blow dried. Before surgical implantion, the tissue adhesive was sterilized with UV-C.

### Synthesis of polymers

Poly(glycidyl methacrylate) (PGMA) was synthesized by dissolving 10.0 g GMA (70.3 mmol), 23 mg AIBN (0.14 mmol), and 156 mg CPBD (0.703 mmol) were dissolved in 30 mL DMF. The solution was sparged with nitrogen gas for 30 minutes and reacted at 65 °C for 18 hours. The polymer was precipitated in over 10 times the volume of cold methanol and re-dissolved in DCM. The polymer was precipitated twice more and dried under vacuum for two days.

### Mutlilayer functionalization

To achieve stable adhesion of conducting hydrogels, monolayers of PGMA and PAH (PGMA-PAH method) was deposited. For the PGMA-PAA method, the Parylene C substrate was first O_2_ plasma treated and PGMA (5 mg/mL in 19:1 CHCl_3_:chlorobenzene) was spin coated at 2000 RPM, followed by baking at 120°C for 30 minutes. The baked substrate was washed in CHCl_3_ for 2 minutes and blow dried to form a PGMA monolayer. PAH (5 mg/mL 29:1 isopropanol:DI H_2_O) was spin coated at 2000 RPM on top of the PGMA monolayer and baked at 80°C for 10 minutes. The baked substrate was washed in DI H_2_O (+0.05% Tween-20) for 2 minutes, rinsed in DI H_2_O, and blow dried to form a PAH monolayer

### Measurement of electrical properties

Conductivity measurements were conducted using a standard four-point geometry (Supplementary Fig. 4a & b). Bulk hydrogels were prepared into discs (10 mm in diameter and 1 mm in thickness) using a biopsy punch, and solvent treatments were applied to the discs. Any changes in physical dimensions that resulted from the solvent treatments were accounted for when determining the conductivity values. Probes were put in direct contact with the non-implanted hydrogels fully hydrated in PBS (Fig. 2e). Conductivity measurements were also carried out for dry spin-coated thin films (Supplementary Fig. 7).

### Measurement of electrochemical properties

The electrochemical measurements were made using the PalmSens electrochemical workstation. To fabricate Au electrodes, physical vapor deposition of 10/50 nm of Ti/Au using the EvoVac (Angstron Engineering) was carried out with a shadow mask on glass substrates. For PEDOT:PSS, PGMA monolayers were deposited on the glass substrates, followed by spin coating PEDOT:PSS (13% 1:13 Triton X-100:DMSO) at 1000 RPM through a PET mask. The PET mask was peeled off, the substrate was baked at 120°C for 30 minutes, and rinsed with DI H_2_O. For ZIPH-3c, multilayer functionalization was carried out with PGMA and PAH. The resultant free amines were reacted in a tetrachloroethane solution of 2% triethylamine and 2% methacryloyl chloride. The ZIPH-3c precursor solution was spin coated at 1000 RPM for 10 seconds through a PET mask, followed by exposure to UV light (365 nm wavelength, 4 mW/cm^2^) for 30 minutes under a humidified atmosphere. The PET mask was peeled off, and three 1 minute cycles of ethanol-TFE solvent annealing was conducted to obtain ZIPH-3c films on Au electrodes. For electrochemical measurements, PDMS wells were used to hold 1x PBS. The Au electrode served as the working electrode, and a Ag/AgCl reference electrode and a platinum electrode were held in contact with PBS (Supplementary Fig. 4C). Electrochemical impedance spectroscopy (EIS) spectra were obtained over a range of 100 kHz to 0.1 Hz with an AC 10 mV sine wave and with no DC offset. Cyclic voltammetry was conducted at a scan rate of 0.1 V/s and with a voltage window of -0.1 V to +0.5 V with respect to Ag/AgCl (Supplementary Fig. 9).

### Measurement of mechanical properties

The tensile properties of hydrogels were tested on a universal testing machine (zwickiLine Z0.5, Zwick Roell Group, Ulm, Germany) with 500-N load cell at room temperature. Hydrogels were cut into JIS K6251-8 industrial standard dumbbells. For ZIPH-3c, ZIPH-u, pSB, and pSS hydrogels, the crosshead speed was set to 30 mm/s. For PEDOT:PSS hydrogel, the crosshead speed was set to 5 mm/s. Cyclic tests were also conducted (Supplementary Fig. 14).

### Cryo-TEM characterization

UV polymerized ZIPH films were prepared following a modified version of the procedures described earlier. The precursor solutions were spin-coated at 2000 RPM for 10 seconds on oxygen plasma-treated copper Quantifoil grids (200 mesh, R1.2/1.3; Electron Microscopy Sciences) supported on PDMS. A portion of the gel was scraped away to expose the mesh pattern of the grid. The hydrogel films were kept in a humidified chamber until further processing. The gels were then back blotted (force = 1, 20s) and plunge frozen into liquid ethane on a Thermo Scientific Vitrobot Mark IV. The Aquilos 2 cryo-FIB/SEM (Thermo Scientific) was used to prepare the lamella. Using the in-column sputter coater the grid was coated with platinum at 1kV and 10 mA current for 15 seconds. The grid was SEM imaged and then atlased using Maps 3.26 (Thermo Scientific) at 10kV accelerating voltage and 13 pA current to find suitable areas for milling. Prior to milling the grids were coated with a layer of organometallic platinum using a gas injection system for 10 sec. The milling process was automated into 4 steps (all at 30kV accelerating voltage): Rough Milling, Medium Milling, Fine Milling, and Thinning using 500, 300, 100, 30 pA current, respectively. The final lamella thickness was targeted between 100 -150 nm. Lamella were imaged on the Thermo Scientific Titan Krios 3Gi at 300kV at cryogenic temperature equipped with a Gatan K3 direct electron detector and BioContinuum in counting CDS mode.

### Conductive AFM

The conductive AFM (C-AFM) measurements were performed by using the C-AFM mode of Cypher ES Environmental AFM with ASTELEC-01 probe and the dual gain C-AFM probe holder (gain=1μA/1nA/V). UV polymerized ZIPH-E and ZIPH-3c films were prepared following the procedures described earlier. The precursor solutions were spin-coated at 6000 RPM for 10 seconds on oxygen plasma-treated Au films on SiO_2_ substrates and dried on a hotplate. Magnets were placed separately on Au and sample film, and were sealed with Ag paste to realize ideal contact. Before measurement, the sample bias was zeroed by applying an offset voltage from the AFM hardware. During the measurement, 1 mV bias was applied across samples and the current was converted to a voltage and then amplified and recorded by the AFM hardware. Scans were acquired using scan rates of 2 μm/s.

### Small-angle X-ray scattering

Small-angle X-ray scattering (SAXS) was performed at the Advanced Photon Source at Argonne National Laboratory on beamline 12-ID-B. Pristine PEDOT:PSS and PEDOT:PSS with 10 mg/mL of EBSA were loaded into capillary tubes (Charles Supper) and sealed. The sample-to-detector distance was 2 m, corresponding to a *q*-range of 0.003-0.5 Å^−1^.

### Grazing-incidence X-ray diffraction

Grazing-incidence X-ray diffraction (GIXD) was performed at the Advanced Photon Source at Argonne National Laboratory on beamline 8-ID-E. UV-initiated ZIPH samples were spin-coated at 1500 RPM for 10 seconds. Thermally initiated ZIPH samples were prepared by sandwiching 10 μL of the precursor solution between a piece of silicon (O_2_ plasma treated) and PET laminated coverslip (10 mm x 17 mm) and heating to 80°C for 15 minutes. PSB hydrogel samples were prepared similarly by sandwiching 3 μL of the precursor solution between a piece of silicon (O_2_ plasma treated) and PET laminated coverslip (10 mm x 17 mm) and exposing to UV light for 10 minutes. The coverslip was gently removed while submerged in water. All samples were dried on a hotplate after polymerization of solvent annealing.

### Scanning electron microscopy – energy dispersive X-ray spectroscopy

Scanning electron microscopy – energy dispersive X-ray spectroscopy (SEM-EDX) was conducted using the TESCAN LYRA3 field-emission SEM with Oxford X-Max detectors for EDX. UV-initiated ZIPH samples were spin-coated at 2000 RPM for 10 seconds. An accelerating voltage of 4 kV was used to locate and analyze the samples (Supplementary Fig. 8).

### Confocal Raman spectroscopy

Raman microscopy was conducted using the HORIBA LabRAM HR Evolution Confocal Raman Microscope. Thermally initiated ZIPH and PEDOT:PSS hydrogels were sealed between a glass slide and coverslip to prevent dehydration during measurement. An objective magnification of 10x was used to focus the 532 nm laser beam on the sample. The scattered light was analyzed using a grating of 600 lines/mm (Supplementary Fig. 3).

### Fabrication of electrodes

ZIPH-3c electrodes were fabricated using procedures shown in Supplementary Fig. 50. First, an oxygen plasma-treated 2”x3” glass slides were thoroughly washed with acetone, water, and isopropanol. To the washed slides, 2% solution of Micro-90 was spin coated at 1000 RPM and dried at 90°C for a minute. On the glass slides, 5 μm thick layers of Parylene C was deposited using the PDS 2010 Labcoter^TM^ 2 (Specialty Coating Systems). AZ 5214-E photoresist (MicroChemicals) was spin coated on the Parylene C and patterned using the Maskless Aligner MLA 150 (Heidelberg Instruments), followed by a photoresist descum process with O_2_ plasma. Then, physical vapor deposition of 10/50 nm of Ti/Au using the EvoVac (Angstron Engineering) was carried out. The photoresist-Ti/Au stack was lifted off in acetone at 40°C for 1 hour.

To achieve stable adhesion of conducting hydrogels, multilayer functionalization using PGMA and PAH was carried out. The resultant amines on the surface were functionalized by soaking in a tetrachloroethane solution of 2% triethylamine and 2% methacryloyl chloride for 2 hours at room temperature. The functionalized substrate was rinsed with chloroform and DI H_2_O. UV initiatable ZIPH precursor solution was spin coated on the functionalized substrate at 1000 RPM for 10 seconds, followed by exposure to UV light (4 mW/cm^2^) for 30 minutes under a humidified atmosphere. Immediately, the hydrogel film was subjected to three cycles of the solvent annealing process and dried at 60°C on a hotplate. The hydrogel film was rinsed with DI H_2_O and blow dried.

For PEDOT:PSS electrodes, PEDOT:PSS (13% 1:13 Triton X-100:DMSO) was spin coated at 300 RPM. Then, the film was baked at 120°C for 30 minutes. The baked film was rinsed with DI H_2_O and blow dried.

Once the hydrogel films were prepared, the outlines of the electrodes were cut with a laser cutter. To enable wired connections to the electrodes, a wet swab was used to remove the hydrogel films on top of the Au contact pads, and 0.005” Ag wires with 0.007” PFA insulation (A-M Systems) were attached to the contact pads with lead-free low temperature solder (Sn42Bi57Ag1, ChipQuik). 50 mg/mL TPU in THF was manually pipetted to cover the Au areas and dried under ambient conditions. Devices were peeled off, rinsed in DI H_2_O, and dried on a polystyrene surface under ambient conditions.

### Electrode-housing assembly

The ends of the Ag wires not soldered to the electrodes were attached to jumper wire female connectors via lead-free solder (Sn99.3Cu0.7, Yihua). The connectors were threaded through the holes of modified two-channel vascular access buttons (Instech) and were secured with a combination of Red Epoxy (Highside Chemicals) and super glue. The full assembly was sterilized in an autoclave at 121°C for 30 minutes before implantation.

### *In vitro* plasma adsorption test

Human plasma adsorption on the hydrogels was evaluated using the 3-(4-Carboxybenzoyl)quinoline-2-carboxaldehyde (CBQCA) assay and bicinchoninic acid (BCA) assay. Hydrogel samples (5 mm in diameter and 1 mm in thickness) were incubated in 150 μL of human plasma (Innovative Research, Inc.) in 48-well plates at 37°C for 2 hours. The hydrogels were washed with 1 mL of PBS 5 times and transferred to 96-well U-bottom plates. 150 μL of 1% SDS-PBS solution was added and shaken for 1 hours. 75 μL of the protein-SDS solution was transferred to 96-well flat bottom plates and diluted with 60 μL of PBS. Then, CBQCA assay was directly carried out to determine the relative amount of proteins adsorbed on the hydrogels. The fluorescence intensities at 550 nm (excitation/emission of 465/550 nm) were recorded using the TECAN Inifinite M200 Pro plate reader. For the BCA assay, 30 μL of the protein-SDS solution was transferred to 96-well flat-bottom plates and standard BCA procedures were applied. The absorbance at 562 nm was recorded using the TECAN Inifinite M200 Pro plate reader.

### Cytotoxicity assay

NIH-3T3 fibroblasts were cultured in DMEM medium (Gibco) supplemented with 10% fetal bovine serum (FBS, Gibco) and 1% penicillin/streptomycin (Gibco). Cytotoxicity of the hydrogels was determined by directly culturing fibroblasts on the hydrogels and using the LIVE/DEAD Viability Cytotoxicity Kit (Invitrogen). Hydrogel discs (5 mm in diameter and 1 mm in thickness) in 96-well flat bottom plates were pre-conditioned in 200 μL DMEM, and cells were directly seeded at a density of 1300 cells per well. After 96 hours, 5 μL of SDS solution (2 wt%) per 100 μL media were added to half of the wells and incubated at room temperature for 10 minutes to establish dead controls. 100 μL of working solution (4 μM calcein AM and 4 μM ethidium homodimer) was added to the experimental wells, and 100 μL of calcein AM solution (4 μM) or ethidium homodimer solution (4 μM) was added to the dead control wells. The cells were incubated at room temperature for 45 minutes and immediately evaluated (LIVE: 494/517 nm, DEAD: 528/617 nm) using the TECAN Infinite M200 Pro plate reader (Supplementary Fig. 13). **Implantation surgeries.** 6-to 8-week-old male C57BL/6 mice were purchased from Jackson Laboratories. Preoperatively, all mice received subcutaneous injections of 5 mg/kg of Meloxicam for pre-surgery analgesia. All mice were anesthetized using 2-4% isoflurane in oxygen, and the backs of all mice were shaved. All mice received eye lubricants to prevent dehydration-induced blindness. Then, the shaved backs of all mice were sterilized by scrubbing with betadine and alcohol pads in an alternating fashion, repeated two times.

#### Implantation of discs

Two longitudinal incisions, 1 cm in length, were made in the upper and lower backs. The blade tips of a pair of scissors were inserted into the subcutaneous space through the incisions, and two subcutaneous pockets, flanking each incision, were made via blunt tissue dissection using the blunt sides of a pair of scissors. Two implants (5 mm in diameter and 1 mm in thickness) were inserted in the subcutaneous pockets on both sides of each incision, towards the limbs of the mice, for a total of four implants per mouse. For sham control, subcutaneous pockets were made but no implants were placed. The incisions were closed via wound clips. Following surgery, all mice were given subcutaneous injections of 5 mg/kg of Meloxicam once every 24 hours for two days. Wound clips were removed after 10 days.

#### Electrode implantation

One longitudinal incision, 2 cm in length, was made in the mid-back region. The blade tips of a pair of scissors were inserted into the subcutaneous space through the incisions, and two subcutaneous pockets, flanking the incision, were made via blunt tissue dissection using the blunt sides of a pair of scissors. After carefully exposing the subcutaneous musculature by pulling back the skin on one flank, an electrode was placed on the lower, lateral side of the latissimus dorsi. For PEDOT:PSS-Dex, a Dex-PDMS backing (∼100 μm thickness) was placed on top of the electrodes. A tissue adhesive (∼100 μm thickness) was placed on top of the electrodes so that a slight overhang around the edges of the electrode adhered to the tissue underneath. The electrode placement procedure was repeated for the other subcutaneous pocket. Then, the felt attached to the vascular access button was tucked underneath the skin, and the remaining incisions longitudinally flanking the vascular access button were closed with wound clips. Following surgery, all mice were given subcutaneous injections of 5 mg/kg of Meloxicam once every 24 hours for two days. Wound clips were removed after 10 days.

### Chronic electrocardiographic recording

First, a mouse was anesthetized using 2-4% isoflurane in oxygen. The protective cap for the external connectors were removed, and the female jumper wire connectors were connected to a custom-built amplifier. An adhesive electrocardiography patch was attached to the tail and connected to ground. Signal acquisition was carried out using the Keithley 2450 (Tektronix).

### Histological processing for H&E and Masson’s trichrome staining

After 1-, 4-, or 12-weeks post-implantation, all mice were euthanized by CO_2_ asphyxiation, followed by cervical dislocation. Afterward, the backs of all the mice were shaved. The implants and the surrounding tissues were explanted and fixed in 10% neutral buffered formalin for 24 hours. After fixation, the implants and tissues were transferred to 70% ethanol. The materials were then paraffin-embedded, sectioned (5 μm), and stained for H&E using standard procedures by the Human Tissue Resource Center at the University of Chicago. Masson’s trichrome staining (MTS) was carried out in-house using standard procedures. Whole-section scans of H&E and MTS-stained tissue sections were conducted using the Olympus VS200 Slideview Slide Scanner.

### Histological analysis

ImageJ software was used to analyze the collagen density and thickness of the fibrotic capsule from MTS-stained sections. Fibrotic capsules were measured at approximately the first, second, and third quarters along the length of the widths of two halves of the implant facing the dermis, for a maximum of six measurements per biological replicate. Care was taken to omit the basement membrane, upon which the panniculus carnosus sits, from the analysis. Three to six technical replicate measurements were taken per biological replicate, depending on the quality of the tissue section. For locations where artifacts prevented accurate measurements, the measurement was either not made for the location or was taken instead at the nearest location without artifact obstruction. For collagen density calculations, the blue pixel brightness was measured with 100 μm wide slices from the surface of the implant to a distance of at least 80 μm as a function of the distance away from the surface of the implant. The technical replicate values at corresponding distances were averaged within the same biological replicate. Subsequently, from the same biological replicate, the data points within each 5 μm increment moving away from the surface of the implant were averaged to obtain pixel brightness values every 5 μm. The pixel brightness values were normalized with respect to the maximum possible pixel brightness value of 255. For measuring the thickness, layers formed only from cells were omitted from the measurements. Technical replicates were averaged to obtain individual biological replicates.

### Immunofluorescence

Materials were fixed, embedded, and sectioned as described above. Tissue sections were de-paraffinized in xylene and rehydrated using an ethanol-to-water gradient. Antigen retrieval was conducted in citrate buffer (pH 5) for 45 minutes at 55°C. Samples were then rinsed in water, permeabilized by incubating in 10% DMSO in PBS for 5 minutes, washed in PBS-tween 20 (PBS-T), and rinsed in water again. Thereafter, the samples were blocked for 1 hour using a 1% bovine serum albumin (BSA) solution. The blocking buffer was removed, and the sections were incubated in either the macrophage panel or fibrosis panel antibodies overnight at 4°C. The macrophage panel antibodies included: rat anti-mouse F4/80 (1:100 dilution, Abcam, Cat. #ab6640), rabbit anti-mouse CD86 (1:200 dilution, Biorbyt, Cat. #orb49101), and goat anti-mouse CD206 (1:100 dilution, R&D Systems, Cat. #AF2535). The fibrosis panel antibodies included: rat anti-mouse F4/80 (1:100 dilution, Abcam, Cat. #ab6640), rabbit anti-mouse collagen I (1:100 dilution, Bio-Rad, Cat. #2150-1410), and goat anti-mouse αSMA (1:100 dilution, Novus Biologicals, Cat. #NB300-978). The next day, the sections were washed twice in PBS-T and once in PBS. The washed sections were incubated for 2 hours at room temperature in a secondary antibody cocktail, consisting of donkey anti-rat IgG AlexaFluor 647 (1:200 dilution, Southern Biotech, Cat. #OB643031), donkey anti-rabbit IgG AlexaFluor 488 (1:500 dilution, Fisher Scientific, Cat. #A21206), and donkey anti-goat IgG AlexaFluor 555 (1:500 dilution, Fisher Scientific, Cat. #A21432). Afterward, the sections were washed twice with PBS-T and once with PBS. The washed sections were counterstained with 4’,6-diamidino-2-phenylindole (DAPI, 1:1000 dilution, Thermo Scientific) for 2 minutes and washed with PBS. Prolong Antifade Gold mounting medium (Invitrogen) was added, and the sections were sealed with coverslips and nail polish. Images were acquired using either Zeiss Axiovert 200m inverted epifluorescence microscope with a Hamamatsu Flash 4.0 camera run by SlideBook 6.0 software (Intelligent Imaging Innovations). Background fluorescence was determined by tissue sections stained only with secondary antibodies and with no primary antibodies (Supplementary Fig. 31).

### Cytokine analysis

Cytokine quantifications were performed using a 14-cytokine LegendPlex custom panel (BioLegend). After 1-or 4-weeks post-implantation, all mice were euthanized by CO_2_ asphyxiation, followed by cervical dislocation. Afterward, the backs of all the mice were shaved. The explanted implant and associated tissues were frozen immediately on dry ice and stored in a -80°C freezer for later use. The frozen tissues were thawed at room temperature and minced with scissors. The minced implants and tissues were transferred to lysis matrix tubes (MP Biomedicals) with 300 μL of T-PER solution (Thermo Scientific), consisting of 5 mM EDTA and protease inhibitor (Thermo Scientific). Three cycles of homogenization were conducted on the FastPrep-24 5G homogenizer (MP Biomedicals) with three minutes on ice between cycles. Subsequently, the tubes were centrifuged for 20 minutes at 10,000 g’s at 4°C, after which a fat layer, a protein layer, and a debris layer formed. Avoiding the fat layer, the protein layer was transferred to low-bind tubes (Fisher Scientific) and centrifuged again for 10 minutes at 10,000 g’s at 4°C. Similar to the first centrifugation step, the protein layer was transferred to fresh low-bind tubes. The centrifugation and protein layer transfer step were conducted once more. The protein solutions were stored in a -80°C freezer for future analysis.

The protein solutions were thawed at room temperature, and LegendPlex assay was conducted using the un-diluted protein solutions, following manufacturer’s protocol on the Agilent Penteon 5-30 flow cytometer. A C3a ELISA kit (Novus Biologicals), a decorin ELISA kit (R & D Systems), and an endostatin ELISA kit (Abcam) were used following manufacturer’s instructions. The protein solution was diluted 20-fold in PBS for C3a ELISA, 5000-fold for decorin ELISA, and 1000-fold for endostatin ELISA. The absorbancees for the ELISAs were recorded at a wavelength of 450 nm using the TECAN Inifinte M200 Pro plate reader.

### Flow cytometry

After 1- or 4-weeks post-implantation, all mice were euthanized by CO_2_ asphyxiation, followed by cervical dislocation. The backs of all the mice were shaved, and subsequently, a depilatory cream was applied for one minute. Afterward, the cream and the hair were washed off with water and 70% ethanol. Approximately 2 cm^2^ of skin, including the implant, was extracted from the back and placed on ice. The implants and associated tissues were finely minced with scissors and digested by adding 200 μL of RPMI-1640 (Sigma), supplemented with Liberase TL (0.5 mg/mL, Roche) and DNase I (0.5 mg/mL, Grade II, Roche). The samples were shaken for 90 minutes at 37°C. Immediately, 200 μL of ice-cold dissociation buffer (1x PBS, 10 mM EDTA, 2% FBS) were added to halt digestion. The digested tissues were pestled through 70-μm cell strainers (Fisher Scientific) and rinsed with 2 mL of ice-cold dissociation buffer. Then, the washed digested tissues were centrifuged for 8 minutes at 400 g’s at 4°C and resuspended in 200 μL of FACS buffer (eBiosciences). The enriched single-cell suspension was transferred to 96-well U-bottom plates, centrifuged, and resuspended in 200 μL of FACS buffer another time.

Aliquots of the single-cell suspension were then stained with propidium iodide (1:200 dilution, BD Bioscences) and counted on an Agilent Penteon 5-30 flow cytometer. Subsequently, 10^6^ cells were incubated in LIVE/DEAD Fixable Blue Dead Cell Stain Kit (1:500 dilution, Invitrogen) for 20 minutes on ice followed by washing with FACS buffer. The cells were treated with Fc block (1:50 dilution, TruStain FcX Plus, BioLegend) for 10 minutes on ice followed by washing with FACS buffer. Then, the cells were stained with the flow cytometry panel shown below. The cells were stained with CCR2 at 37 °C for 1 hour, and the cell surface stains were carried out on ice for 20 minutes. The cells were fixed with Cytofix/Cytoperm buffer (BD Bioscences), intracellular stains were carried out at room temperature, and run on the Cytek Aurora flow cytometer the next day. All analyses were performed in FlowJo Flow Cytometry Analysis Software (Treestar) using the gating strategy shown in Supplementary Fig. 27. PE was used as the exclusionary channel for CD31, Tie-2, Ter-119, and Ep-CAM, which are collectively abbreviated as Lin. Macrophages were defined as CD45^+^ CD11b^+^ Ly-6G^-^ Ly-6C^-/lo^ F4/80^hi^, monocytes were defined as CD45^+^ CD11b^+^ Ly-6G^-^ Ly-6C^hi^ F4/80^lo^, neutrophils were defined as CD45^+^ CD11b^+^ Ly-6G^+^, T cells were defined as CD45^+^ CD3^+^, and fibroblasts were defind as CD45^-^ Lin^-^ CD140a^+^.

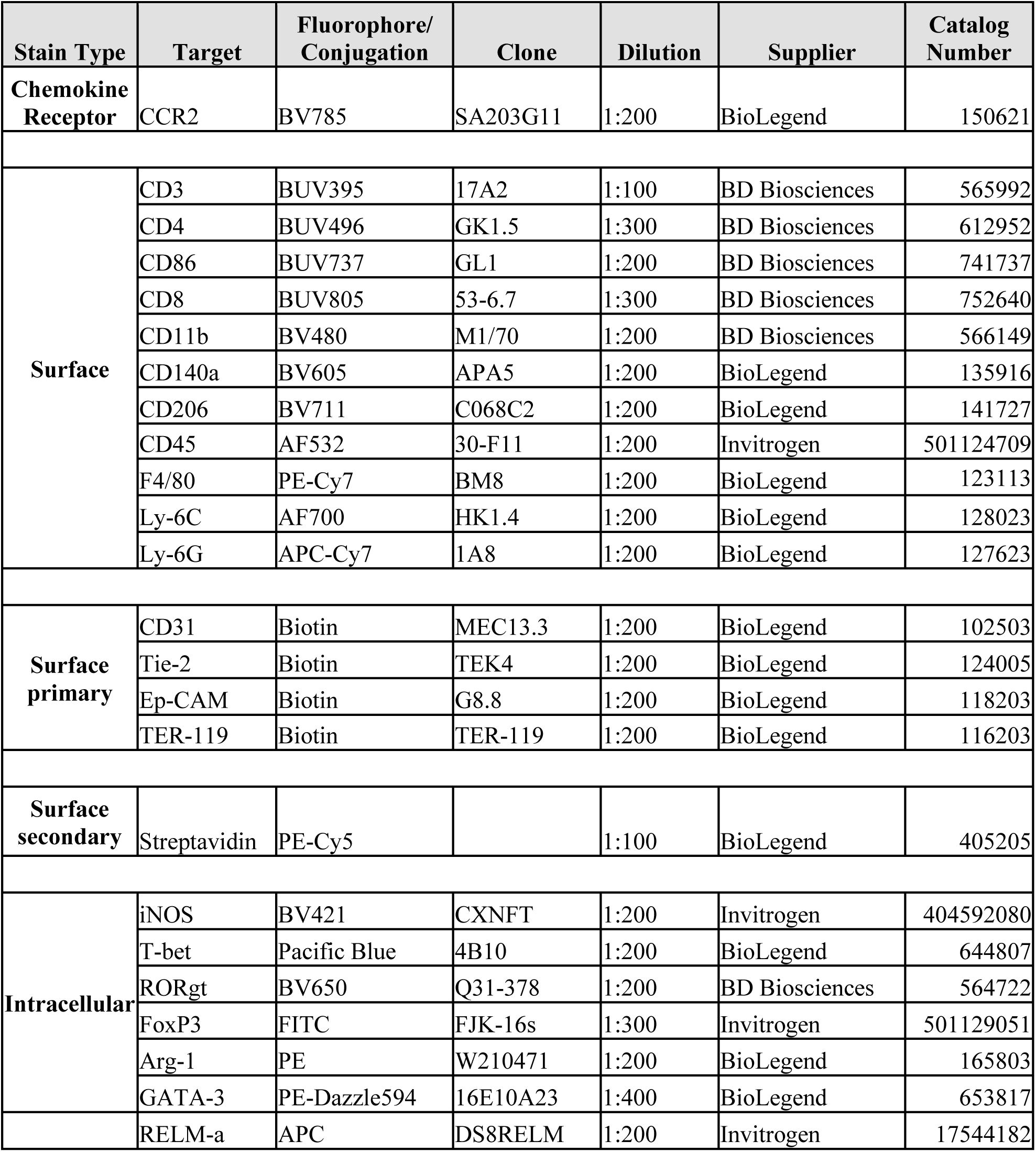

| Stain Type | Target | Fluorophore/<br>Conjugation | Clone | Dilution | Supplier | Catalog<br>Number |
| --- | --- | --- | --- | --- | --- | --- |
| **Chemokine**<br>**Receptor** | CCR2 | BV785 | SA203G11 | 1:200 | BioLegend | 150621 |
| **Surface** | CD3 | BUV395 | 17A2 | 1:100 | BD Biosciences | 565992 |
|  | CD4 | BUV496 | GK1.5 | 1:300 | BD Biosciences | 612952 |
|  | CD86 | BUV737 | GL1 | 1:200 | BD Biosciences | 741737 |
|  | CD8 | BUV805 | 53-6.7 | 1:300 | BD Biosciences | 752640 |
|  | CD11b | BV480 | M1/70 | 1:200 | BD Biosciences | 566149 |
|  | CD140a | BV605 | APA5 | 1:200 | BioLegend | 135916 |
|  | CD206 | BV711 | C068C2 | 1:200 | BioLegend | 141727 |
|  | CD45 | AF532 | 30-F11 | 1:200 | Invitrogen | 501124709 |
|  | F4/80 | PE-Cy7 | BM8 | 1:200 | BioLegend | 123113 |
|  | Ly-6C | AF700 | HK1.4 | 1:200 | BioLegend | 128023 |
|  | Ly-6G | APC-Cy7 | 1A8 | 1:200 | BioLegend | 127623 |
| **Surface**<br>**primary** | CD31 | Biotin | MEC13.3 | 1:200 | BioLegend | 102503 |
|  | Tie-2 | Biotin | TEK4 | 1:200 | BioLegend | 124005 |
|  | Ep-CAM | Biotin | G8.8 | 1:200 | BioLegend | 118203 |
|  | TER-119 | Biotin | TER-119 | 1:200 | BioLegend | 116203 |
| **Surface**<br>**secondary** | Streptavidin | PE-Cy5 |  | 1:100 | BioLegend | 405205 |
| **Intracellular** | iNOS | BV421 | CXNFT | 1:200 | Invitrogen | 404592080 |
|  | T-bet | Pacific Blue | 4B10 | 1:200 | BioLegend | 644807 |
|  | RORgt | BV650 | Q31-378 | 1:200 | BD Biosciences | 564722 |
|  | FoxP3 | FITC | FJK-16s | 1:300 | Invitrogen | 501129051 |
|  | Arg-1 | PE | W210471 | 1:200 | BioLegend | 165803 |
|  | GATA-3 | PE-Dazzle594 | 16E10A23 | 1:400 | BioLegend | 653817 |
|  | RELM-a | APC | DS8RELM | 1:200 | Invitrogen | 17544182 |

### NanoString analysis

Gene expression was evaluated using a custom-built multiplexed 50-gene FBR panel (NanoString Technologies). After 4-weeks post-implantation, all mice were euthanized by CO_2_ asphyxiation, followed by cervical dislocation. Afterward, the backs of all the mice were shaved. The explanted material and associated tissues were soaked in RNAlater solution (Invitrogen) overnight at 4°C. The next day, the excess RNAlater solution was discarded, and the tissue samples were stored at -80°C. For RNA isolation, the tissues were thawed, finely diced, and 600 μL of Trizol reagent (Invitrogen) with 10 μL/mL of 2-mercaptoethanol was added to approximately 100 mg of tissue. The tissues were mechanically ground down and incubated for 5 minutes at room temperature. After the addition of 120 μL of chloroform, the tube was shaken vigorously for 15 seconds and incubated for 2 minutes at room temperature. The tubes were centrifuged at 12,000 xg for 15 minutes at 4°C. 350 μL of the aqueous phase was transferred to columns from the RNeasy mini kit (Qiagen), and the RNA isolation was completed using the manufacturer’s instructions. All samples were checked for a 260/280 ratio of 2 before proceeding to further steps. 100 ng of RNA was processed according to NanoString manufacturer protocols, and RNA levels (absolute copy numbers) were obtained via nCounter (NanoString Technologies). Group samples were analyzed using nSolver analysis software (NanoString Technologies). *GUSB, ACTB, B2M,* and *TBP* were used as reference genes.

## Acknowledgements

This work is supported by the US National Institutes of Health Director’s New Innovator Award (1DP2EB034563), the National Science Foundation (DMR-2105367), and the start-up fund from the University of Chicago. S.Wai acknowledges the support from the National Defense Science and Engineering Graduate Fellowship Program and the Office of Naval Research. Parts of this work were carried out at the Soft Matter Characterization Facility of the University of Chicago. This work made use of the shared facilities at the University of Chicago Materials Research Science and Engineering Center, supported by National Science Foundation under award number DMR-2011854. This work made use of the Focused Ion Beam-Scanning Electron Microscopy (FIB-SEM) core facility (RRID:SCR_025212) in the Department of the Geophysical Sciences at the University of Chicago. We thank T. Lavoie and the University of Chicago Advanced Electron Microscopy Core Facility (RRID:SCR_019198) for their assistance with cryogenic transmission electron microscopy.in conducting cryo-TEM. This work made use of the Pritzker Nanofabrication Facility, part of the Pritzker School of Molecular Engineering at the University of Chicago, which receives support from Soft and Hybrid Nanotechnology Experimental (SHyNE) Resource (NSF ECCS-2025633), a node of the National Science Foundation’s National Nanotechnology Coordinated Infrastructure [RRID: SCR_022955]. This research was performed on APS beam time awards (DOI: https://doi.org/10.46936/APS-173634/60014531 and https://doi.org/10.46936/APS-168076/60014530) from the Advanced Photon Source, a U.S. Department of Energy (DOE) Office of Science user facility operated for the DOE Office of Science by Argonne National Laboratory under Contract No. DE-AC02-06CH11357. We thank the University of Chicago Animal Resources Center (RRID:SCR_021806) for animal housing and use of their facility and equipment. All the animal experiments performed in this research were approved by the Institutional Animal Care and Use Committee of the University of Chicago under the protocol ACUP 72702. We thank The University of Chicago Human Tissue Resource Center (RRID:SCR_019199) for their assistance with embedding, sectioning, and H&E staining. Whole slide scanning and immunofluorescence imaging was performed in the Integrated Light Microscopy Core at the University of Chicago (RRID: SCR_019197), which receives financial support from the Cancer Center Support Grant (P30CA014599). Flow cytometry was performed at the Cytometry and Antibody Technology Facility at the University of Chicago (RRID:SCR_017760), which receives financial support from the Cancer Center Support Grant (P30CA014599). RNA level determination using the NanoString nCounter was performed at the University of Chicago Human Immunological Monitoring Core (RRID:SCR_017916).

## Author contributions

Wang conceived the research. S.Wai, M.V.T., and S.Wang designed the experiments. S.Wai prepared and characterized the materials, synthesized polymers, and fabricated devices. S.Wai conducted *in vitro* and *in vivo* experiments and analyzed data. J.W. and R.L. assisted in carrying out *in vivo* experiments. S.K. provided conceptual advice and technical support for *in vivo* experiments. N.L., Y.D., and J.S. performed the GIXD characterization. Y.D. performed C-AFM. T.L. carried out sample processing and imaging for cryo-TEM. S.S. and M.W. provided conceptual advice and technical support for device fabrication. T.F. provided conceptual advice and technical support for circuits involved in ECG measurements. K.C.S. provided conceptual advice on the behavior of ionic polymers. S.Wai, M.V.T., and S.Wang co-wrote the paper. All authors reviewed and commented on the manuscript.

