## Supplementary Information for "Zwitterionic hydrogel designs for conducting polymers enable bioelectronics with suppressed foreign body responses"

*Corresponding author.

**Contents**

**Supplementary Discussion ………………………………………………………………………2**

**Supplementary Figures…………………………………………………………………………..5**

**Supplementary Tables…………………………………………………………………………..58**

**Supplementary Discussion**

*Effects of solvent treatment on PEDOT:PSS surface properties*

We assumed that the surface properties of PEDOT:PSS are largely governed by PSS since PSS forms a hydrophilic shell around the PEDOT core. For this assumption to be valid, the change in PSS to PEDOT ratio needs to be negligible. Raman spectroscopy confirmed that any changes in PSS content were negligible after solvent annealing and PBS washes (Supplementary Fig. 3).

*Mechanical effects on the FBR*

In general, the lower the Young’s modulus, the milder the FBR for any given material^10,70,71^. To eliminate confounding effects from variations in mechanical properties, all implant types tested were kept at a Young’s modulus of around 10 kPa, except for PEDOT:PSS implants. Due to limitations in the intrinsic mechanical properties of PEDOT:PSS, the Young’s moduli of the PEDOT:PSS implants were limited to 720 kPa. To account for the higher Young’s modulus of PEDOT:PSS compared to other implant types in the study, PSS hydrogels with a Young’s modulus of 10 kPa to match the rest of the implants were included. Because PEDOT:PSS is composed of a hydrophobic PEDOT core and a hydrophilic PSS shell that face outwards towards the aqueous environment, the PSS hydrogel is meant to mimic the surface properties of PEDOT:PSS.

*Immune response type and macrophage polarization states in the FBR*

The scientific consensus is that type 2 immunity and M2 macrophages (or alternatively activated macrophages) are responsible for the progression of fibrotic diseases^72–77^. However, a literature survey^51^ by Kara Spiller *et al.* on the effects of macrophage polarization state on specifically the FBR found that there are just as many studies claiming that M1 macrophages are associated with the exacerbation of FBR as there are for M2 macrophages. This could be due to the fact that macrophage polarization states are highly dynamic and exist on a continuum^37,51^, complicating the interpretation of data. Some suggested that anti-inflammatory macrophages, which have commonly been categorized as M2 macrophages, should instead be classified as regulatory macrophages (M_reg_s)^78^, in accordance with the anti-inflammatory role of regulatory T cells (T_reg_s). Regardless, we have decided to describe macrophages using the more widely recognized M1/M2 dichotomy in our work.

*The macrophage-T cell interplay for ZIPH-3c*

The lack of fibrosis associated with ZIPH-3c more closely aligns with the scientific consensus on fibrotic diseases, in which the type 1 immune response counteracts the progression of fibrosis^72–77^. At the 4-week timepoint, ZIPH-3c is associated with a high M1/M2 macrophage ratio (based on CD86/CD206 and iNOS/Arg-1 ratios), a high fraction of T_h_1 cells with respect to the total number of CD4^+^ T cells, a high fraction of T-bet^+^ CD8^+^ T cells with respect to the total number of CD8^+^ T cells, and a high CD8^+^/CD4^+^ T cell ratio (Supplementary Fig. 24), collectively implying a type 1 immune response. However, cytokines associated with T_h_1 activation, such as IFN-γ, TNF-α, and IL-12, except for IL-18 at the 8-week timepoint, were not upregulated at all the timepoints tested (Supplementary Fig. 32-36). This could be due to ZIPH-3c being associated with the highest fraction of T_reg_s with respect to the total number of CD4^+^ T cells, consistent with previous observations that T_reg_s can suppress IFN-γ expression in T_h_1 cells without disrupting T-bet expression^79^. The strong presence of T_reg_s may also be a contributing factor for ZIPH-3c being associated with a very low number of T cells, although this does not explain why ZIPH-3c is associated with a very high number of macrophages. If T_reg_s are responsible for counteracting the activation of the type 1 immune response, it is probably IL-10-independent, as reported previously^79^, since the concentration of IL-10 is not particularly high compared to the other implants. This also raises the question of what the role of the type 1 immune response is in this context if there is suppressed T_h_1 activation. This complicated interplay should be investigated in future studies.

*The role of MCP-1 in the suppression of the FBR for ZIPH-3c*

MCP-1 is an inflammatory cytokine that is involved in T_h_2 polarization^80,81^ and the recruitment of monocytes and monocyte-derived macrophages^81^. MCP-1 could be responsible for the slight increase in T_h_2 numbers 1-week post-implantation. But it does not match the massive upregulation of MCP-1, the T_h_2 population is lower compared to the other conditions at later timepoints, and ZIPH-3c seems to have a type 1 immunity bias, as established previously (Supplementary Fig. 23-25). MCP-1 could be responsible for the high number of macrophages, but this also does not provide a satisfactory explanation. At the level that is observed for ZIPH-3c, MCP-1 should lead to high levels of CCR2^+^ macrophage recruitment, but ZIPH-3c is associated with one of the smallest fractions of CCR2^+^ macrophages relative to the total number of macrophages at all timepoints (Supplementary Fig. 23-25). This leaves two possibilities: (1) the monocyte-derived CCR2^+^ macrophages rapidly downregulate CCR2 expression upon maturation due to some unknown factors; and (2) the chemotactic effects of MCP-1 are inhibited by some unknown factors, and the high number of macrophages is not a direct consequence of the massive MCP-1 upregulation. Either way, further investigation is needed to elucidate the role of MCP-1 in the immune response to ZIPH-3c.

*Fibrosis and angiogenesis after 12-weeks post-implantation*

Both the collagen density and thickness of ZIPH-3c decrease between weeks 12 and 24. One plausible explanation is that macrophages are replacing the collagen layer as the fibrotic capsule is remodeled over time. This can be seen from the Masson’s trichrome staining results (Fig. 3 and Supplementary Fig. 15), in which the area occupied by collagen at 12-weeks post-implantation is occupied by cells by 24-weeks post-implantation. Further investigations are needed to explain this result.

Given the substantial upregulation in MCP-1 at all timepoints and, to a lesser extent, VEGF at early timepoints (Supplementary Fig. 32-36), one would expect there to be equally substantially increased angiogenesis around the implant for ZIPH-3c. However, this is not observed. There may be endogenous angiogenesis inhibitors that being upregulated in the peri-implant region, but further investigations are needed to find the cause of this observation. For electrophysiological devices, the lack of angiogenesis is of minor concern unless it causes necrosis because access to ions in the fluid in the peri-implant region is what determines signal quality, not blood supply.

*Effect of the mechanical properties of electrodes on the FBR*

All electrodes were made with 5-μm-thick Parylene C substrates. The difference in mechanical properties between electrode types comes from the conductive layers, ZIPH-3c and PEDOT:PSS, but both ZIPH-3c and PEDOT:PSS layers are less than 1-μm, which were assumed to have negligible influences on the overall mechanical properties of the much stiffer Parylene C substrate (E ~ GPa), eliminating mechanical effects as the confounding variable on signal quality.

*ECG recordings*

The large variation in voltage retention at early timepoints may originate from the shifting of the electrodes and the dynamic nature of the extracellular matrix and cell population in the peri-implant region. Even though the electrodes are fixed to the surface of the muscles on the lower rib cage with tissue adhesives, they may shift until tissues grow in to stabilize the electrode location and orientation. For PEDOT:PSS-Dex, even though no fibrotic tissue forms, loose layers of adipose and miscellaneous tissue types can form around the edges to stabilize the electrode, as can be seen from the histological sections of Dex-PDMS implants. The reason there is a big initial drop in voltage retention in week 1 may also be due to the shift in the electrode location and orientation. Lastly, ZIPH-3c and PEDOT:PSS-Dex electrodes have similar voltage retention at later timepoints, possibly because the diffusion and migration of ions in hydrogel matrices are dictated by water content, as suggested by some hydrogel diffusion models^82^. The lower collagen density for ZIPH-3c leads to a fibrotic capsule with a higher water content compared to that of PEDOT:PSS, improving signal quality.

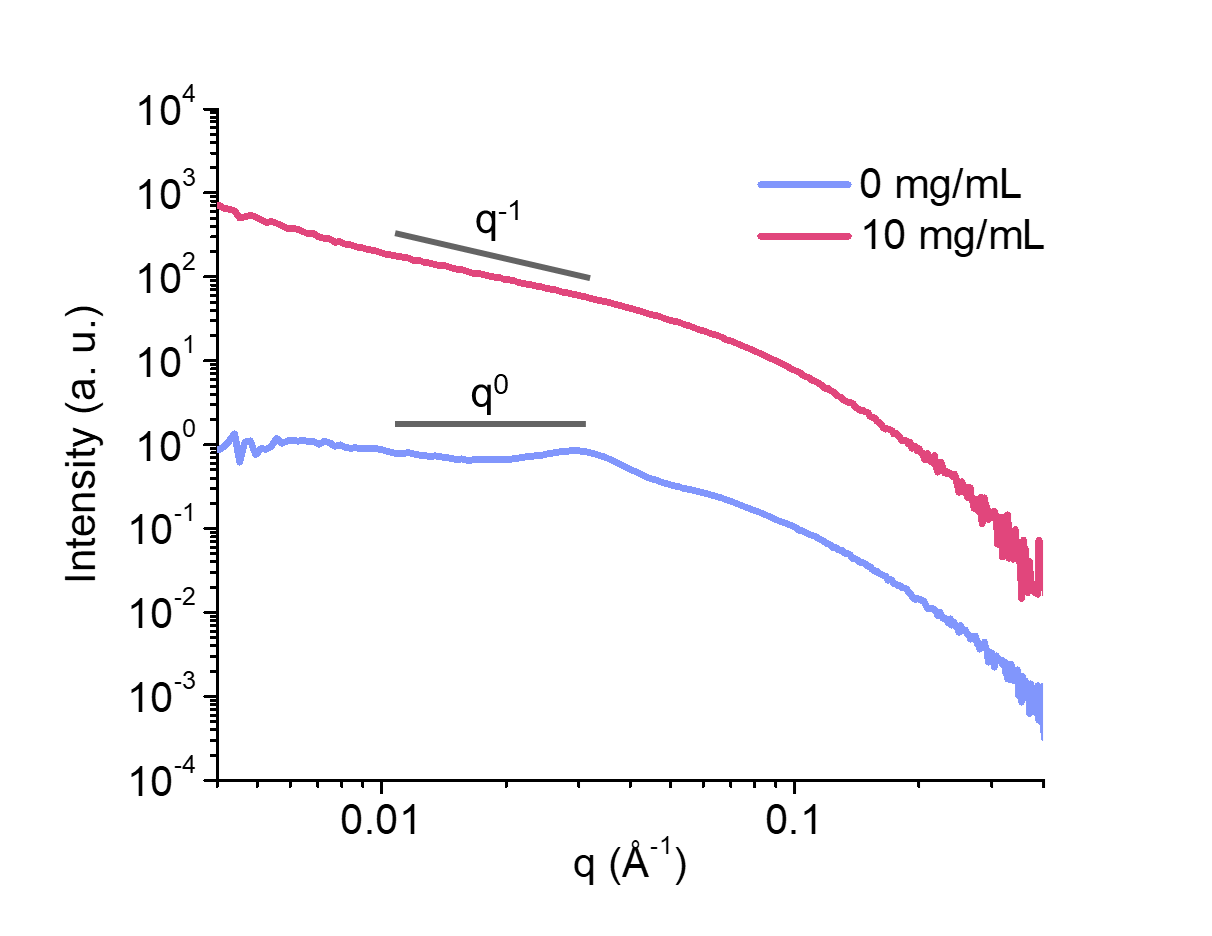
**Supplementary Figures**

**Supplementary Figure 1.** Small-angle X-ray scattering (SAXS) of PEDOT:PSS with 0 or 10 mg/mL of 4-ethylbenzenesulfonic acid (EBSA). The transition of the slopes at low- to mid-q from q^0^ to q^-1^ signifies that the PEDOT:PSS nanoparticles transition from spheroidal to rod-like particles after the addition of EBSA.

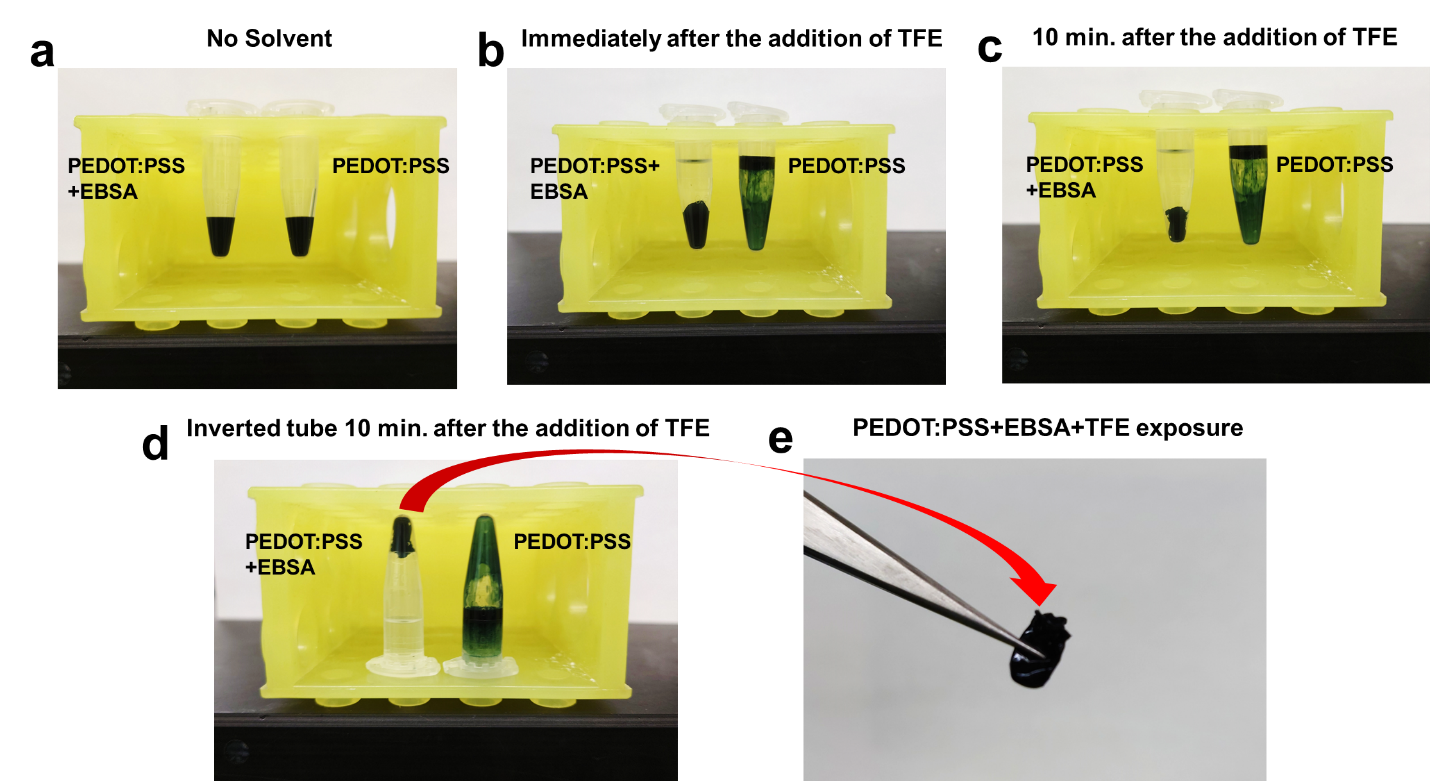

**Supplementary Figure 2.** The combination of EBSA and TFE facilitates the formation of solid, sponge-like pieces of PEDOT:PSS. **a,** Comparison of PEDOT:PSS+EBSA and PEDOT:PSS before TFE addition. No obvious difference is observed. **b,** Comparison immediately after TFE addition. With EBSA, PEDOT:PSS solidifies. Without EBSA, TFE sinks, and the PEDOT:PSS suspension floats to the top difference due to the difference in density compared to water (1.39 g/cm^3^ vs. 0.997 g/ cm^3^). **c,** Comparison 10 min. after TFE addition. The PEDOT:PSS+EBSA solid shrinks with time. **d,** Comparison after the tubes are inverted. PEDOT:PSS+EBSA does not flow down because it solidified, but PEDOT:PSS without EBSA the mixture flows down. **e,** PEDOT:PSS+EBSA solid picked up with a tweezer.

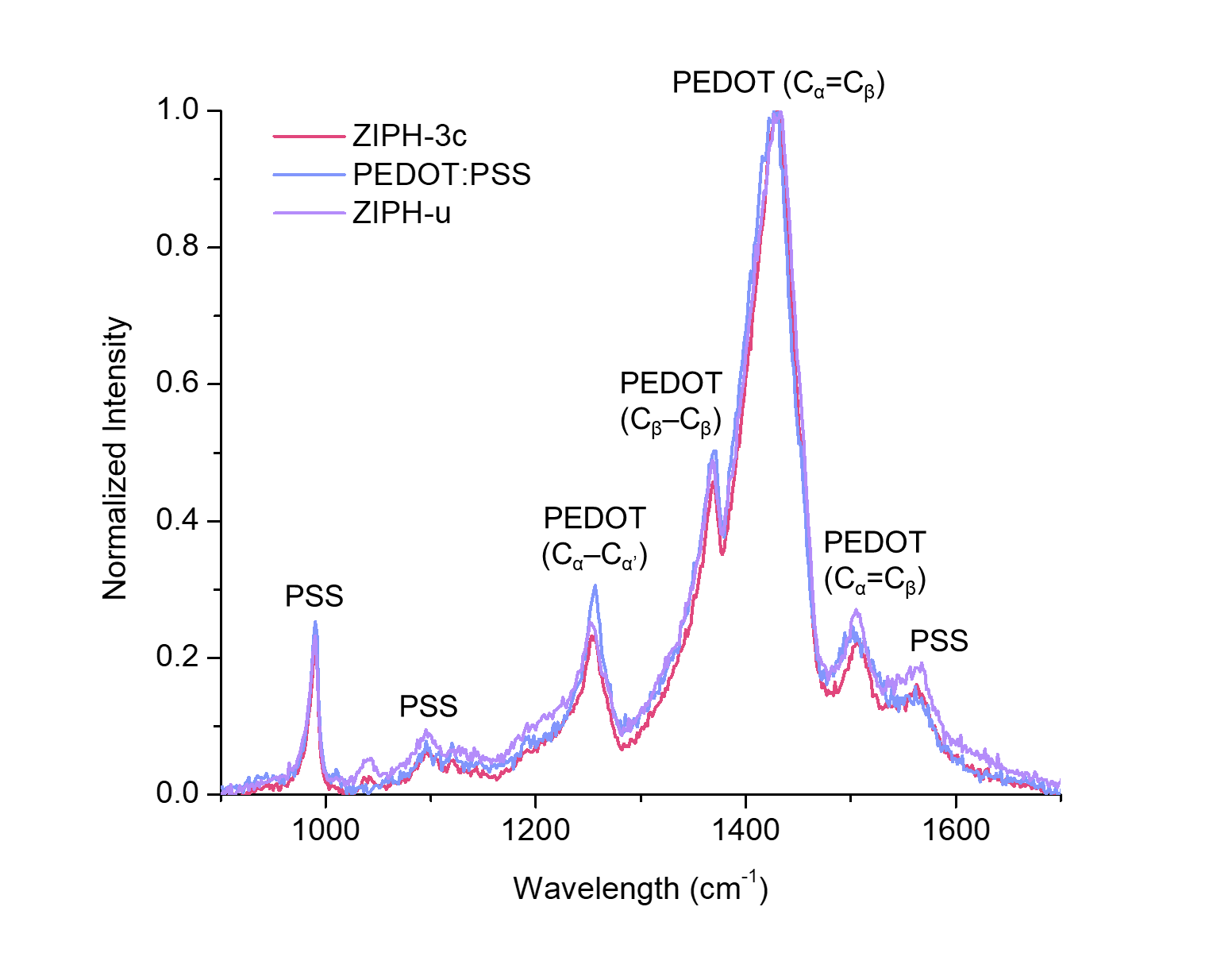

**Supplementary Figure 3.** Comparison of Raman spectra of ZIPH-3c, PEDOT:PSS hydrogel, and ZIPH-u. The spectra overlap very closely which implies that the differences in PEDOT to PSS ratios are negligible.

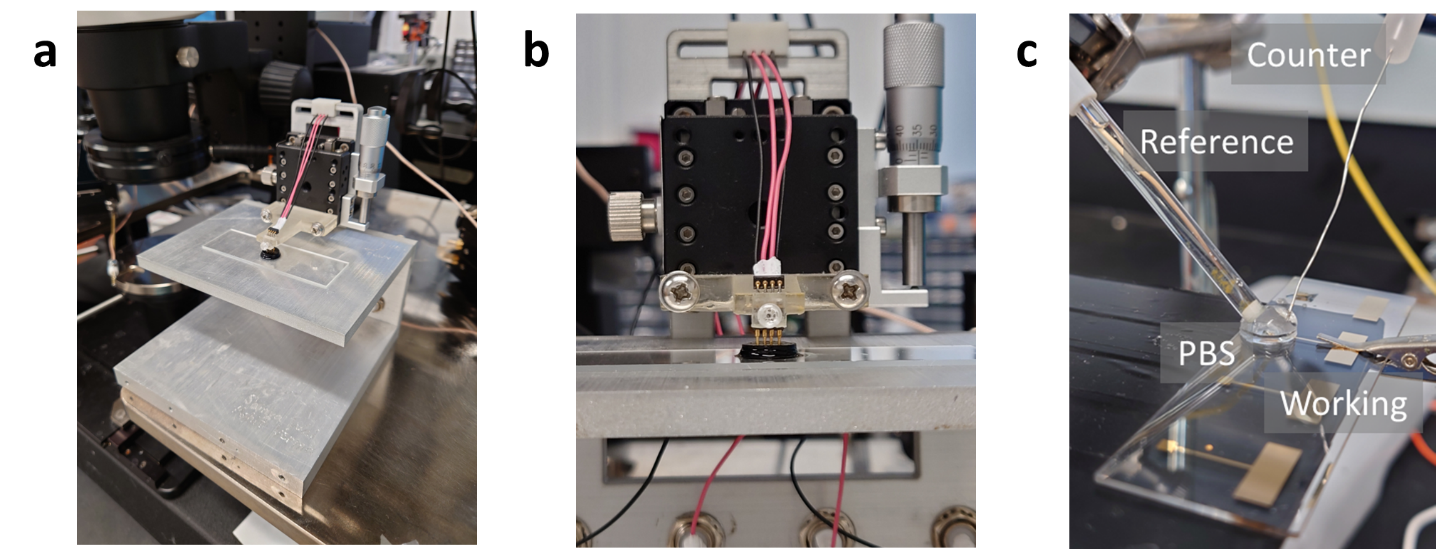

**Supplementary Figure 4. a**, 4-point probe setup. **b**, Close up view of the 4-point probe setup. **c**, 3-terminal electrochemical measurement setup.

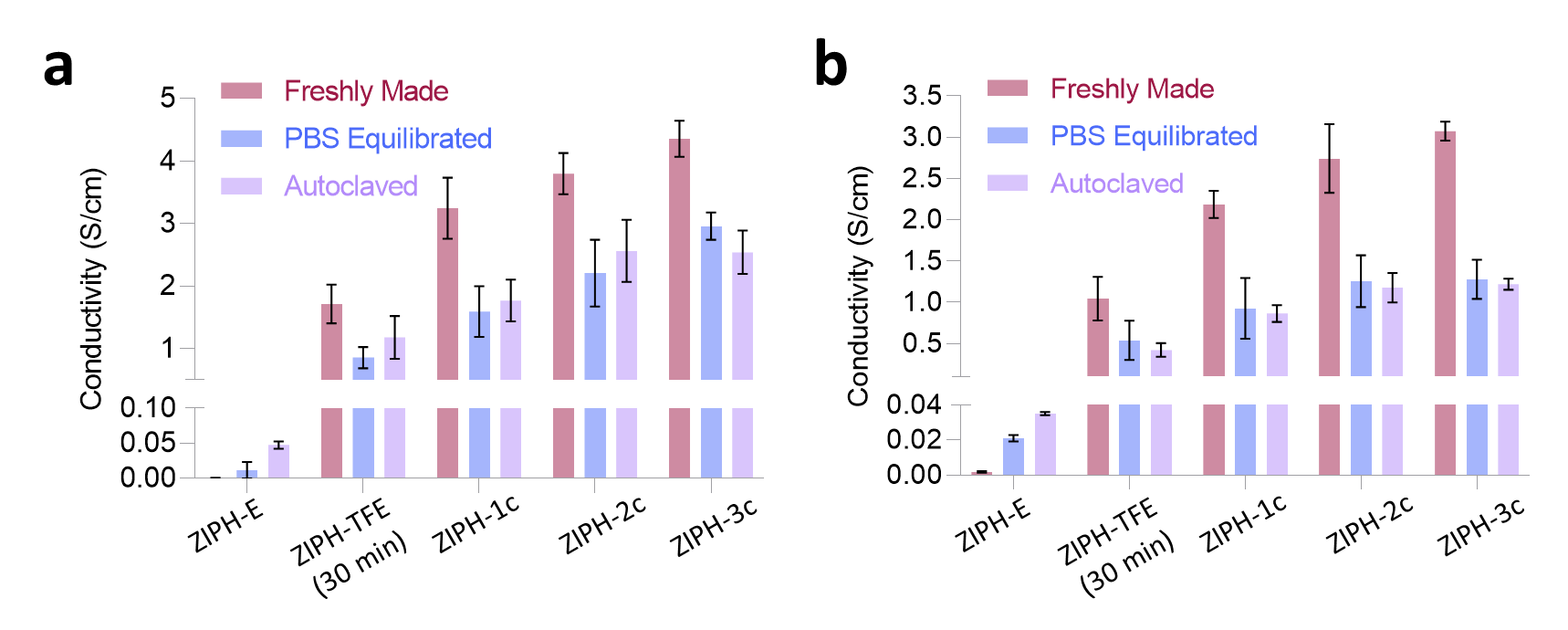

**Supplementary Figure 5.** Crosslinking densities, solvent annealing, and post-treatments affect the conductivity of ZIPH **a,** Conductivity comparisons of ZIPH made with 1.0 mol% crosslinking density. **b,** Conductivity comparison of ZIPH made with 1.8 mol% crosslinking density. Crosslinking mol% values were calculated with respect to the monomer concentration.

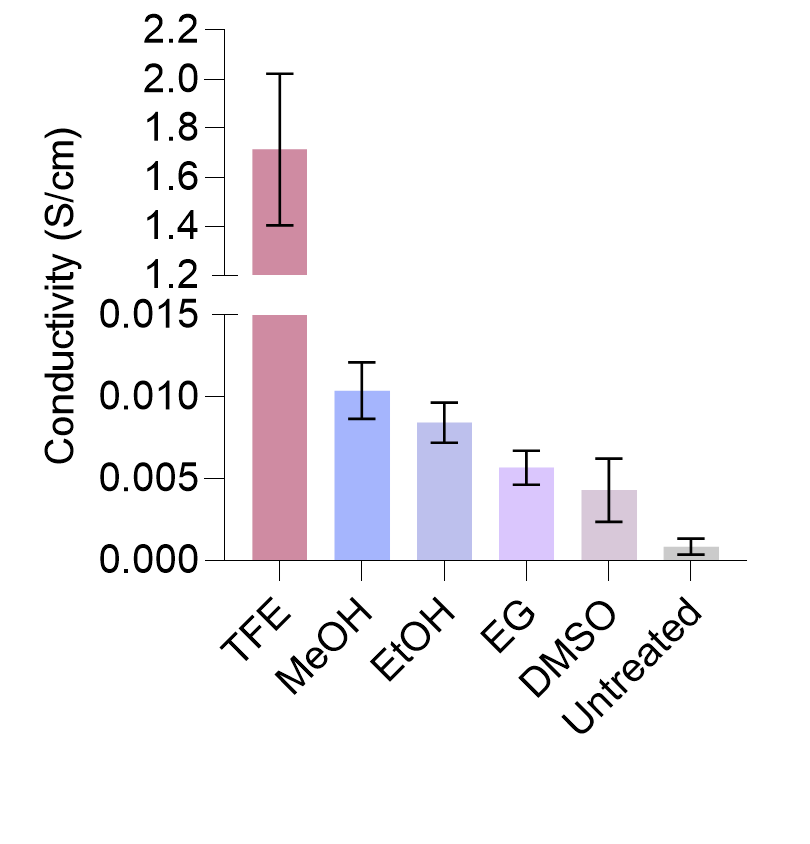
**Supplementary Figure 6.** The conductivities of ZIPH-E bulk samples treated with excess amounts of various solvents for 30 minutes. EG, ethylene glycol; DMSO, dimethyl sulfoxide

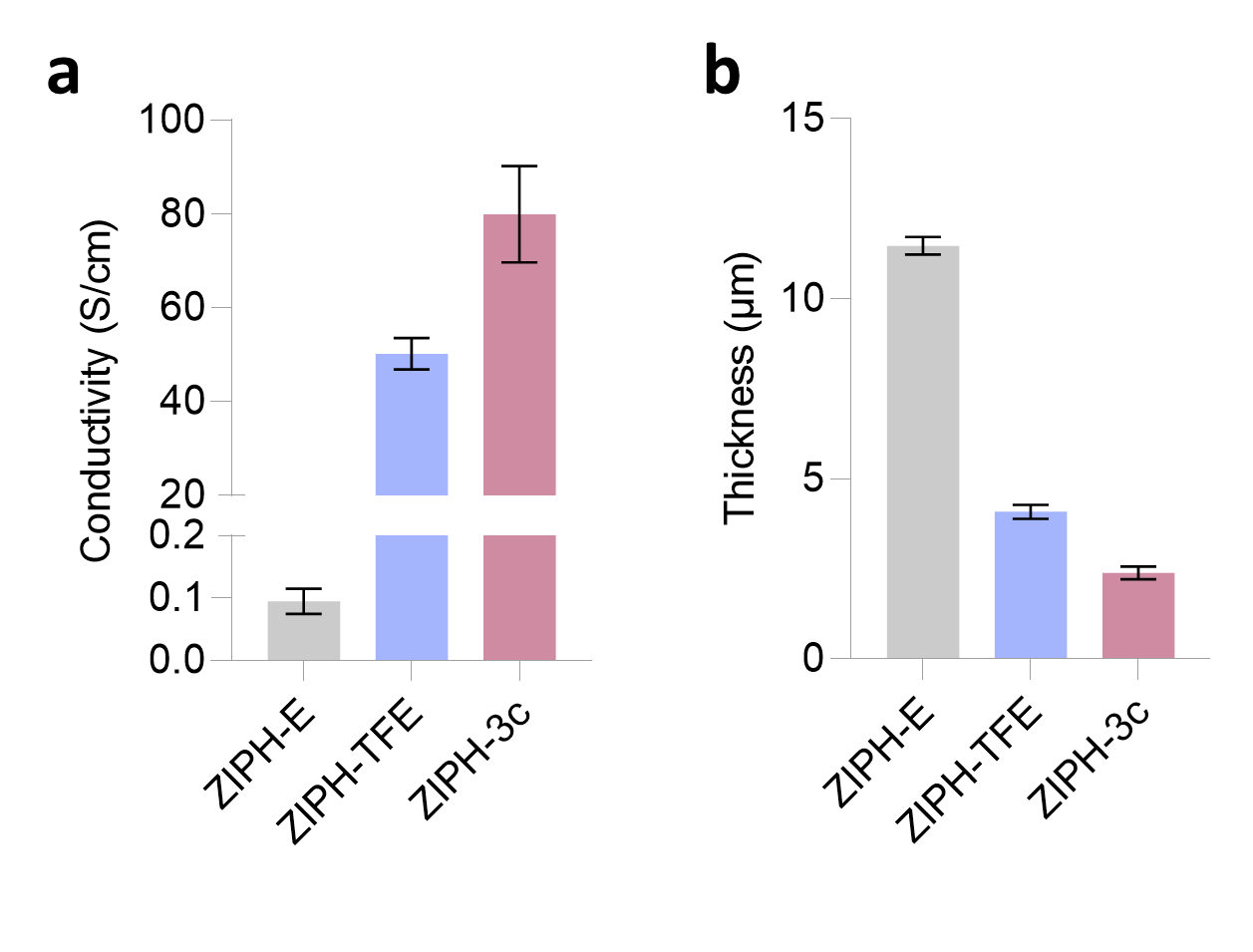
**Supplementary Figure 7.** Comparisons of **a,** conductivity values and **b,** thicknesses of thin film ZIPH samples subjected to different treatments. Note that before the solvent annealing steps, ZIPH-TFE and ZIPH-3c are the same as ZIPH-E. Thin film ZIPH-3c has one of the highest conductivity values ever reported for double network PEDOT:PSS hydrogels.

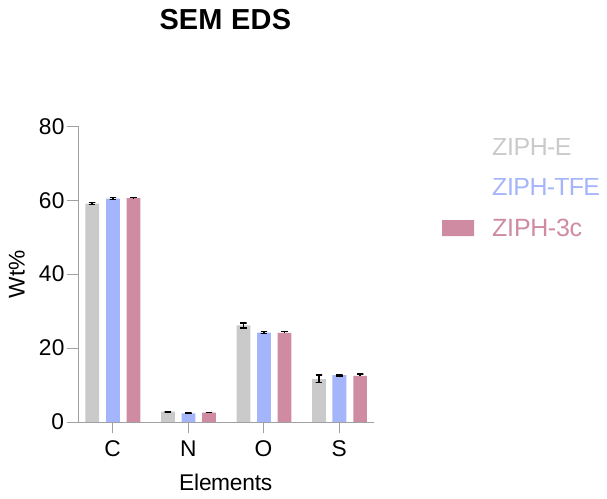
**Supplementary Figure 8.** Comparisons of elemental compositions of thin film samples from Supplementary Fig. 44 using SEM EDX. No significant changes in elemental compositions are observed after solvent annealing which suggests that the decrease in film thickness is due to the collapse of the pores. Gray, ZIPH-E; Blue, ZIPH-TFE; Red, ZIPH-3c.

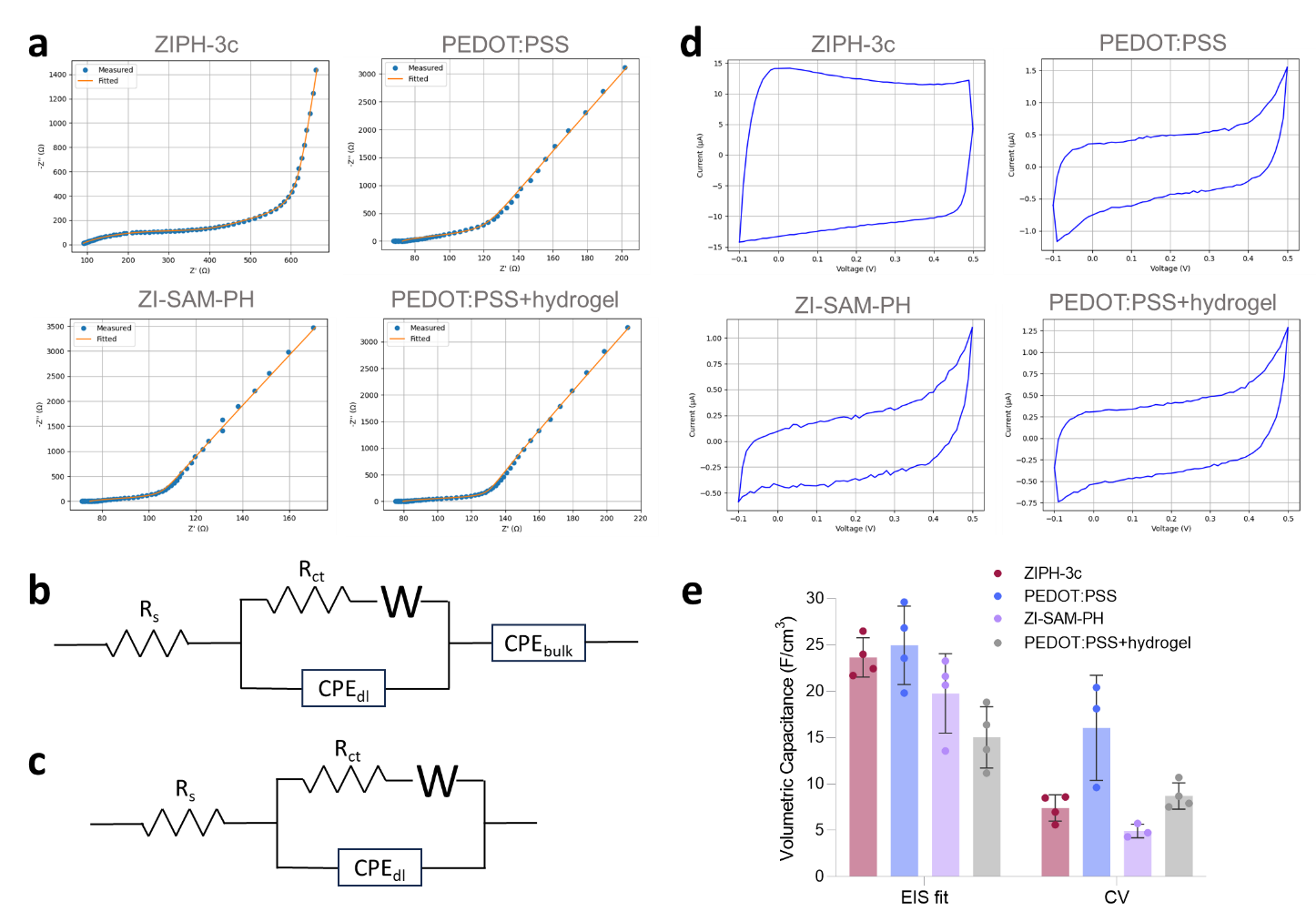

**Supplementary Figure 9.** Comparisons of electrochemical properties. **a**, Nyquist plot and fitting results. **b**, Equivalent circuit model for ZIPH-3c. **c**, Equivalent circuit model for PEDOT:PSS, ZI-SAM-PH, and PEDOT:PSS+hydrogel (hydrogel coating). R_s_ is the solvent resistance, R_ct_ is the charge transfer resistance, W is the finite-space Warburg element, CPE_dl_ is the double-layer constant phase element, and CPE_bulk_ is the bulk constant phase element. **d**, CV curves conducted at 0.1 V/s. **e**, Comparison of capacitances calculated from the EIS fits and the CV curves.

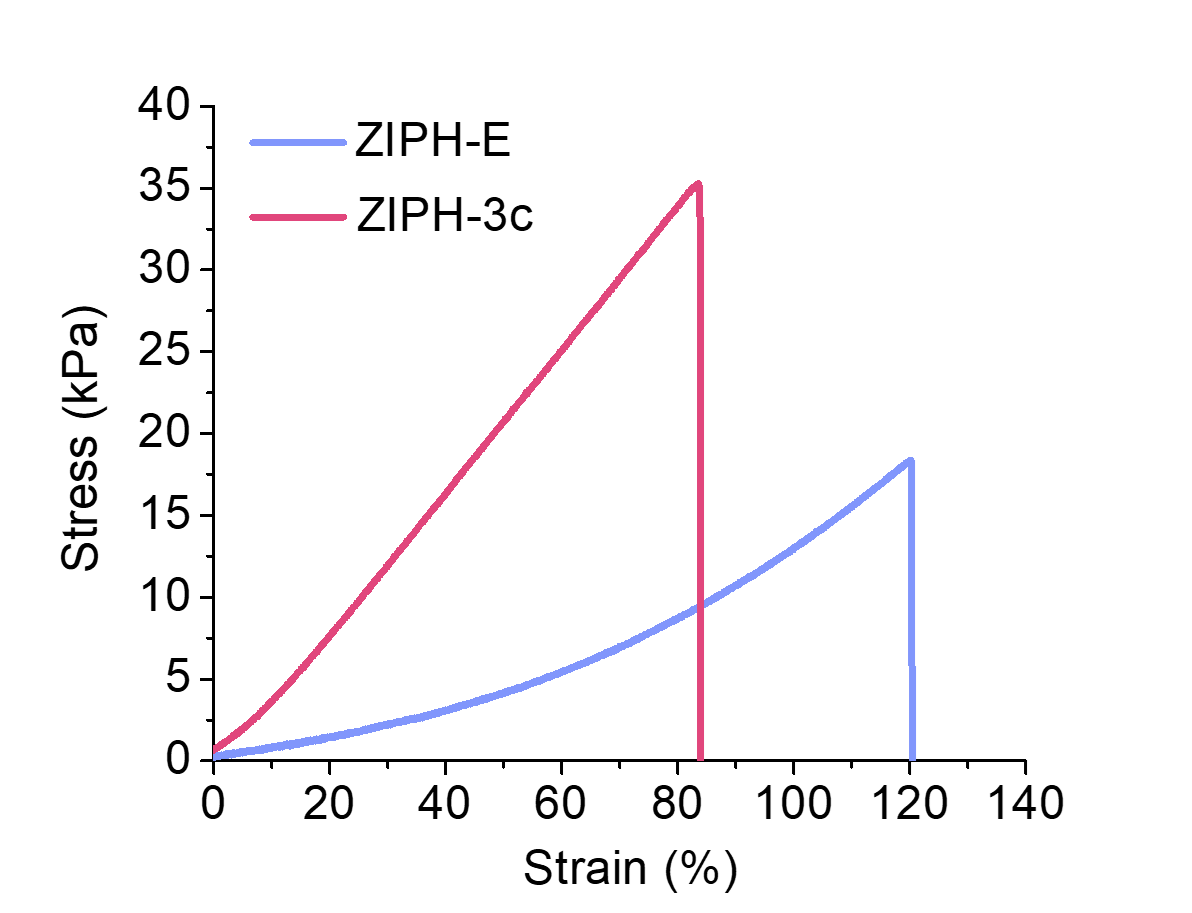

**Supplementary Figure 10.** Tensile curves of ZIPH made with 1.8 mol% crosslinking density and with or without 3 cycles of solvent annealing.

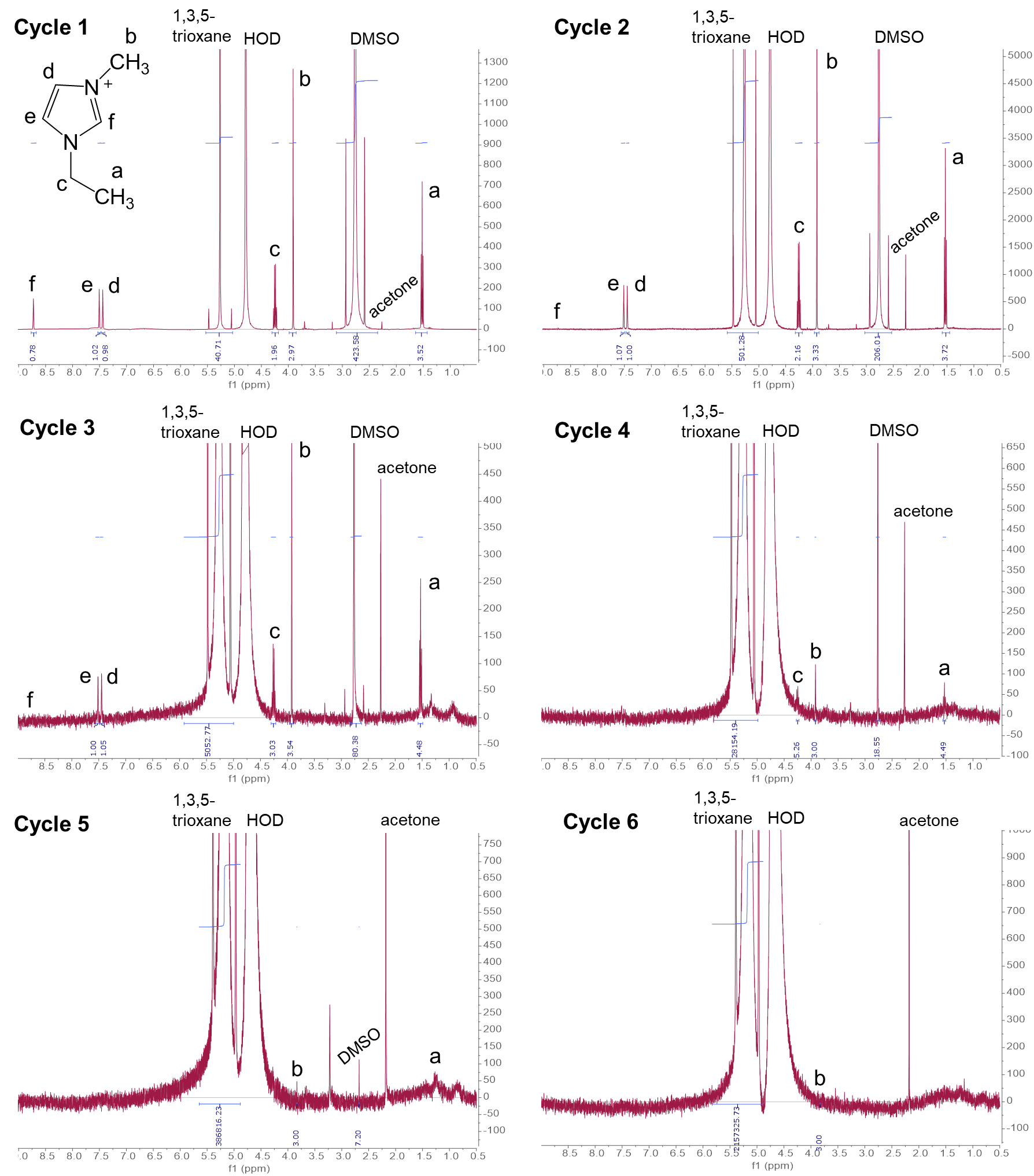

**Supplementary Figure 11.** ^1^H NMR spectra of the supernatants used to calculate the concentrations of EMIM OTf and DMSO in Supplementary Fig. 12 from each wash cycle of PEDOT:PSS hydrogel.

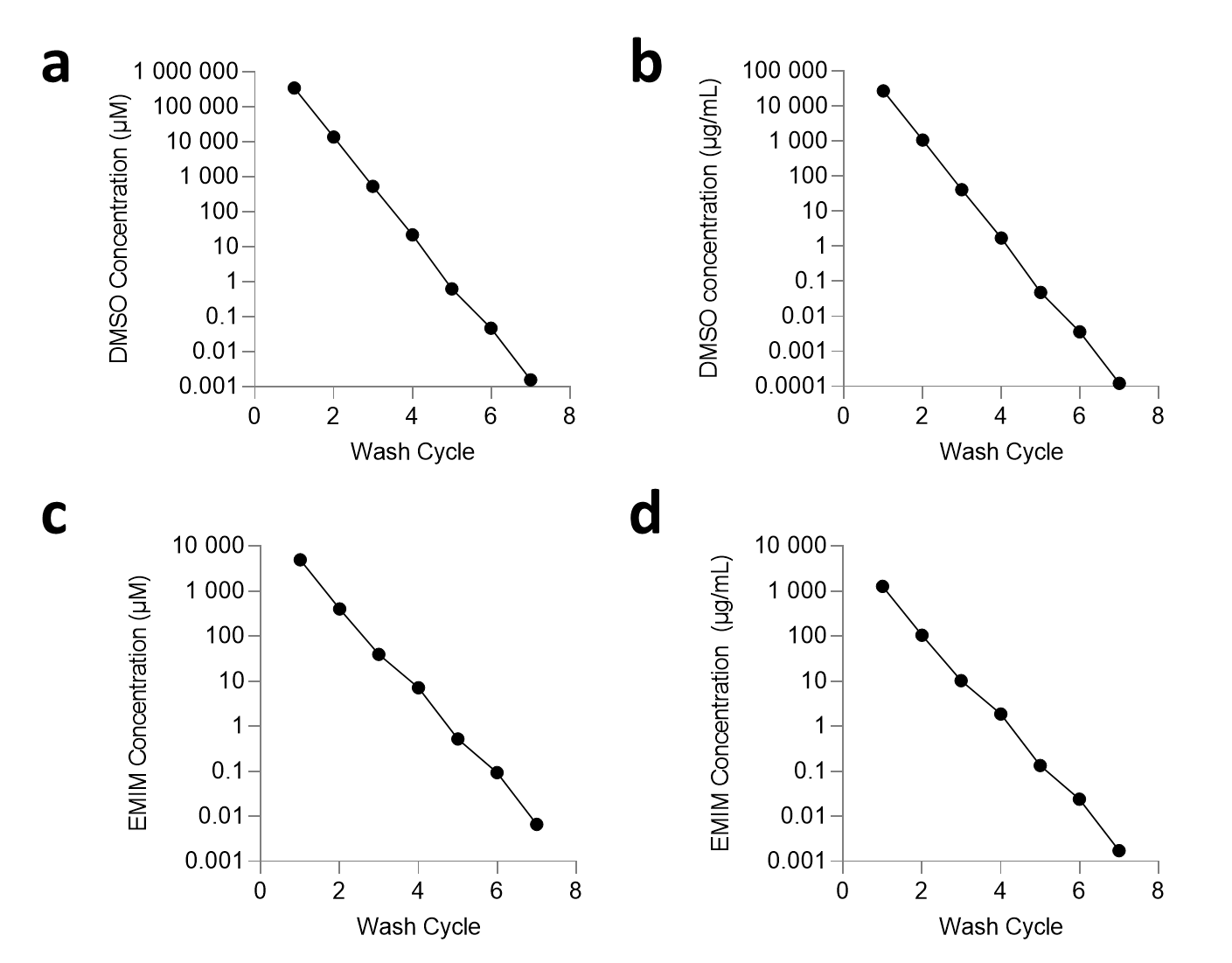

**Supplementary Figure 12.** Quantitative ^1^H NMR of the D_2_O-PBS wash supernatants of the PEDOT:PSS hydrogel used in this work. DMSO concentration (a,b) and EMIM concentration (c,d) after the final wash show that DMSO and EMIM concentrations decrease to nM levels after the final wash. Concentrations at cycle 6 are estimated from the baseline noise of the ^1^H NMR spectra, and concentrations at cycle 7 are extrapolated from the concentrations of previous washes.

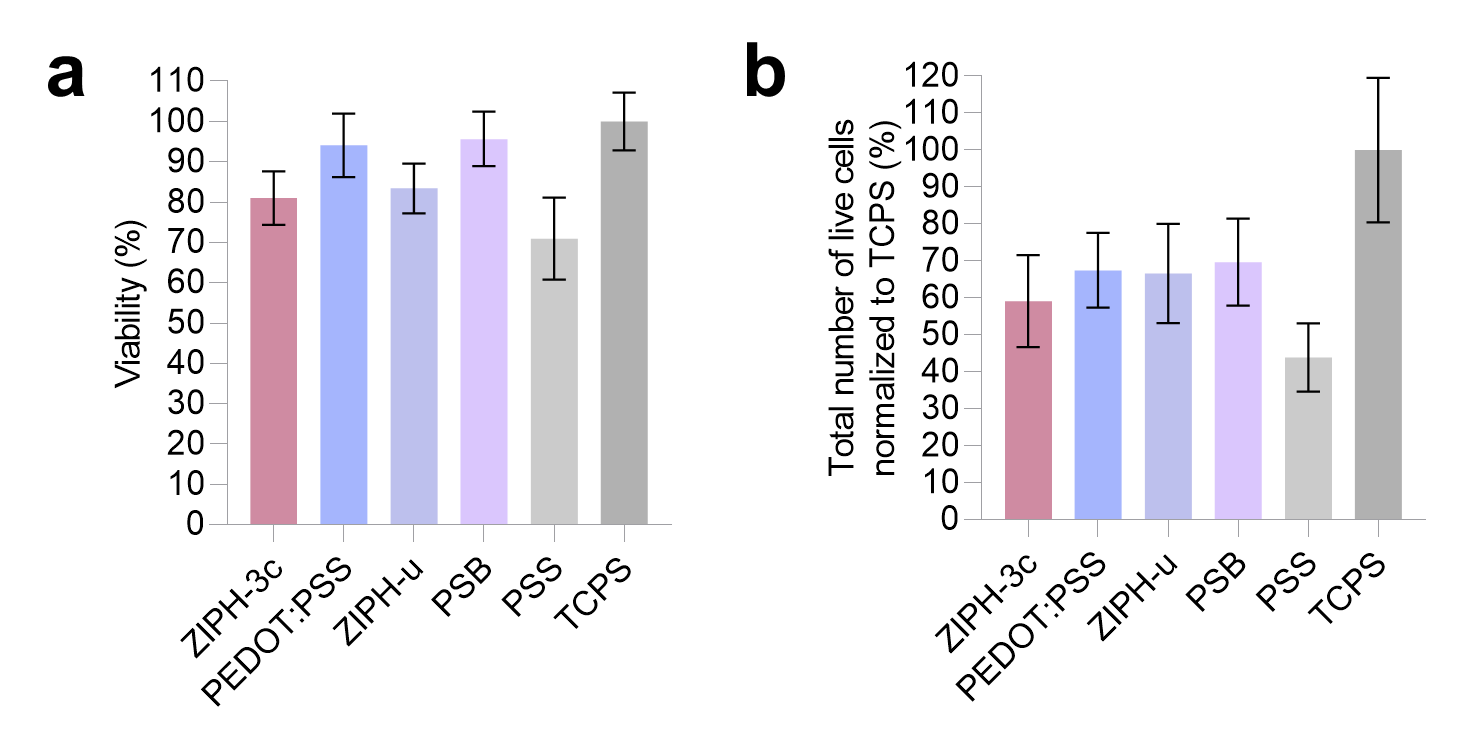
**Supplementary Figure 13.** Cytotoxicity assay of hydrogels. **a,** Percent viability of hydrogels. **b,** Total number of live cells normalized to TCPS. NIH-3T3 cells were cultured directly on hydrogel discs. Calcein-AM (live) and ethidium homodimer-1 (dead) were used for the cytotoxicity assay. TCPS, tissue culture polystyrene.

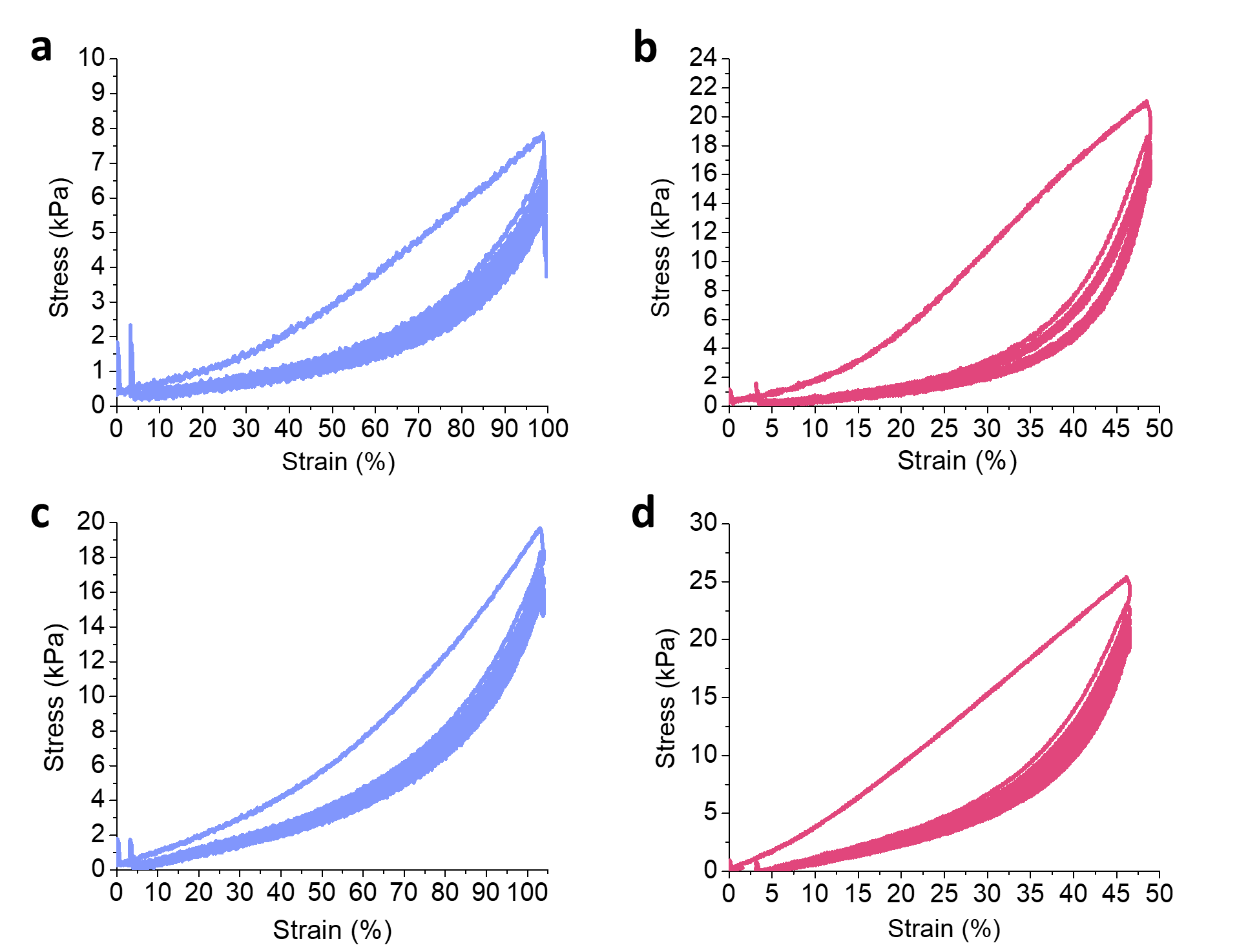

**Supplementary Figure 14.** Cyclic tensile tests for the optimization of mechanical properties of ZIPH. **a,** ZIPH made with 1.0 mol% crosslinking density and no solvent annealing, **b,** ZIPH made with 1.0 mol% crosslinking density and 3 cycles of solvent annealing, **c,** ZIPH made with 1.8 mol% crosslinking density and no solvent annealing, and **d,** ZIPH made with 1.8 mol% crosslinking density and 3 cycles of solvent annealing.

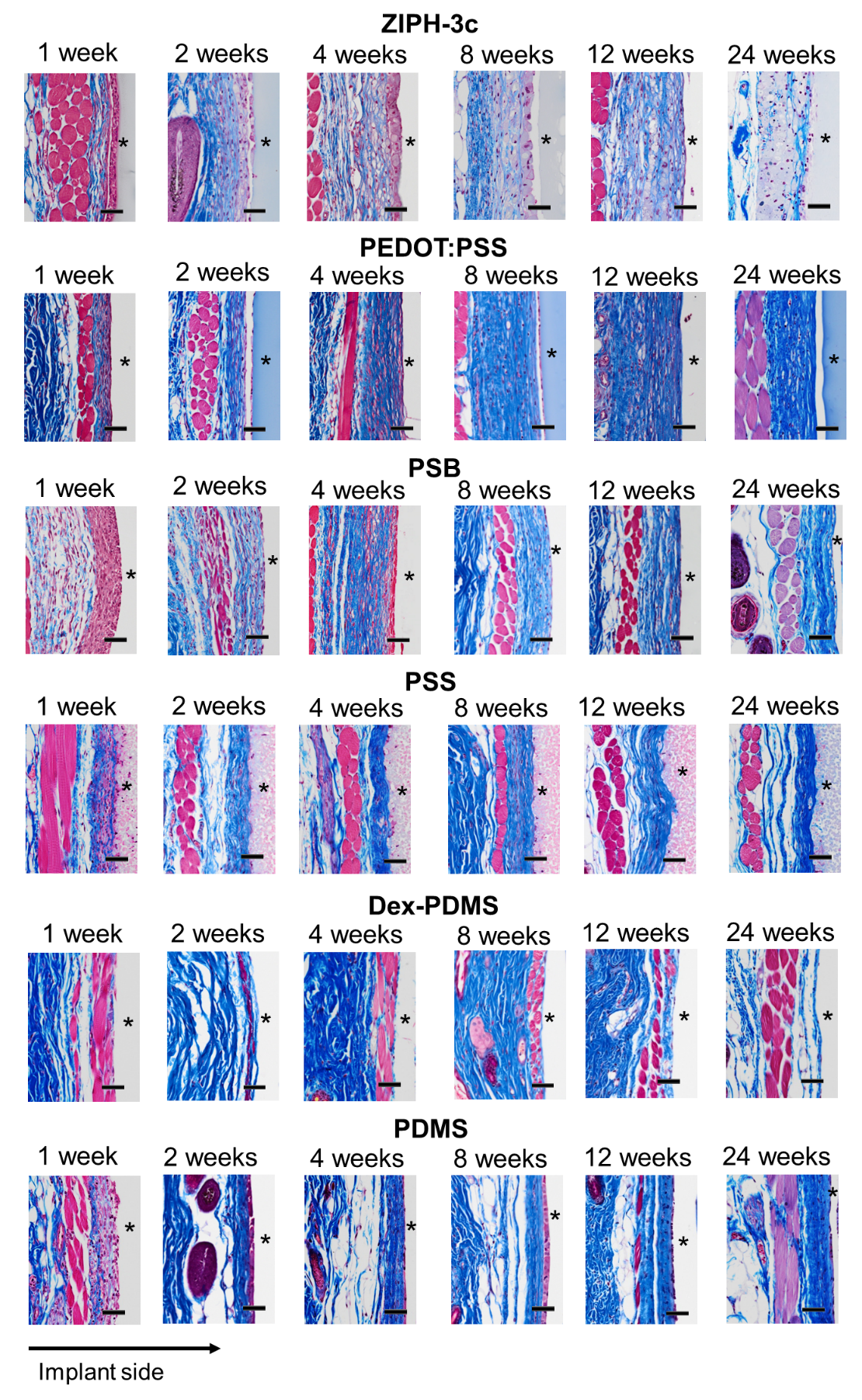
**Supplementary Figure 15.** Representative Masson’s trichrome stained tissues of ZIPH-3c, PEDOT:PSS, PSB, PSS, Dex-PDMS, and PDMS at 1-, 2-, 4-, 8-, 12-, 24-week post-implantation. Locations of implants are marked by asterisks. Scale bar, 50 μm.

**Supplementary Figure 16.** Collagen densities as a function of distance away from the surface of the implants for ZIPH-3c, PEDOT:PSS, and PSB at 1-, 2-, 4-, 8-, 12-, and 24-week timepoints (**a-f**).
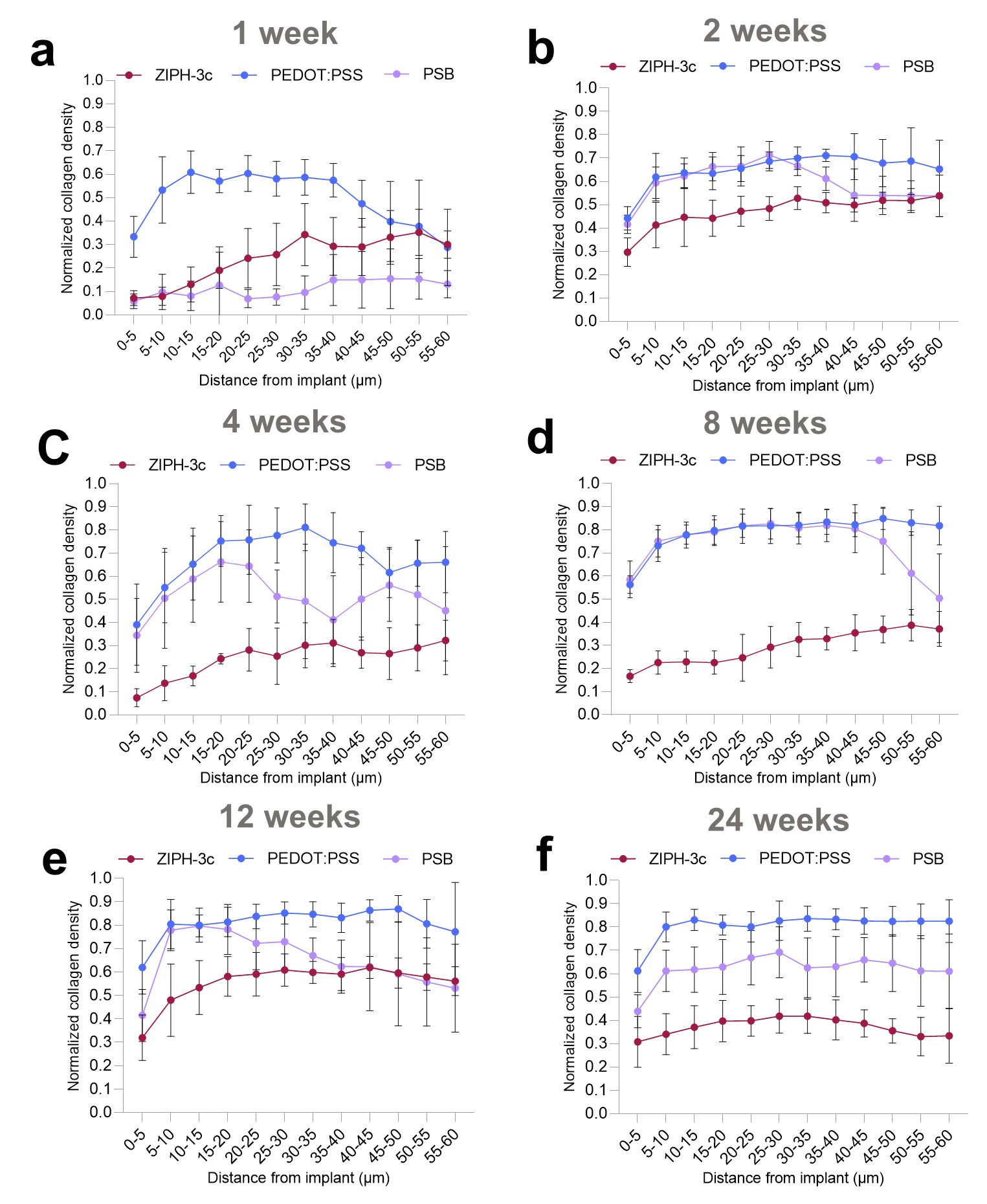

**Supplementary Figure 17.** Collagen densities as a function of distance away from the surface of the implants for PSS, PDMS-Dex at 1-, 2-, 4-, 8-, 12-, and 24-week timepoints (**a-f**).
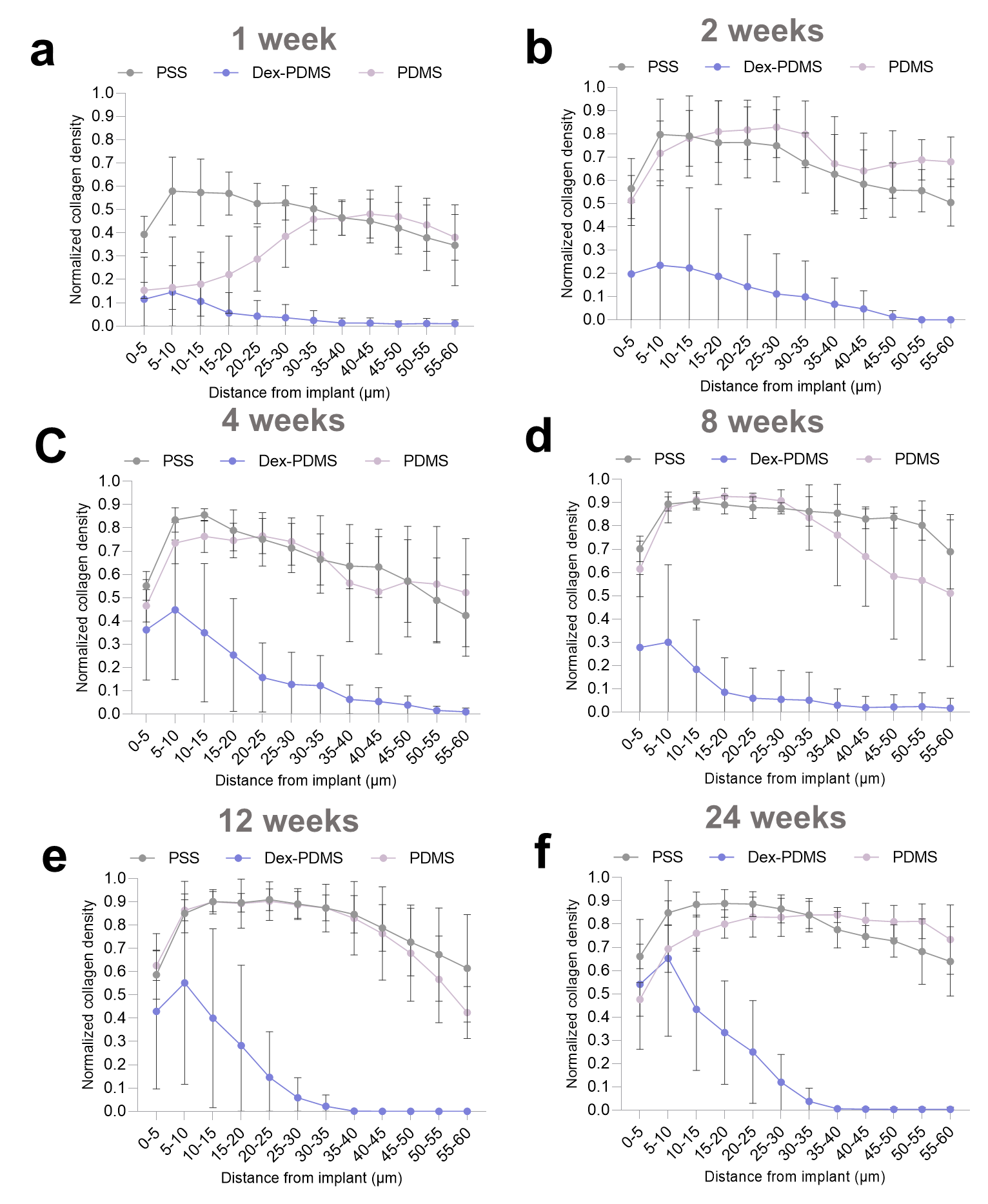

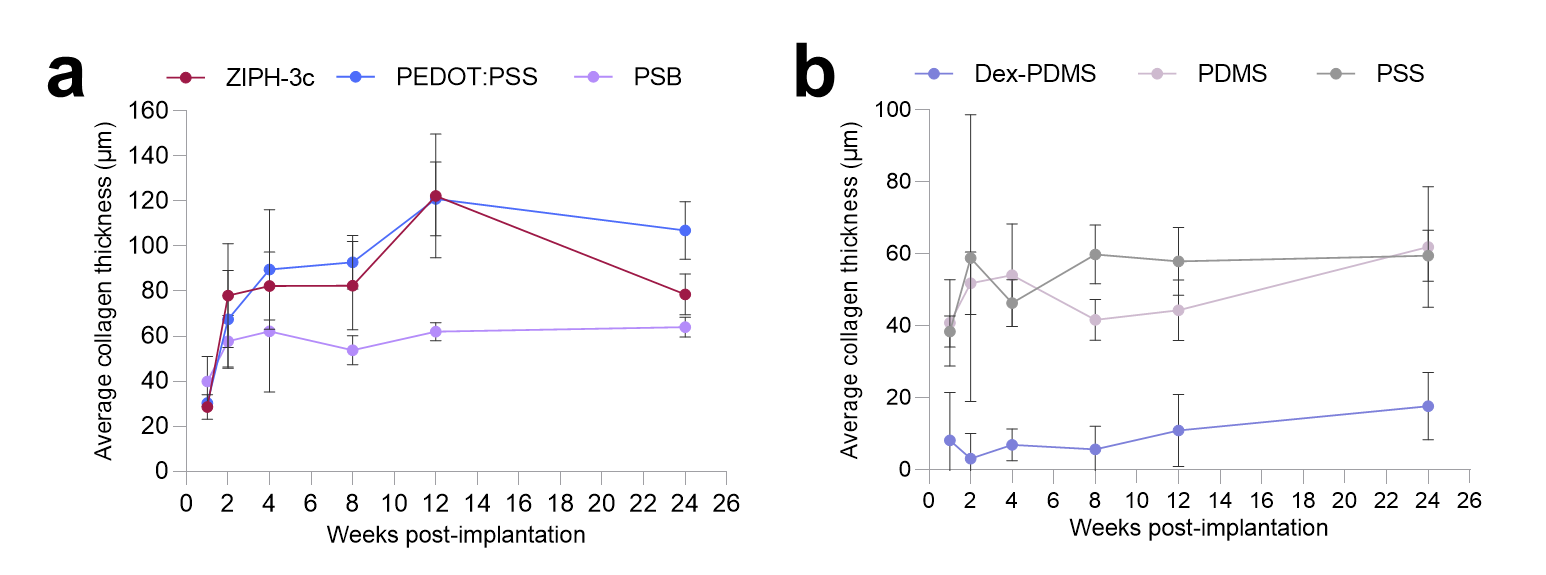

**Supplementary Figure 18.** Fibrotic capsule thicknesses over time for ZIPH-3c, PEDOT:PSS, and PSB (**a**) and Dex-PDMS, PDMS, and PSS (**b**).

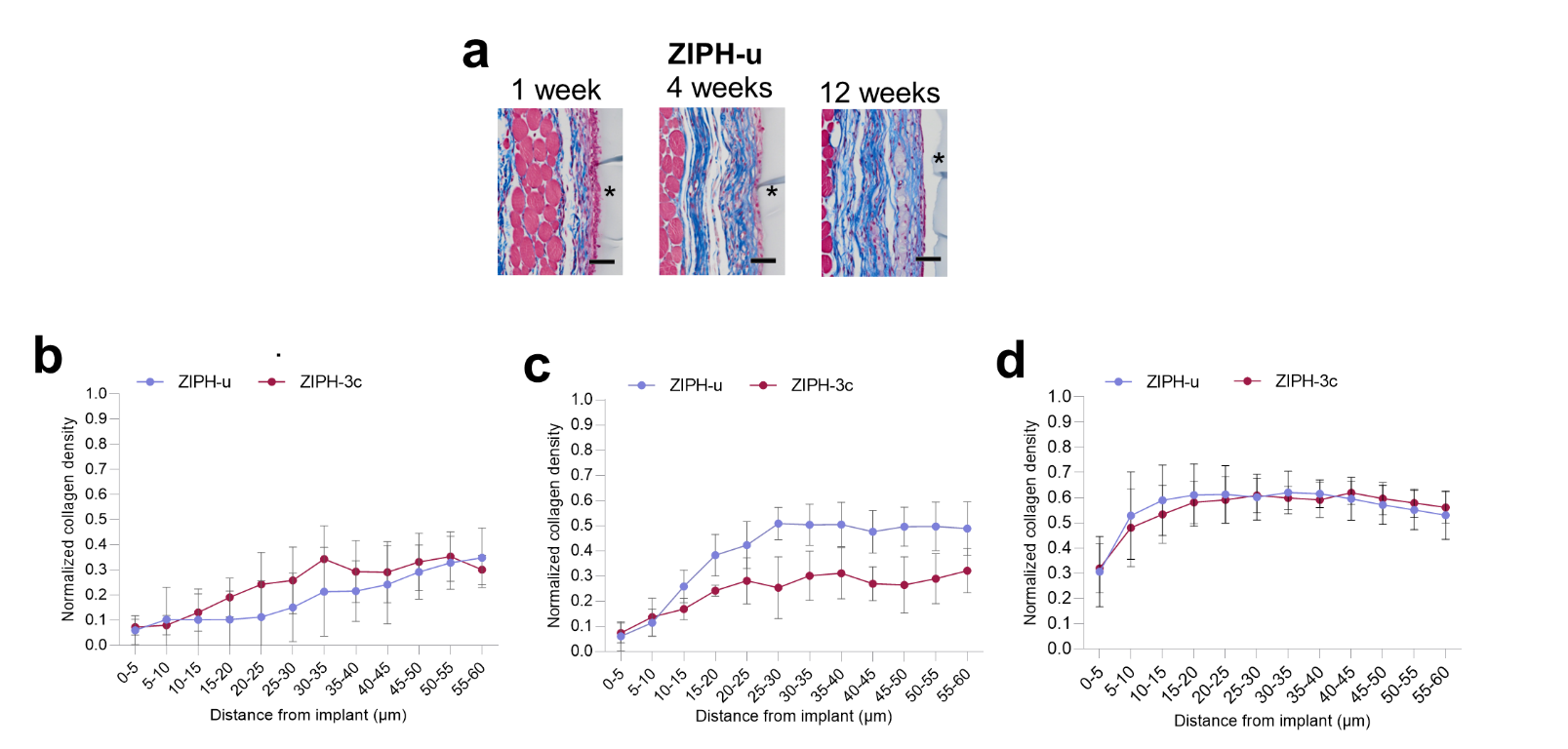

**Supplementary Figure 19.** Collagen densities as a function of distance away from the surface of the implants for ZIPH-u. Representative Masson’s trichrome stained tissues at 1-, 4-, and 12-weeks post-implantation (**a**). Collagen density comparisons of ZIPH-u and ZIPH-3c at 1-, 4-, and 12-week post-implantation (**b-d**).

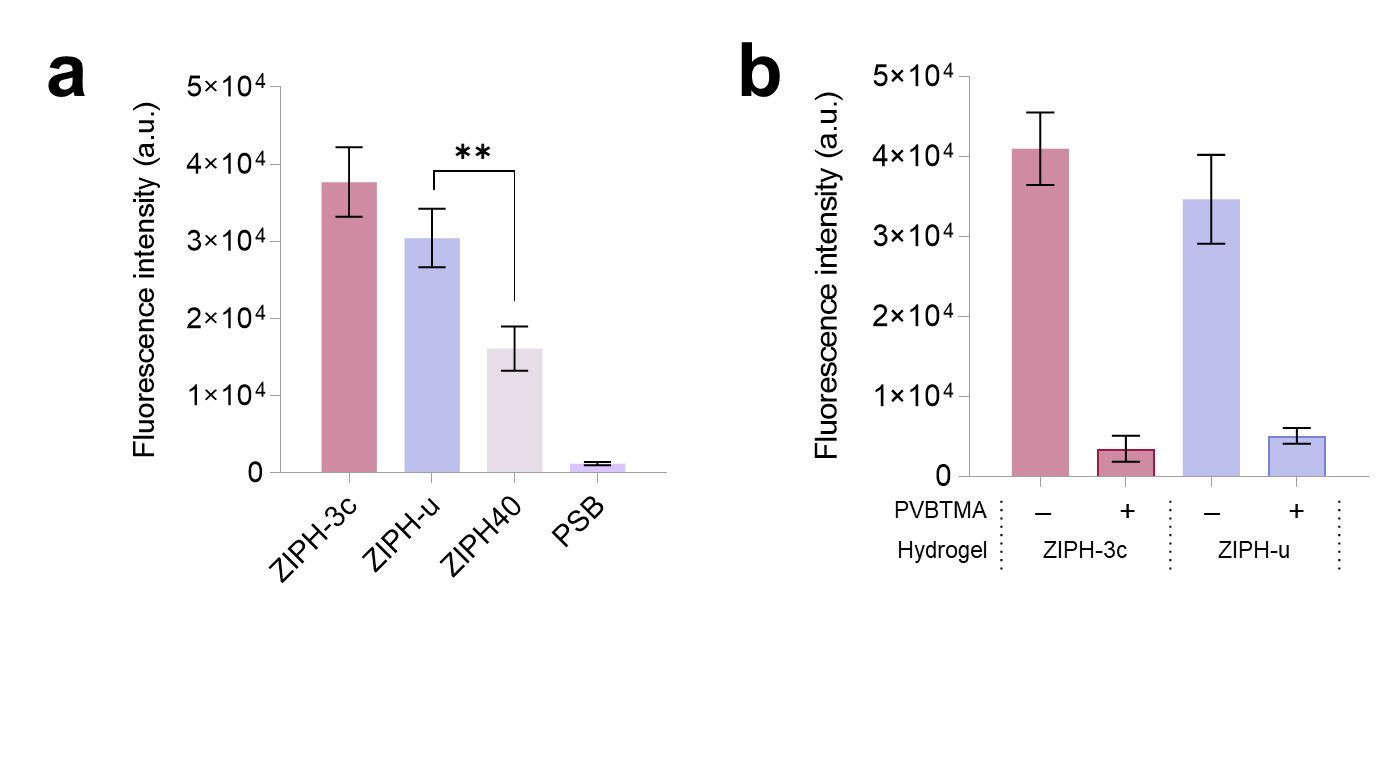
**Supplementary Figure 20. a,** Comparison of positively charged lysozyme protein adsorption on ZIPH-3c, ZIPH-u, ZIPH40 (40% PSB), and PSB. Increasing PSB leads to decrease in lysozyme adsorption. **b,** Comparison of positively charged lysozyme protein adsorption on ZIPH-3c and ZIPH-u non-neutralized or neutralized with PVBTMA. The neutralization of PSS by PBTMA leads to lower protein adsorption. All data were acquired using CBQCA assay.

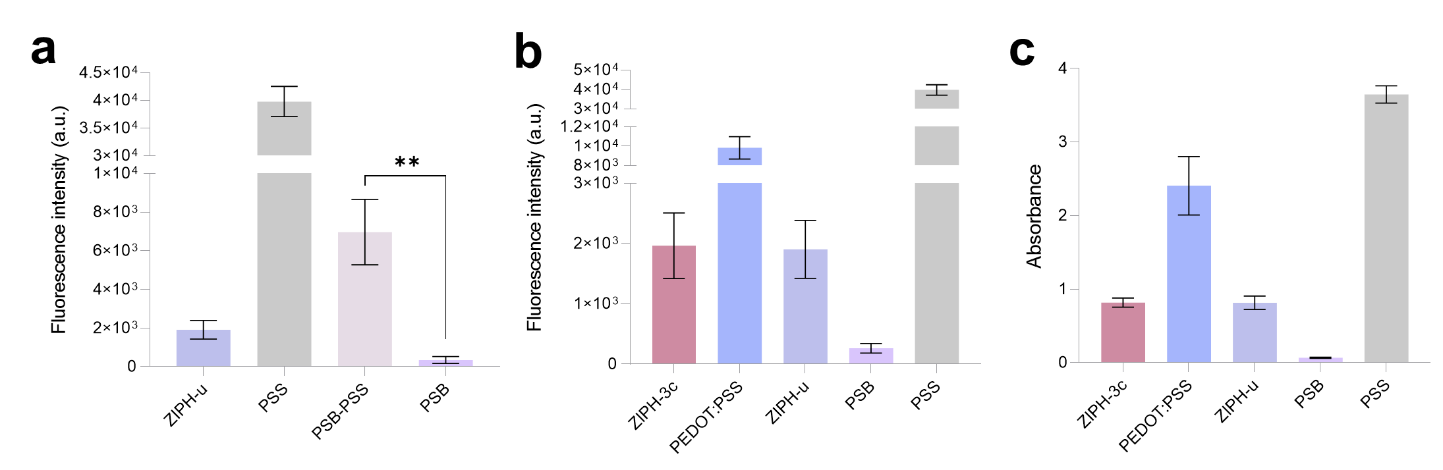
**Supplementary Figure 21.** **a,** Comparison of blood plasma protein adsorption on ZIPH-u, PSS hydrogel, PSB-PSS hydrogel, and PSS determined via CBQCA assay. Adding PSS to PSB hydrogel leads to higher protein adsorption. **b,** Comparison of blood plasma protein adsorption on ZIPH-3c, PEDOT:PSS hydrogel, ZIPH-u, PSB, and PSS determined via CBQCA assay. **c,** Confirmation of the results from (**b**) by using BCA instead of CBQCA.

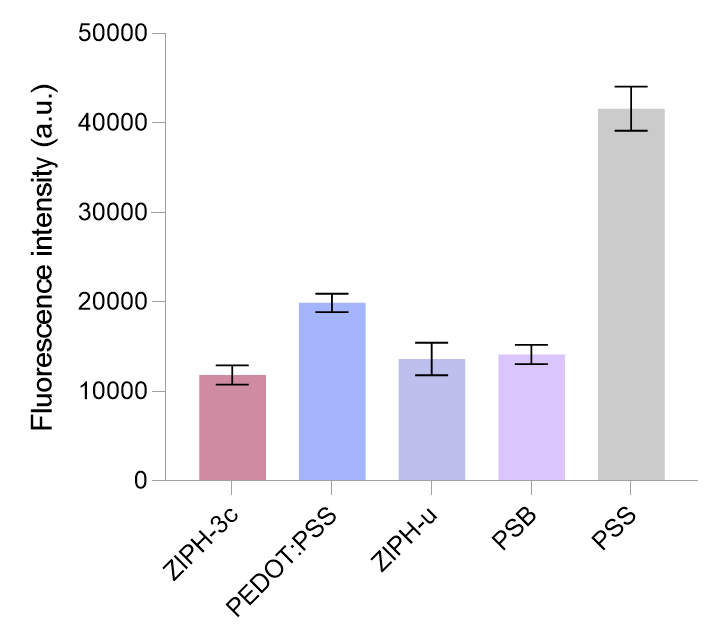
**Supplementary Figure 22.** Comparison of fibrinogen (1 mg/mL) adsorption on ZIPH-3c, PEDOT:PSS hydrogel, ZIPH-u, PSB, and PSS determined via CBQCA assay.

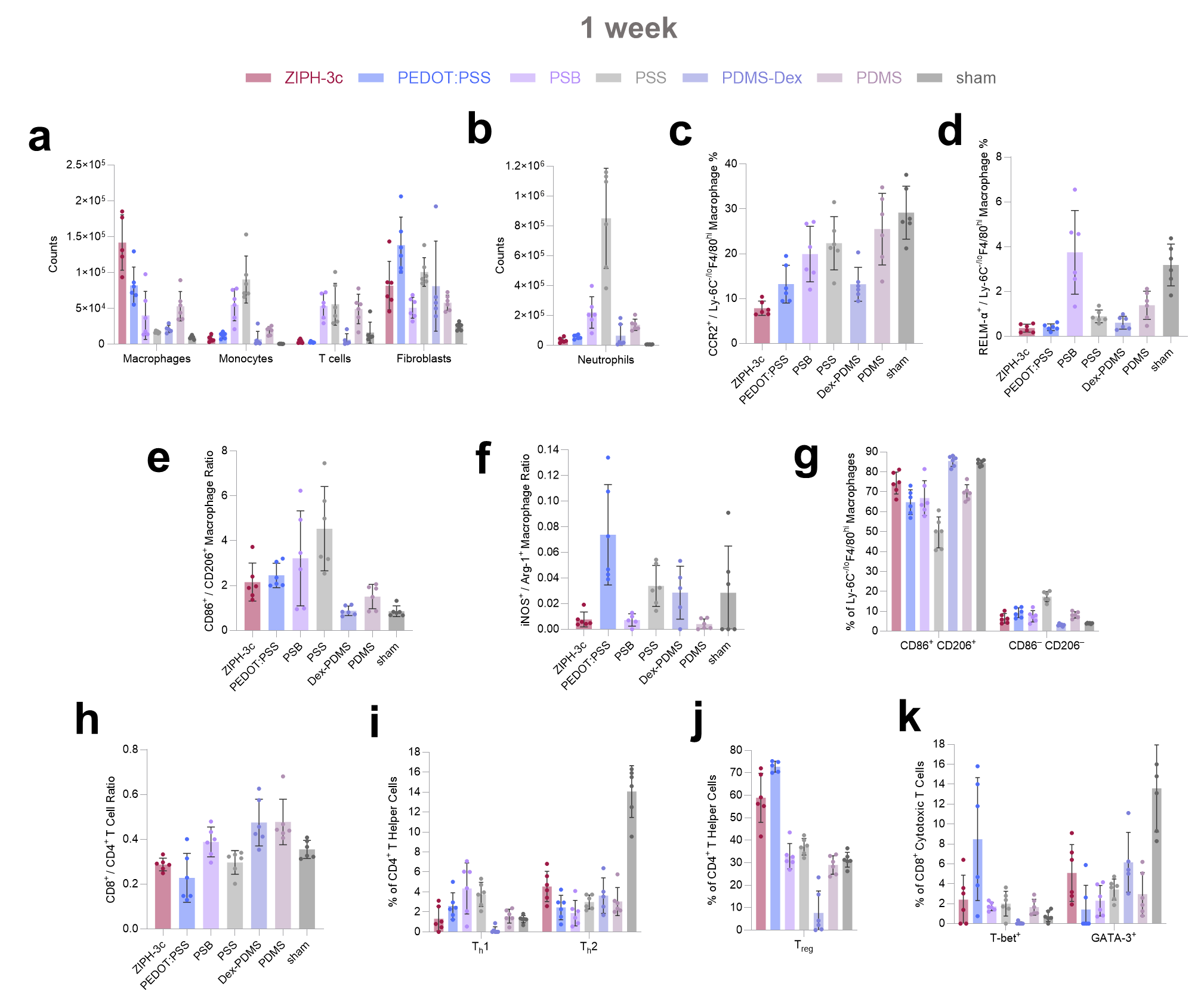

**Supplementary Figure 23.** Flow cytometry data for 1-week post-implantation. **a,** Total Ly-6C ^–/lo^ F4/80^hi^ macrophage, Ly-6C^hi^ F4/80^lo^ monocyte, CD3^+^ T cell, and CD45^–^ Lin^–^ CD140a^+^ fibroblast counts. **b,** Total Ly-6G^+^ neutrophil counts. **c,** Percentage of Ly-6C^–/lo^ F4/80^hi^ macrophages that are CCR2^+^. **d,** Percentage of Ly-6C^–/lo^ F4/80^hi^ macrophages that are RELM-α^+^. **e,** CD86^+^ / CD206^+^ ratios among Ly-6C^–/lo^ F4/80^hi^ macrophages. **f**, iNOS^+^ / Arg-1^+^ ratios among Ly-6C^-/lo^ F4/80^hi^ macrophages. **g**, Percentages of Ly-6C^–/lo^ F4/80^hi^ macrophages that are CD86^+^ CD206^+^ (double positive) and CD86^–^ CD206^–^ (double negative). **h**, CD8^+^ / CD4^+^ ratios among CD3^+^ T cells. **i**, Percentage of CD3^+^ CD4^+^ T helper cells that are T_h_1 (T-bet^+^) and T_h_2 (GATA-3^+^). **j**, Percentage of CD3^+^ T cells that are T_reg_ (FoxP3^+^). **k**, Percentage of CD3^+^ CD8^+^ cytotoxic T cells that are T-bet^+^ and GATA-3^+^.

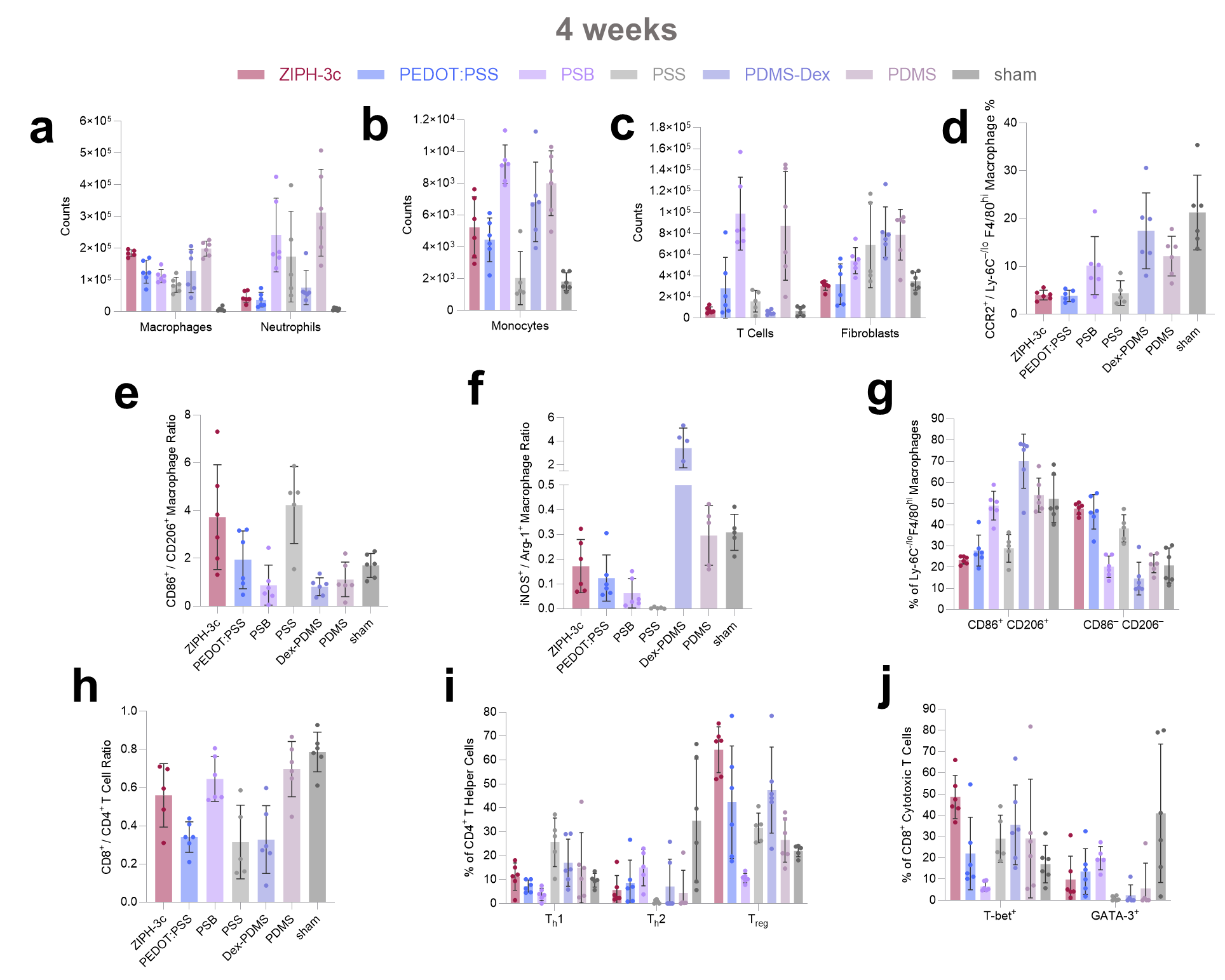

**Supplementary Figure 24.** Flow cytometry data for 4-week post-implantation. **a,** Total Ly-6C ^–/lo^ F4/80^hi^ macrophage and Ly-6G^+^ neutrophil counts. **b**, Total Ly-6C^hi^ F4/80^lo^ monocyte counts. **c**, Total CD3^+^ T cell and CD45^–^ Lin^–^ CD140a^+^ fibroblast counts. **d,** Percentage of Ly-6C^–/lo^ F4/80^hi^ macrophages that are CCR2^+^. **e,** CD86^+^ / CD206^+^ ratios among Ly-6C^–/lo^ F4/80^hi^ macrophages. **f**, iNOS^+^ / Arg-1^+^ ratios among Ly-6C^-/lo^ F4/80^hi^ macrophages. **g**, Percentages of Ly-6C^–/lo^ F4/80^hi^ macrophages that are CD86^+^ CD206^+^ (double positive) and CD86^–^ CD206^–^ (double negative). **h**, CD8^+^ / CD4^+^ ratios among CD3^+^ T cells. **i**, Percentage of CD3^+^ CD4^+^ T helper cells that are T_h_1 (T-bet^+^), T_h_2 (GATA-3^+^), and T_reg_ (FoxP3^+^). **j**, Percentage of CD3^+^ CD8^+^ cytotoxic T cells that are T-bet^+^ and GATA-3^+^.

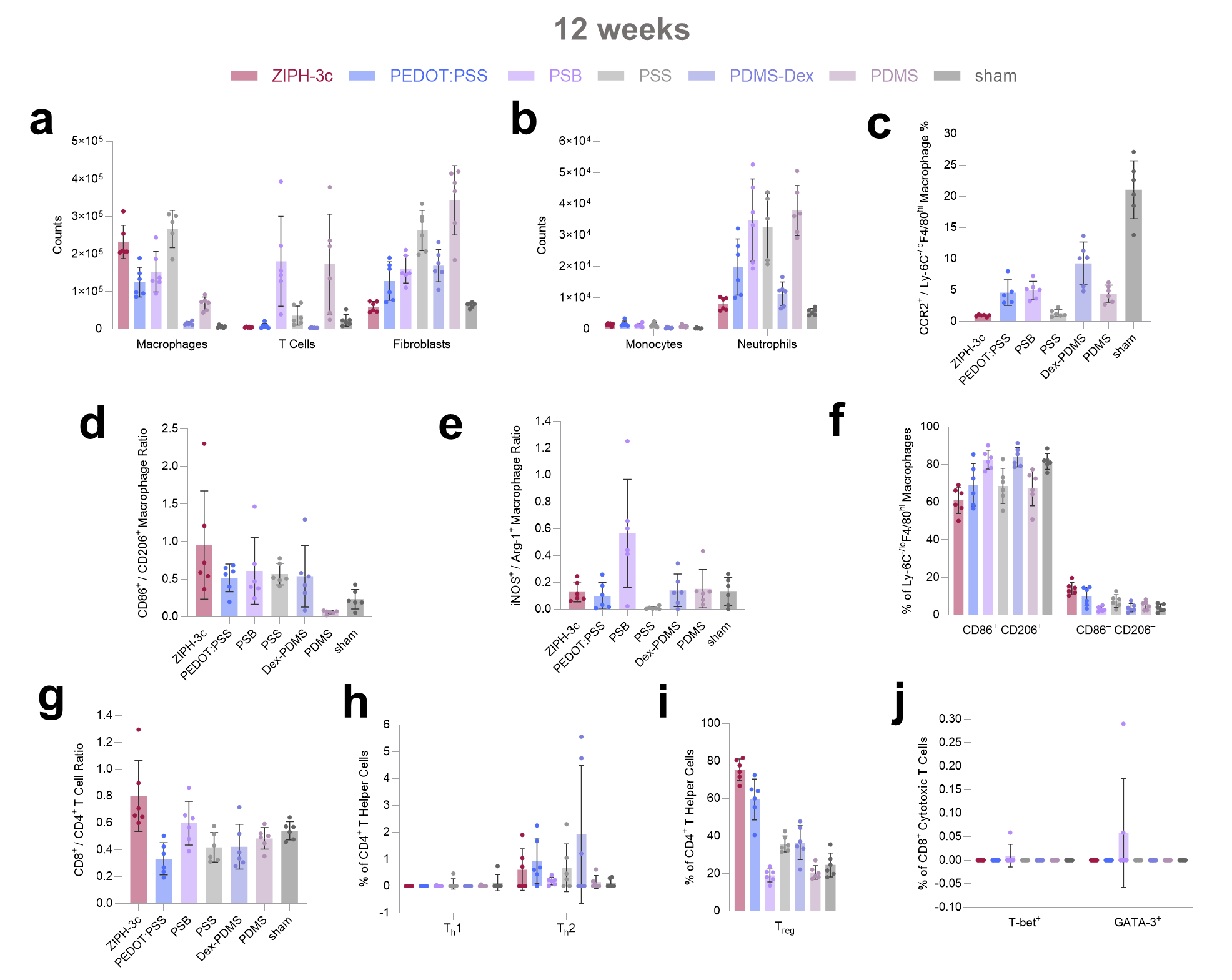

**Supplementary Figure 25.** Flow cytometry data for 12-week post-implantation. **a,** Total Ly-6C ^–/lo^ F4/80^hi^ macrophage, CD3^+^ T cell, and CD45^–^ Lin^–^ CD140a^+^ fibroblast counts. **b,** Total Ly-6C^hi^ F4/80^lo^ monocyte and Ly-6G^+^ neutrophil counts. **c,** Percentage of Ly-6C^–/lo^ F4/80^hi^ macrophages that are CCR2^+^. **d,** CD86^+^ / CD206^+^ ratios among Ly-6C^–/lo^ F4/80^hi^ macrophages. **e**, iNOS^+^ / Arg-1^+^ ratios among Ly-6C^-/lo^ F4/80^hi^ macrophages. **f**, Percentages of Ly-6C^–/lo^ F4/80^hi^ macrophages that are CD86^+^ CD206^+^ (double positive) and CD86^–^ CD206^–^ (double negative). **g**, CD8^+^ / CD4^+^ ratios among CD3^+^ T cells. **h**, Percentage of CD3^+^ CD4^+^ T helper cells that are T_h_1 (T-bet^+^) and T_h_2 (GATA-3^+^). **i**, Percentage of CD3^+^ T cells that are T_reg_ (FoxP3^+^). **j**, Percentage of CD3^+^ CD8^+^ cytotoxic T cells that are T-bet^+^ and GATA-3^+^.

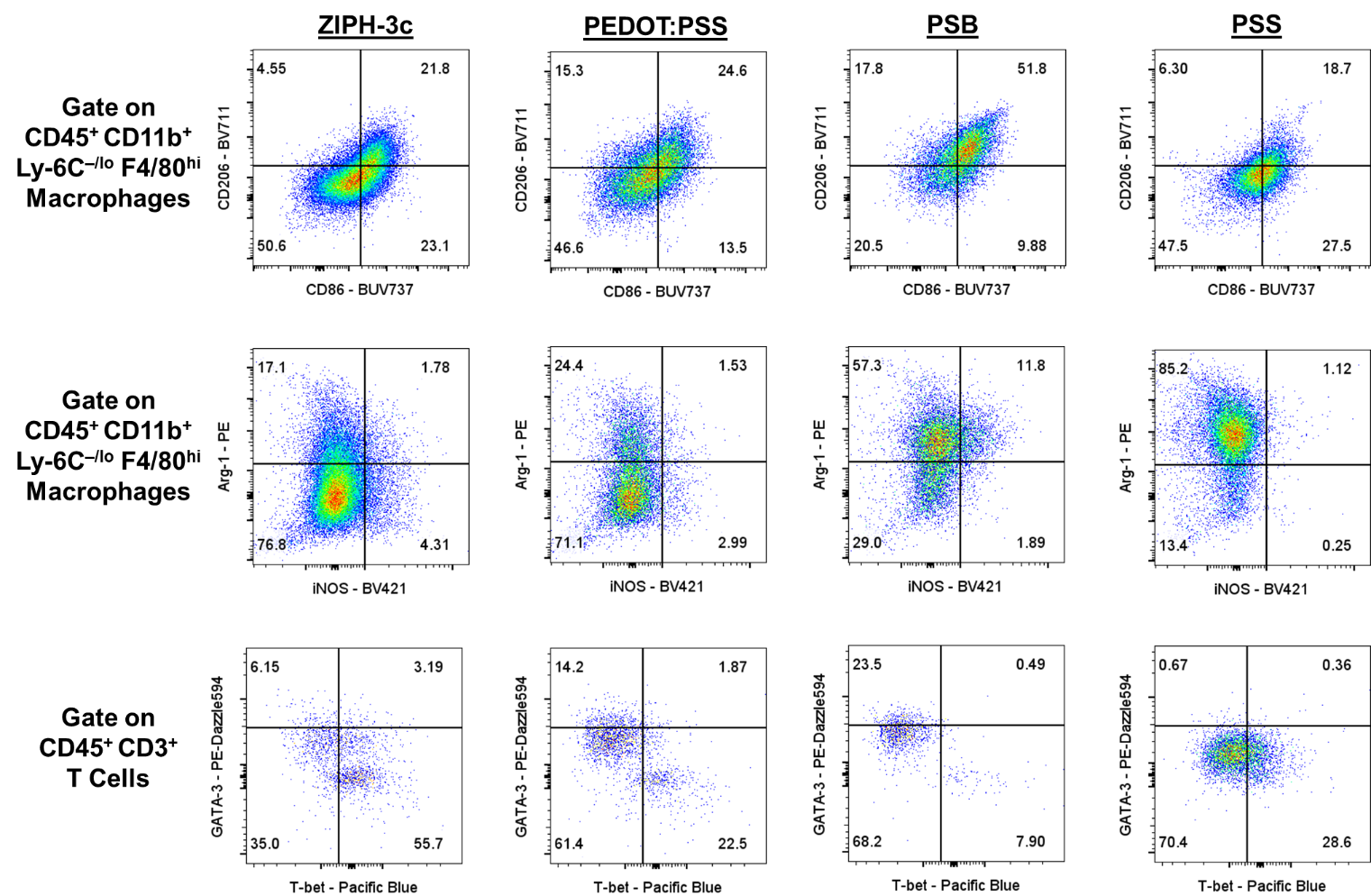

**Supplementary Figure 26.** Representative flow cytometry plots of CD86/CD206, iNOS/Arg-1, and T-bet/GATA-3 cross gates for ZIPH-3c, PEDOT:PSS, PSB, and PSS 4-weeks post-implantation.

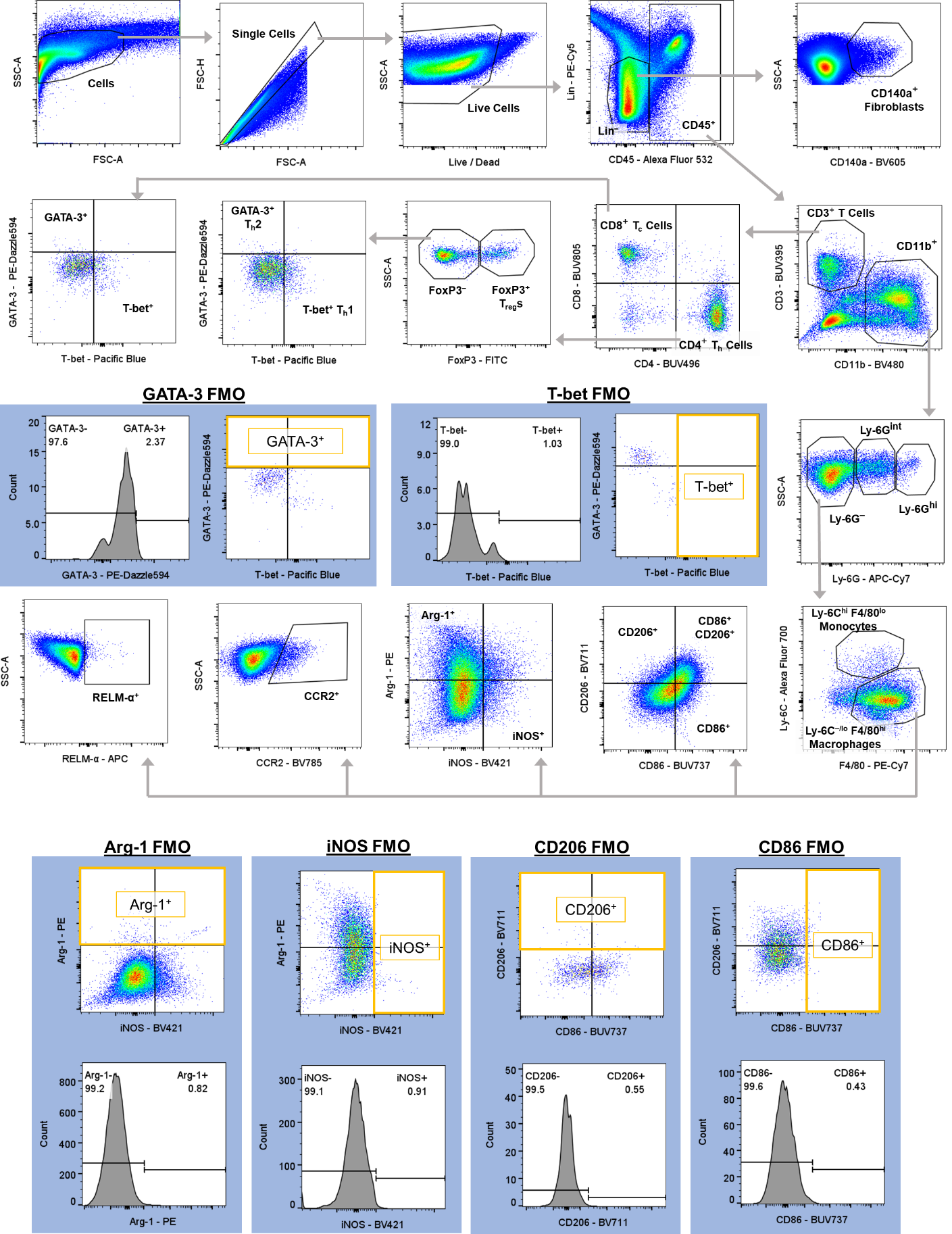
**Supplementary Figure 27.** Flow cytometry gating strategy for all flow cytometry experiments conducted in this work.

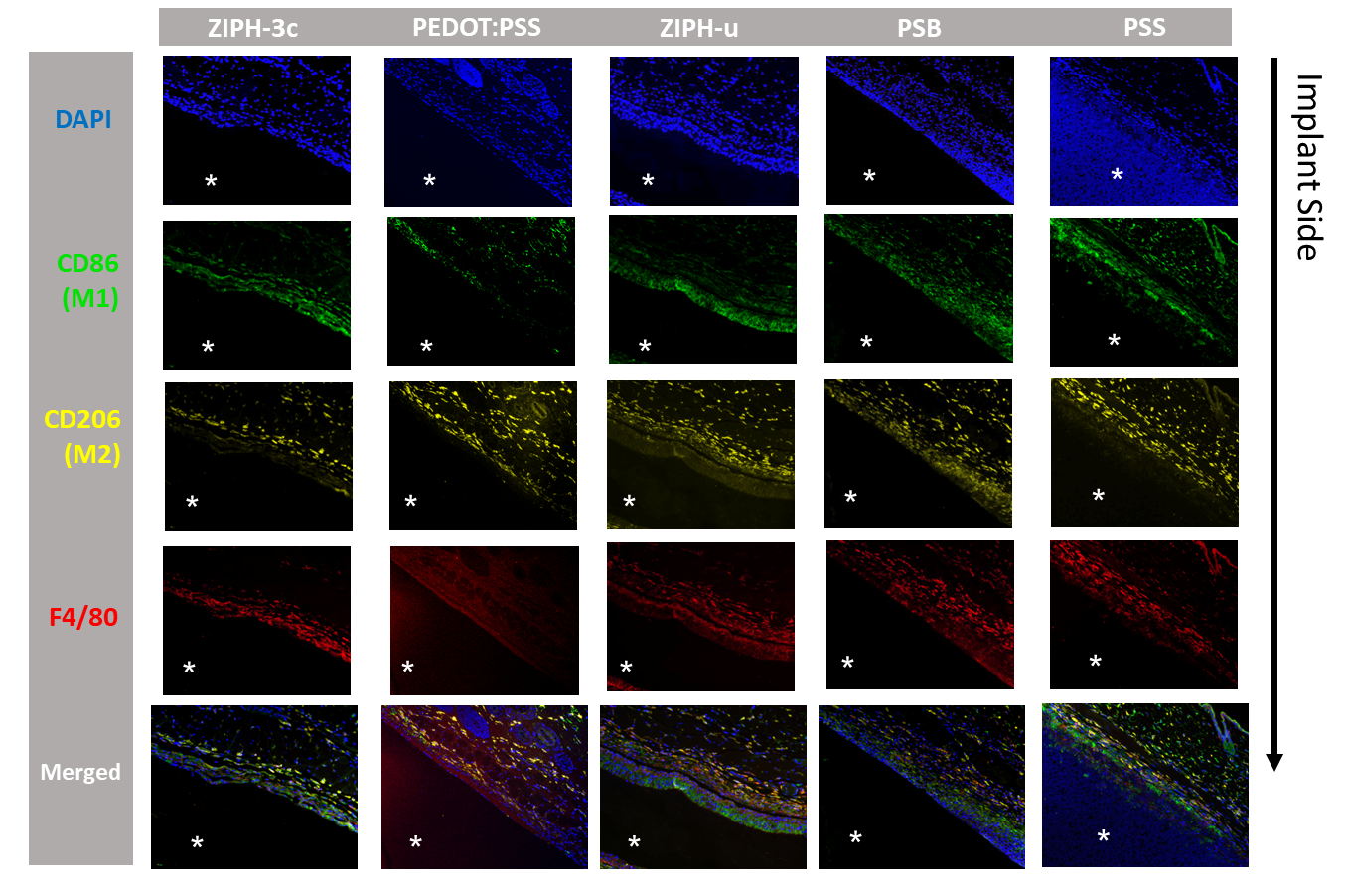
**Supplementary Figure 28.** Immunofluorescence microscopy images of tissue sections stained for markers associated with macrophage polarization states 1-week post-implantation. Implant locations are marked by asterisks

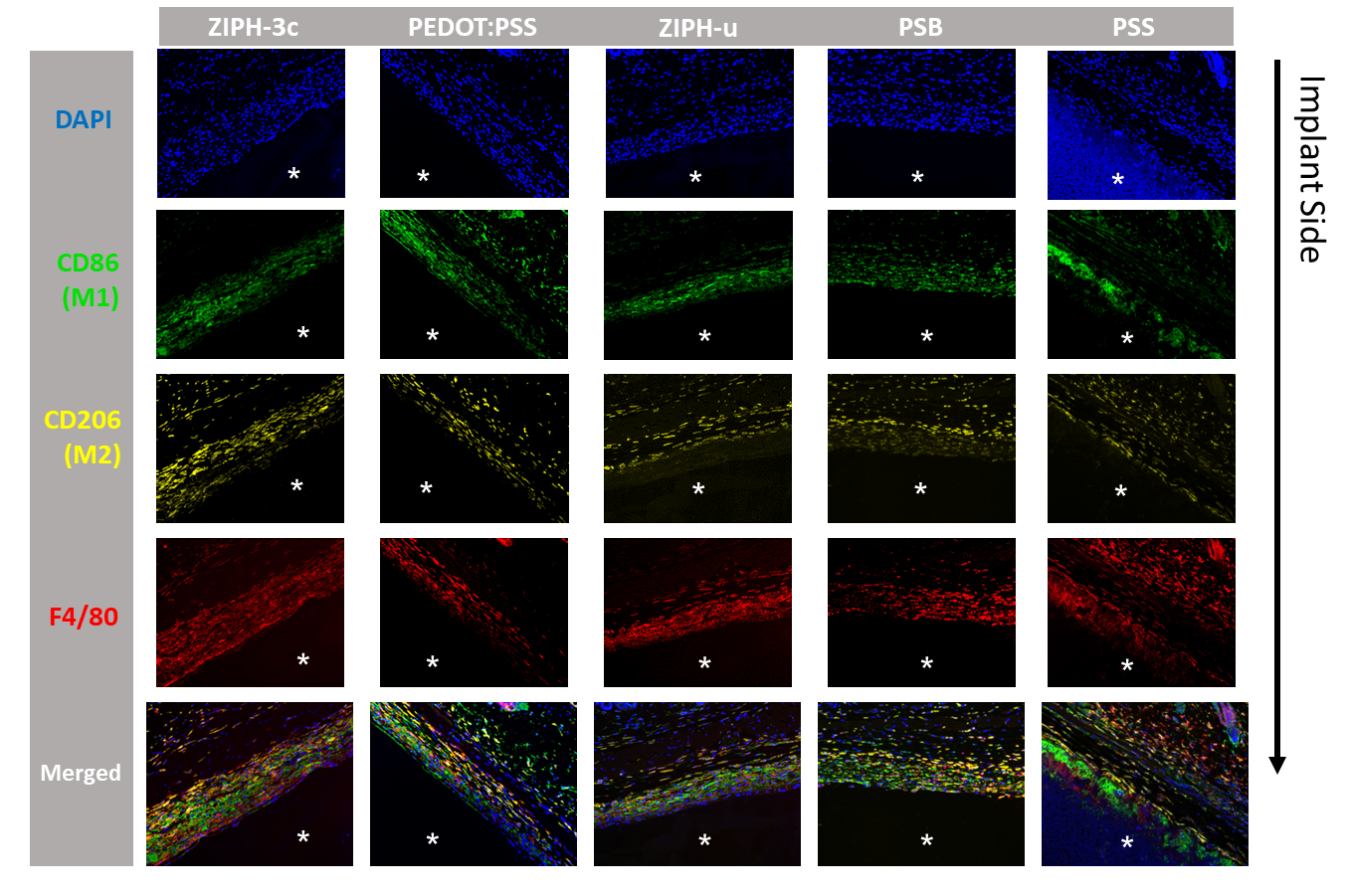
**Supplementary Figure 29.** Immunofluorescence microscopy images of tissue sections stained for markers associated with macrophage polarization states 4-weeks post-implantation. Implant locations are marked by asterisks.

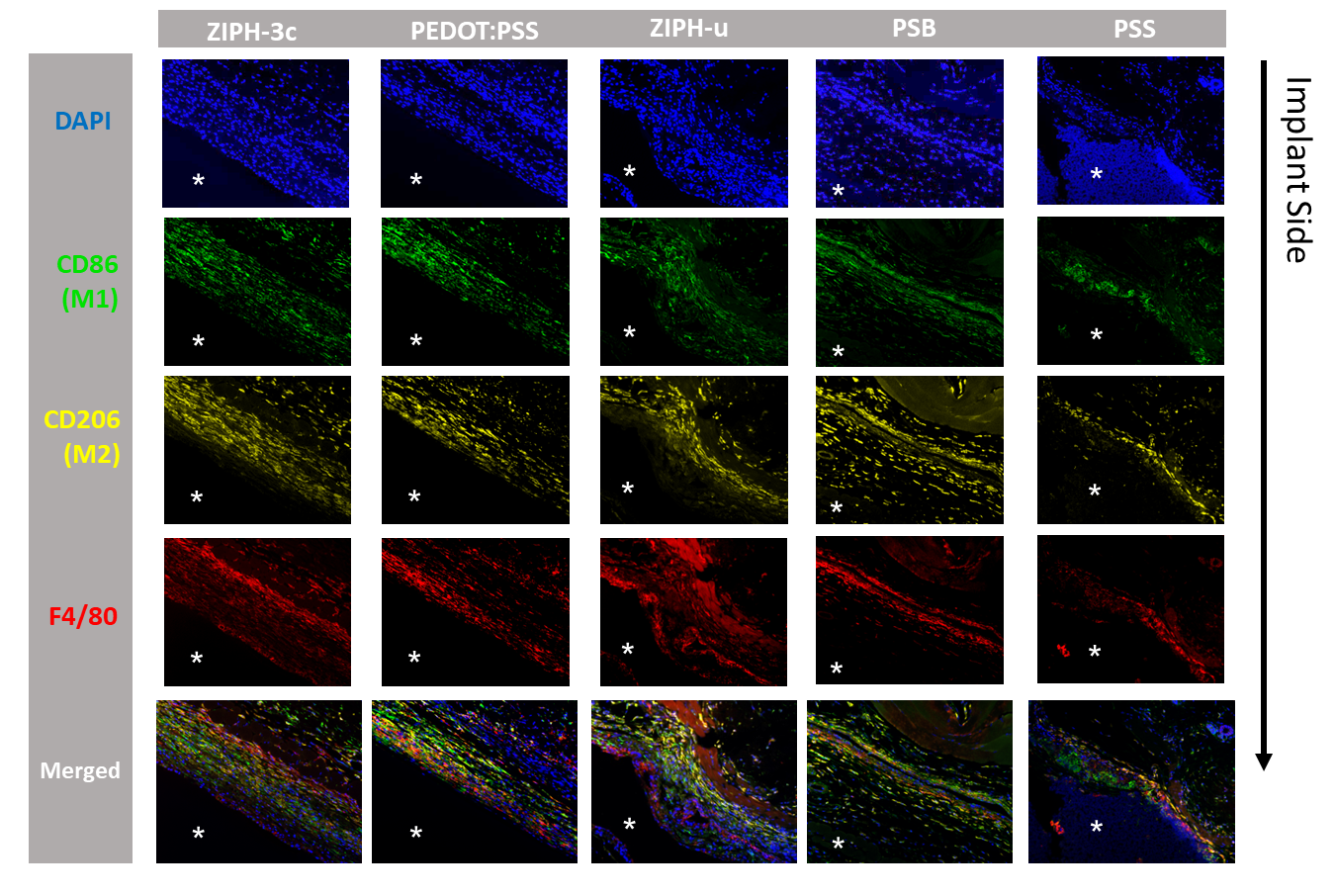
**Supplementary Figure 30.** Immunofluorescence microscopy images of tissue sections stained for markers associated with macrophage polarization states 12-weeks post-implantation. Implant locations are marked by asterisks

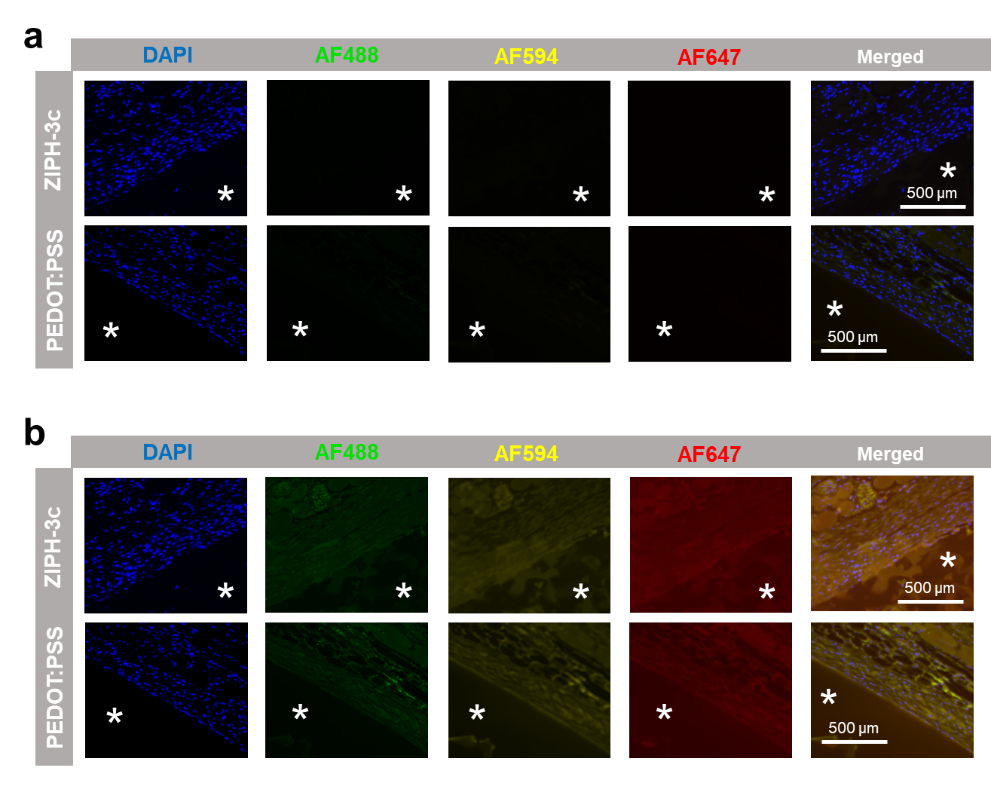

**Supplementary Figure 31.** As immunofluorescence background controls, tissue sections were stained using the same procedures as the experimental groups except the primary antibody step was omitted. Images kept at the original brightness (**a**) and non-DAPI channel images with pixel brightnesses increased over 10-fold (**b**) show that the background features bear no resemblance to the positive features observed from the experimental groups. Implant locations are marked with asterisks.

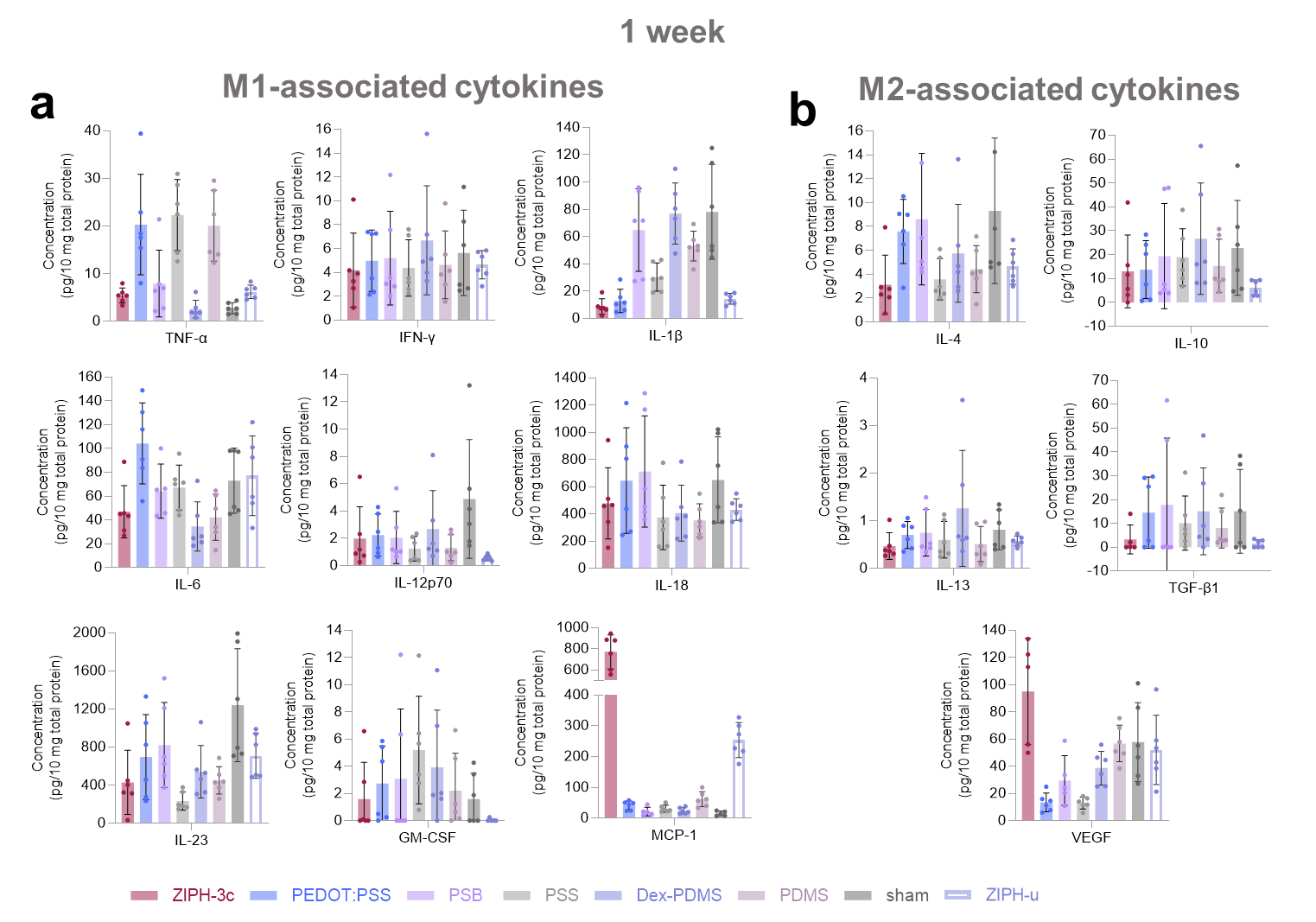

**Supplementary Figure 32.** Cytokine concentrations determined via LegendPlex assay divided into **a,** M1-associated cytokines and **b,** M2-associated cytokines 1-week post-implantation.

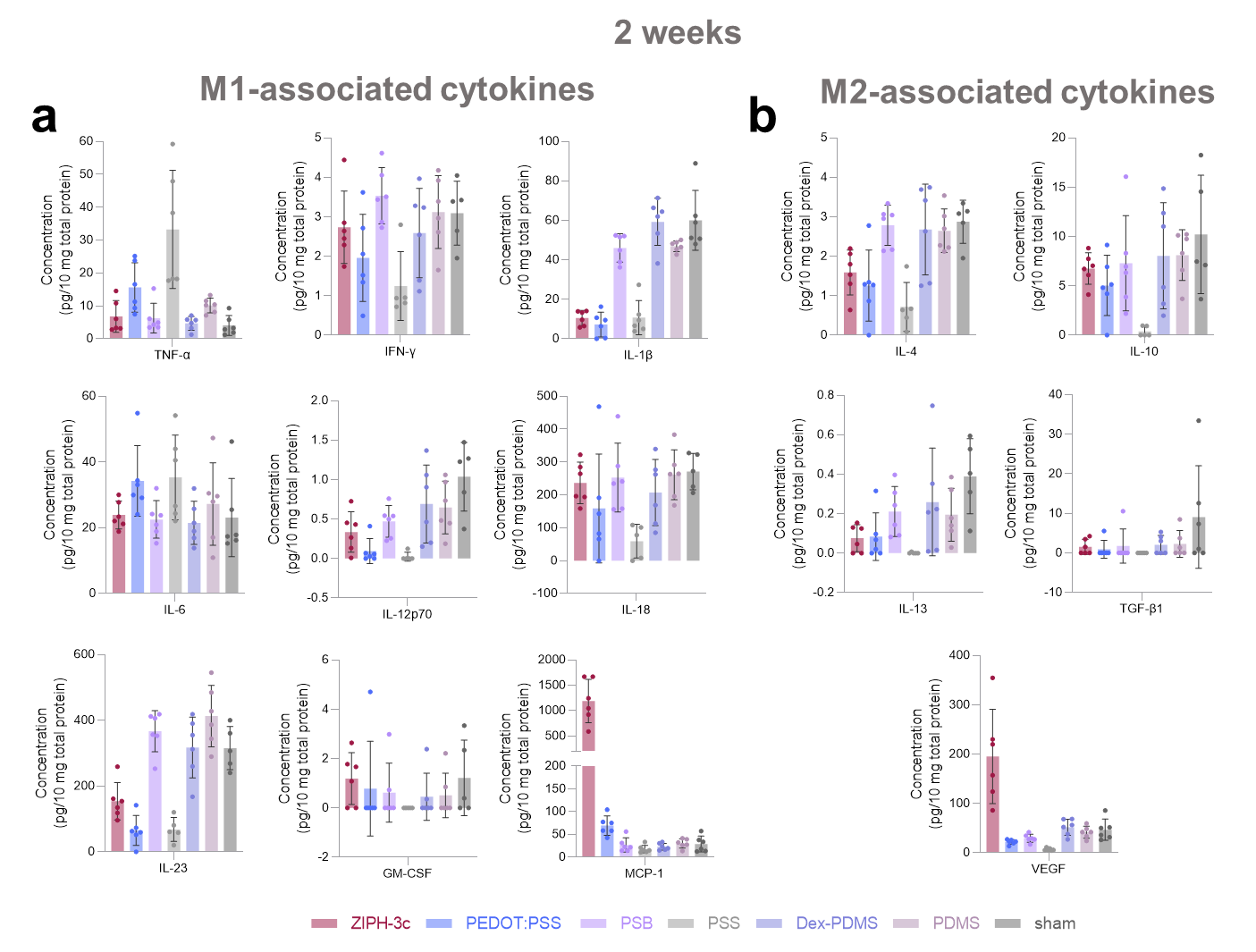
**Supplementary Figure 33.** Cytokine concentrations determined via LegendPlex assay divided into **a,** M1-associated cytokines and **b,** M2-associated cytokines 2-weeks post-implantation.

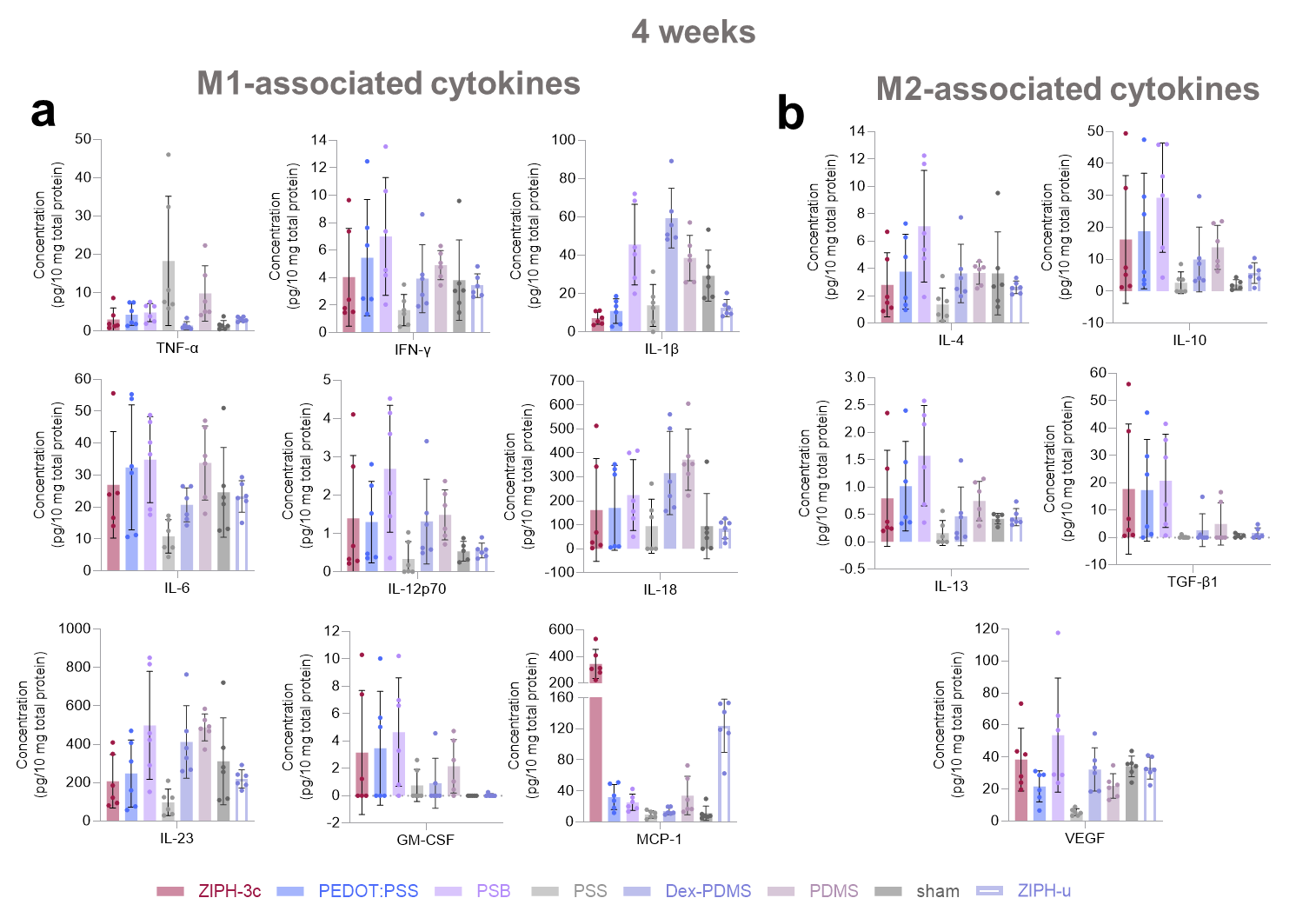
**Supplementary Figure 34.** Cytokine concentrations determined via LegendPlex assay divided into **a,** M1-associated cytokines and **b,** M2-associated cytokines 4-week post-implantation.

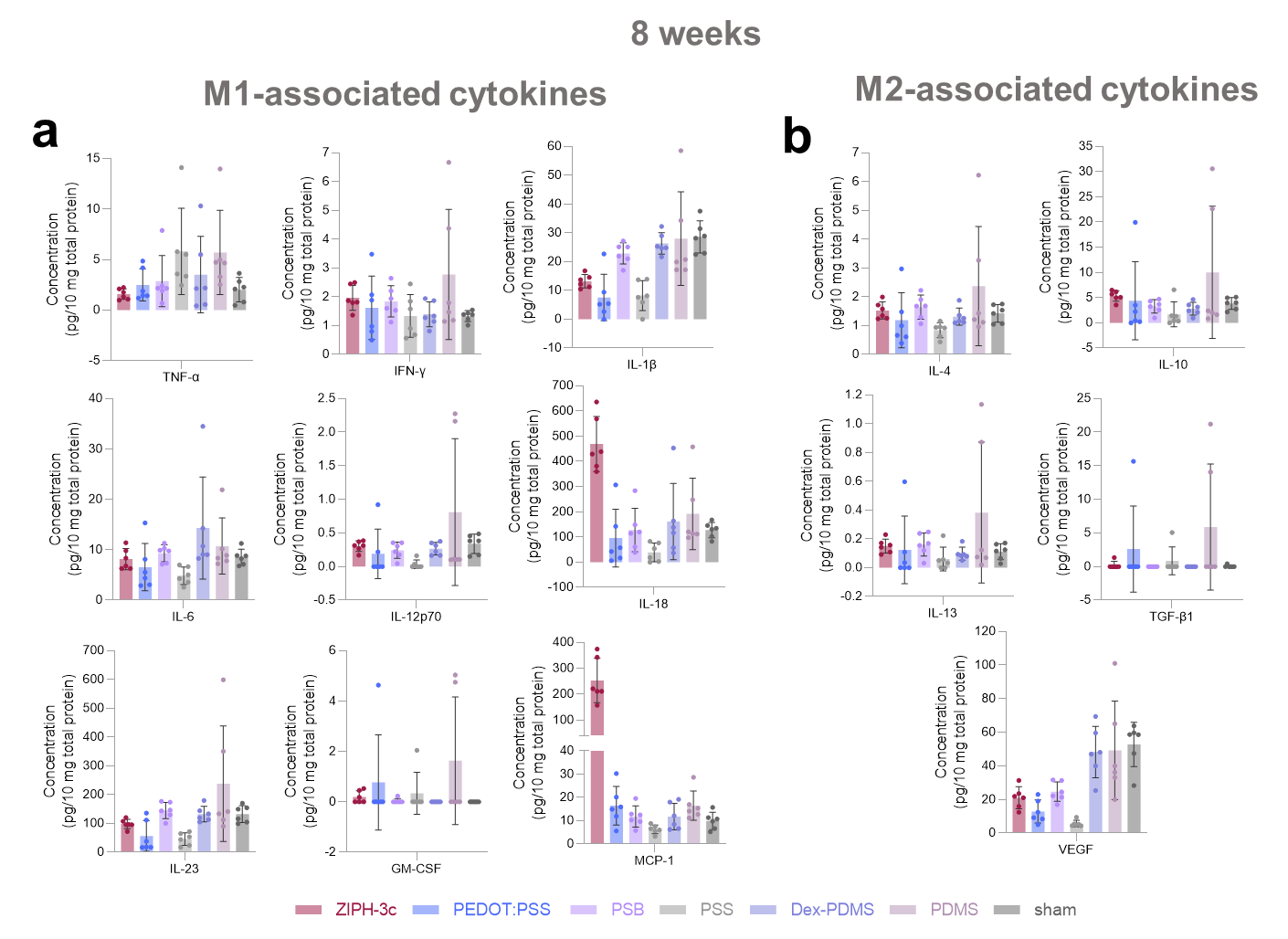
**Supplementary Figure 35.** Cytokine concentrations determined via LegendPlex assay divided into **a,** M1-associated cytokines and **b,** M2-associated cytokines 8-week post-implantation.

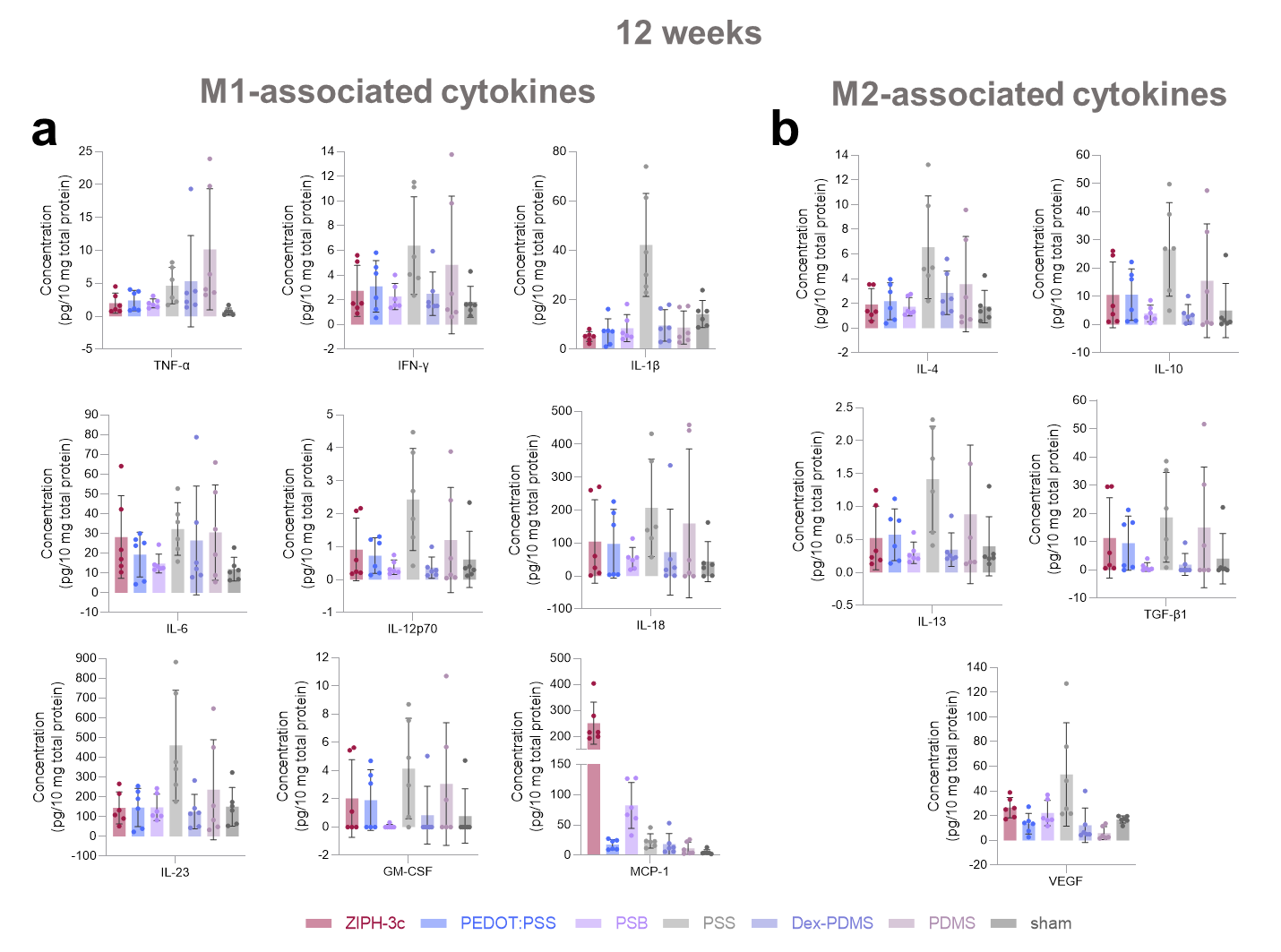
**Supplementary Figure 36.** Cytokine concentrations determined via LegendPlex assay divided into **a,** M1-associated cytokines and **b,** M2-associated cytokines 12-week post-implantation.

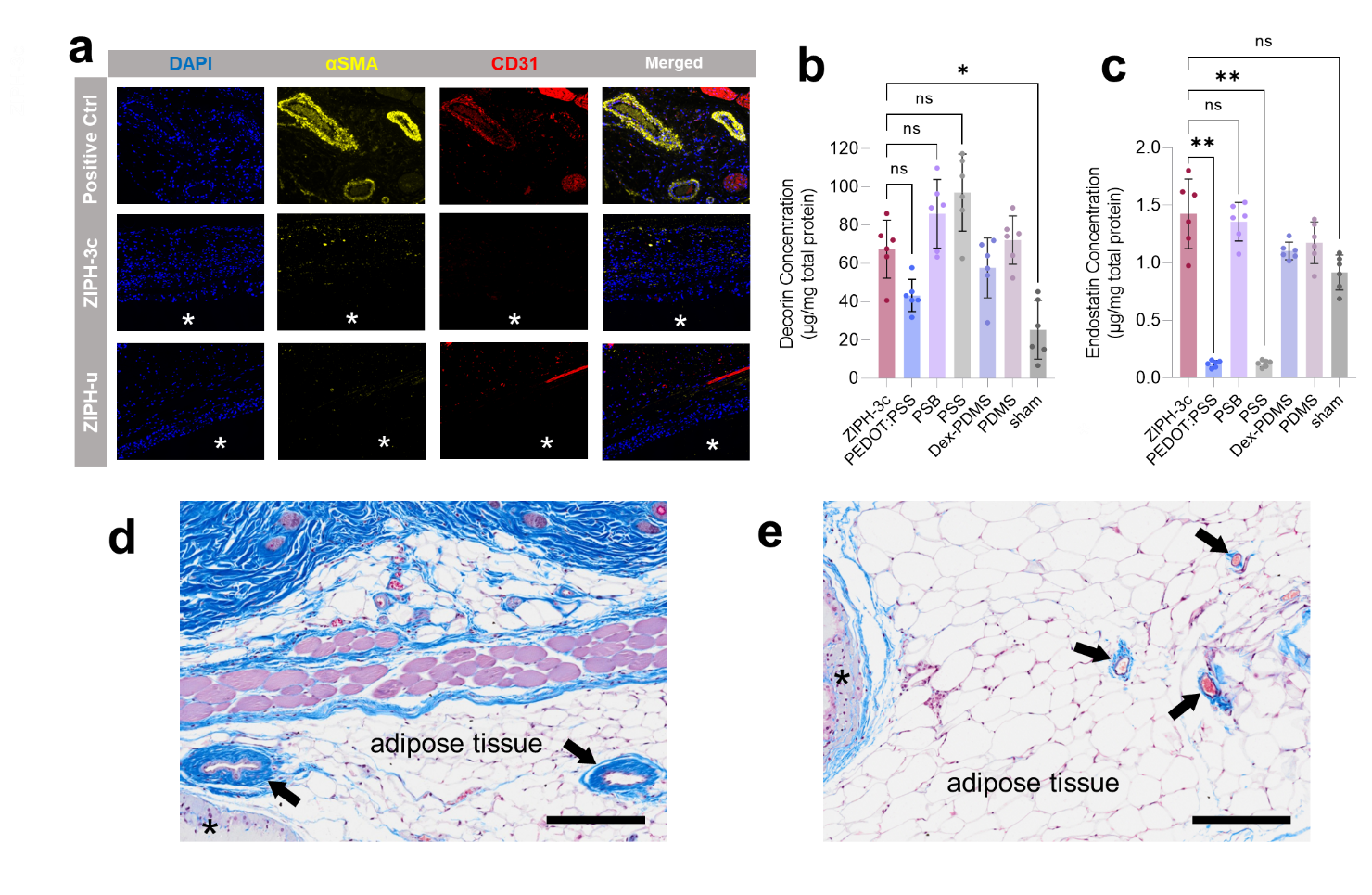
**Supplementary Figure 37.** Angiogenesis was not initially observed but becomes prominent at 24-weeks post-implantation. (**a**) Immunofluorescence microscopy images of ZIPH-3c and ZIPH-u stained with αSMA and CD31 at 4-weeks post-implantation. No angiogenesis is observed in the peri-implant region. The papillary dermis was used as the positive control. Implant locations are marked with asterisks. (**b**, **c**) Concentrations of endogenous angiogenesis inhibitors decorin (**b**) and endostatin (**c**) determined with ELISA 2-weeks post-implantation, when the concentrations of MCP-1 and VEGF were at their peaks. ZIPH-3c is associated with some of the highest concentrations of decorin and endostatin. (**d**, **e**) Masson’s trichrome stained tissues from the peri-implant region of ZIPH-3c at 24-weeks post-implantation. Blood vessels are marked with arrows. Implant locations are marked with asterisks. Scale bar, 200 μm.

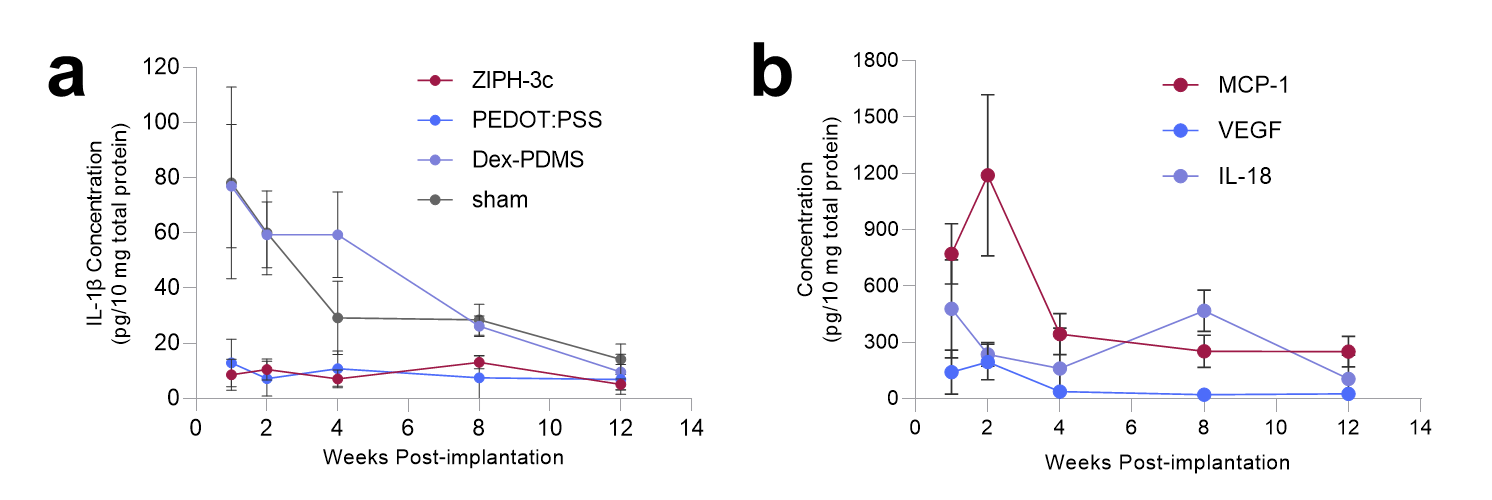

**Supplementary Figure 38.** Peri-implant concentrations of key cytokines. **a**, Concentration of IL-1β over time for ZIPH-3c, PEDOT:PSS, Dex-PDMS, and sham. **b**, Concentrations of MCP-1, VEGF, and IL-18 over time for ZIPH-3c.

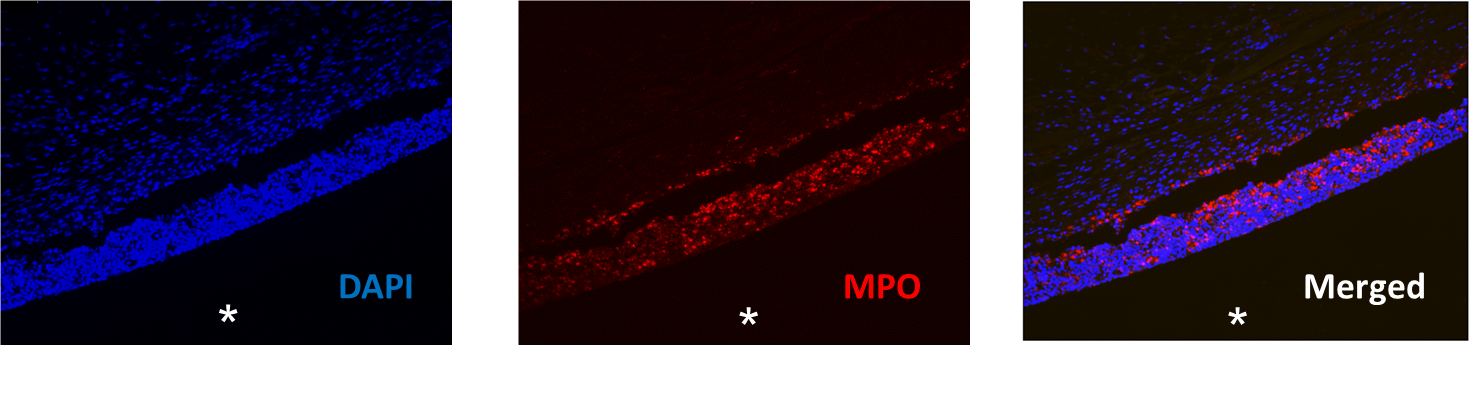
**Supplementary Figure 39.** Tissue section of PSB 1-week post-implantation stained for DAPI and myeloperoxidase (MPO). MPO is a marker for neutrophils, confirming that the high number of neutrophils associated with PSB 1-week post-implantation are associated with the implant. Implant location is marked with an asterisk.

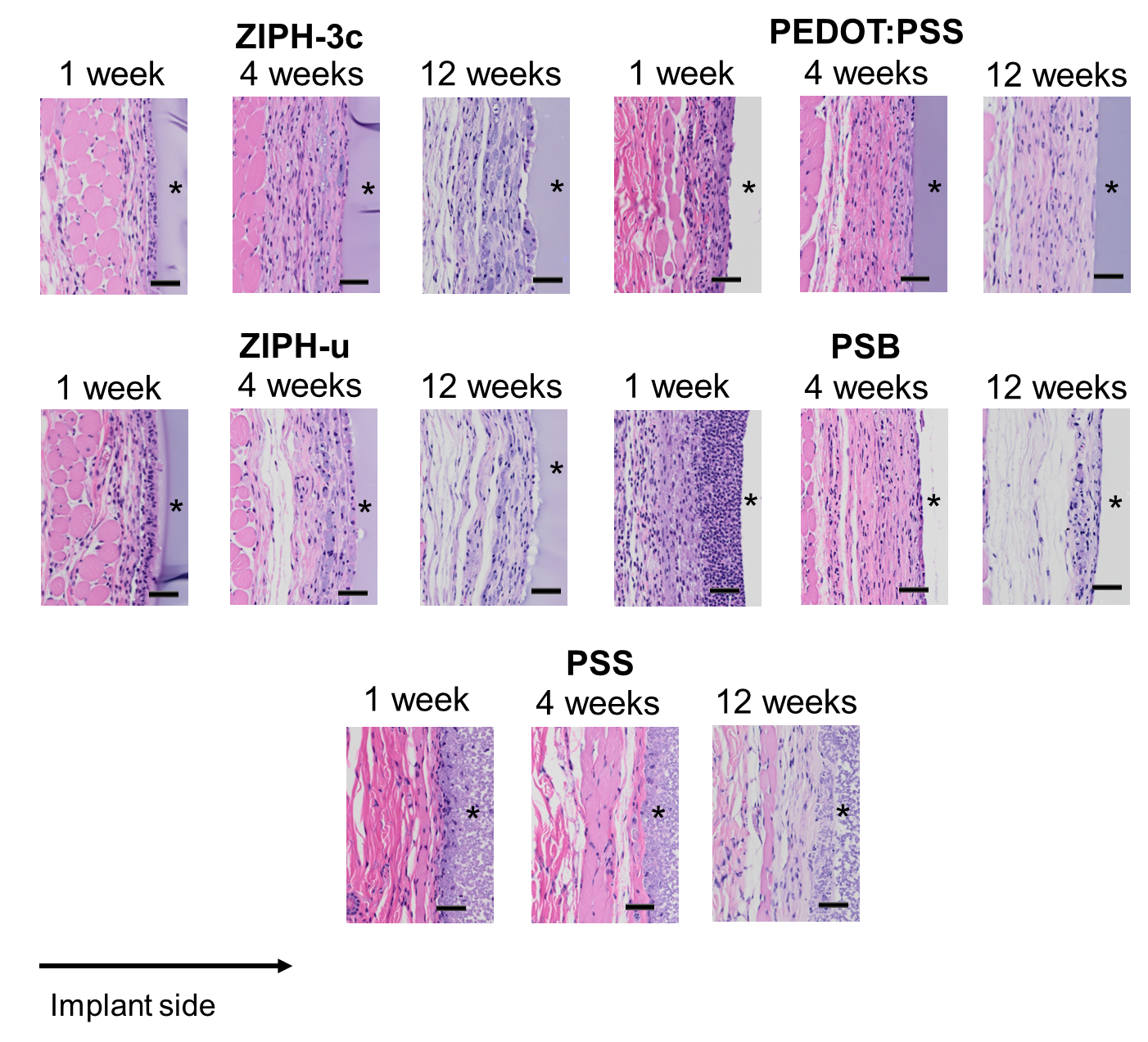

**Supplementary Figure 40.** Representative H&E-stained tissue sections 1-, 4-, and 12-weeks post-implantation. Scale bars, 50 μm.

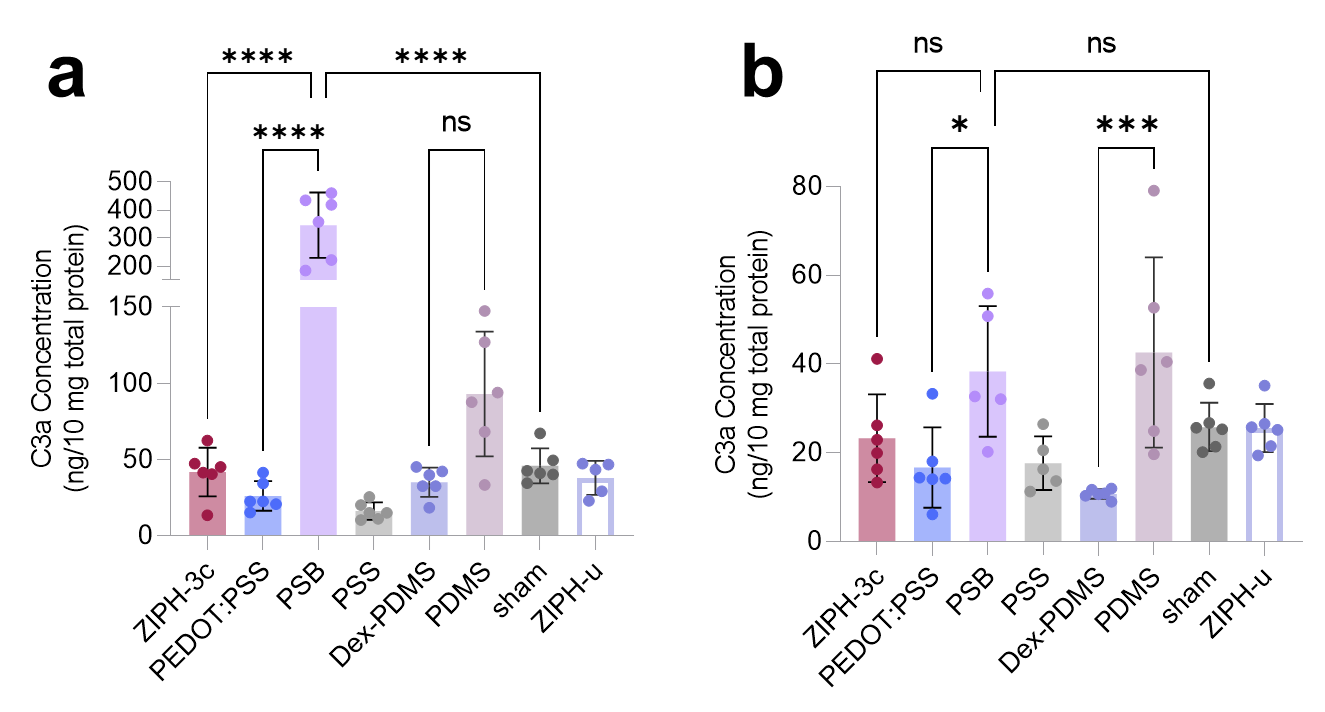
**Supplementary Figure 41.** C3a concentration determined via ELISA **a,** 1-week post-implantation and **b,** 4-week post-implantation.

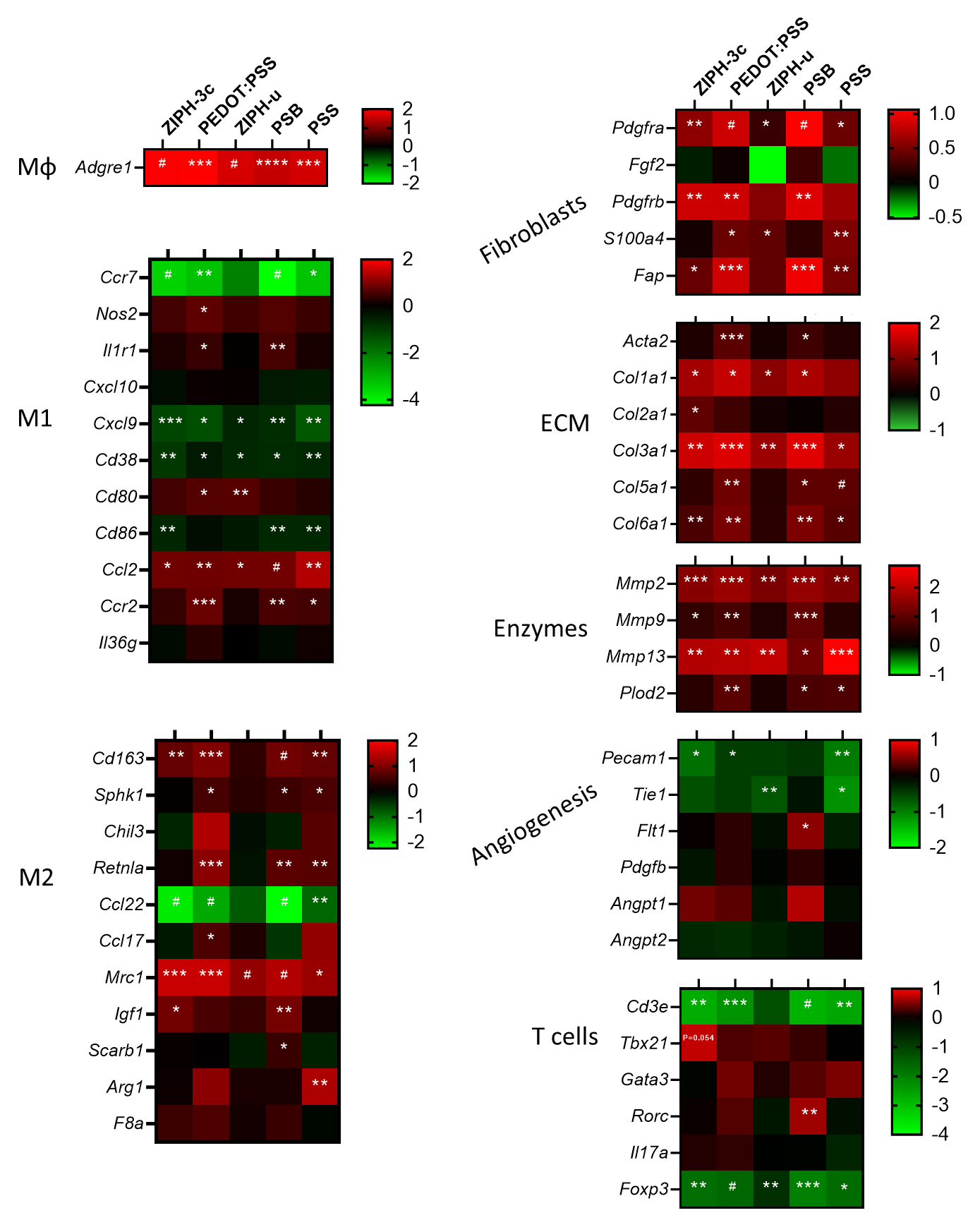
**Supplementary Figure 42.** Heat maps of RNA expression normalized to sham control 4-weeks post-implantation. Data were acquired using NanoString. N=4-5 mice per treatment. *, P < 0.05; **, P < 0.01; ***, P < 0.001; #, P < 0.0001 compared to sham.

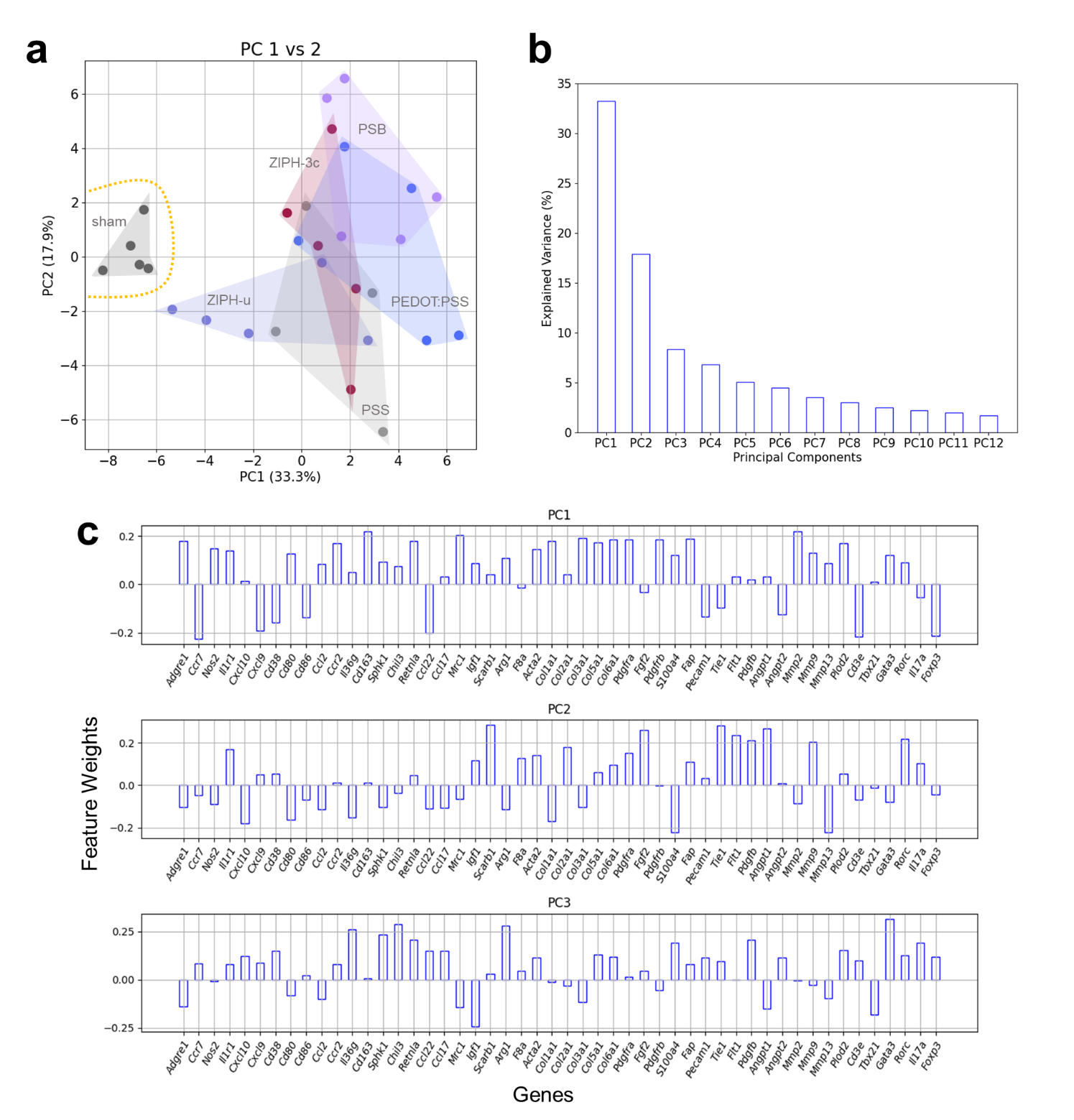
**Supplementary Figure 43.** Principle component analysis (PCA) of 50-genes, acquired via NanoString, 4-weeks post-implantation. **a,** Each sample plotted on PC1-PC2 vector space. **b,** Top 12 PCs and their explained variances. **c,** Genes and their associated weights for each PC.

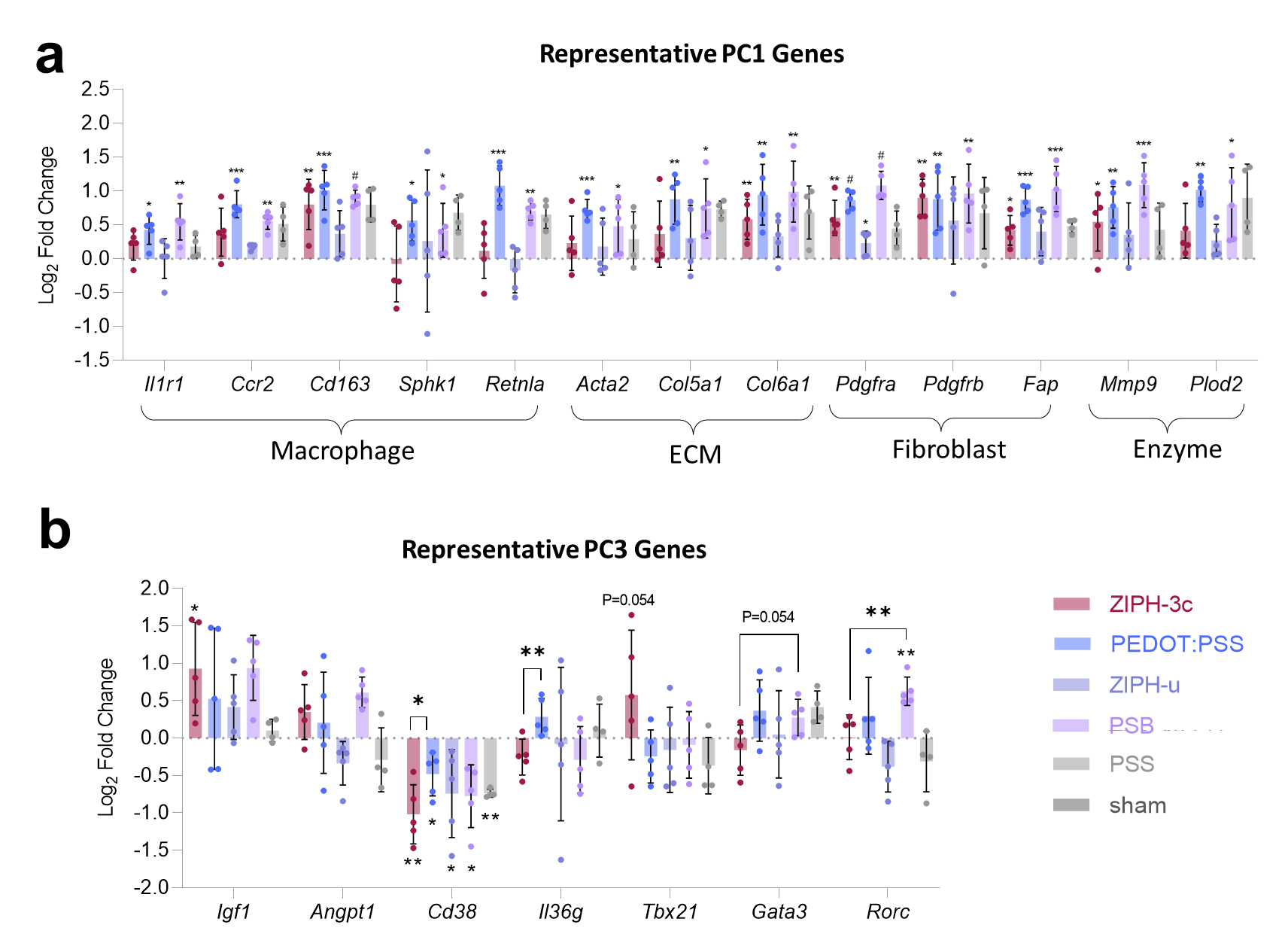

**Supplementary Figure 44.** Representative genes for all implant types tested for **a,** PC1 and **b,** PC3

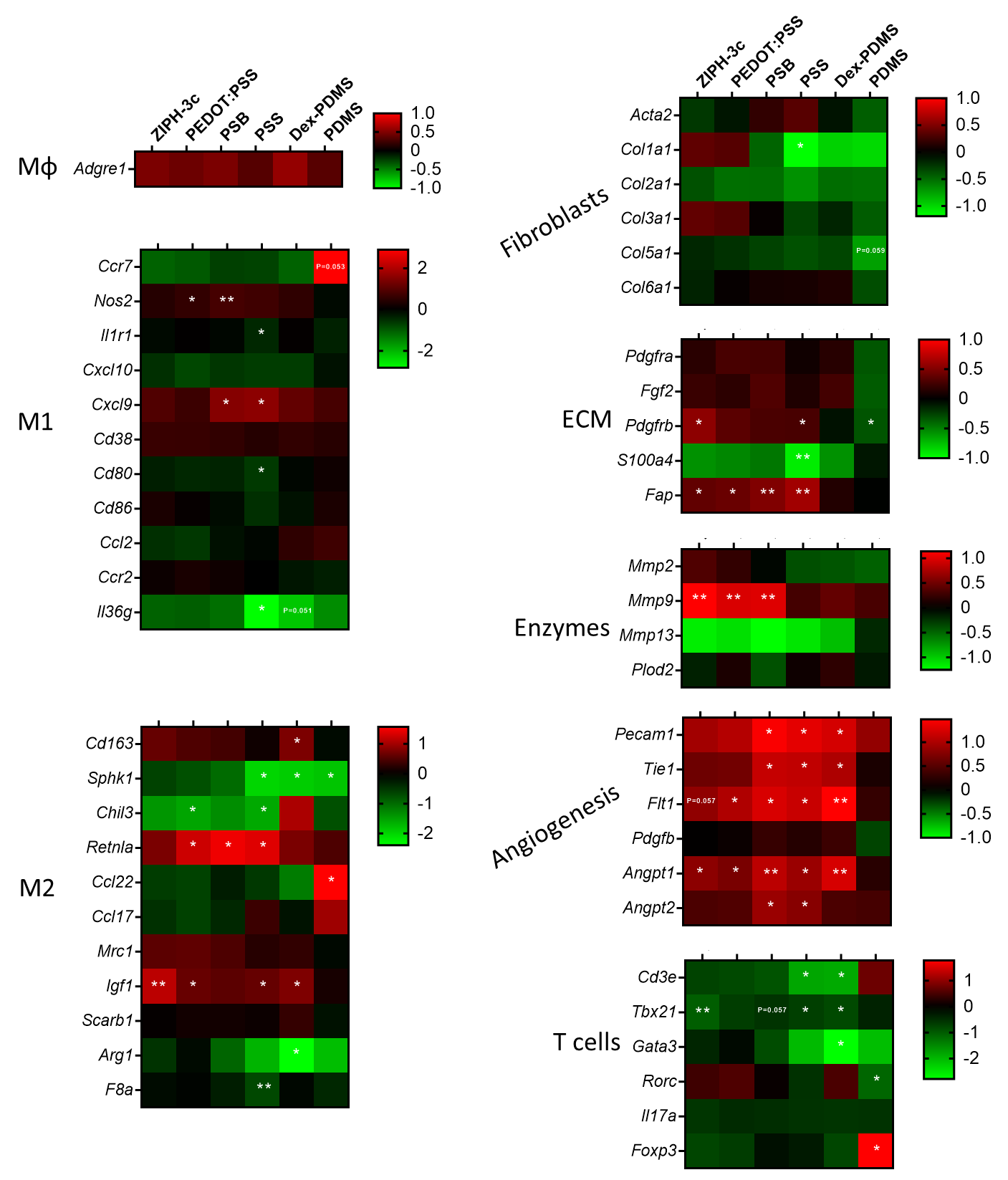
**Supplementary Figure 45.** Heat maps of RNA expression normalized to sham control 12-weeks post-implantation. Data were acquired using NanoString. N=4-5 mice per treatment. *, P < 0.05; **, P < 0.01 compared to sham.

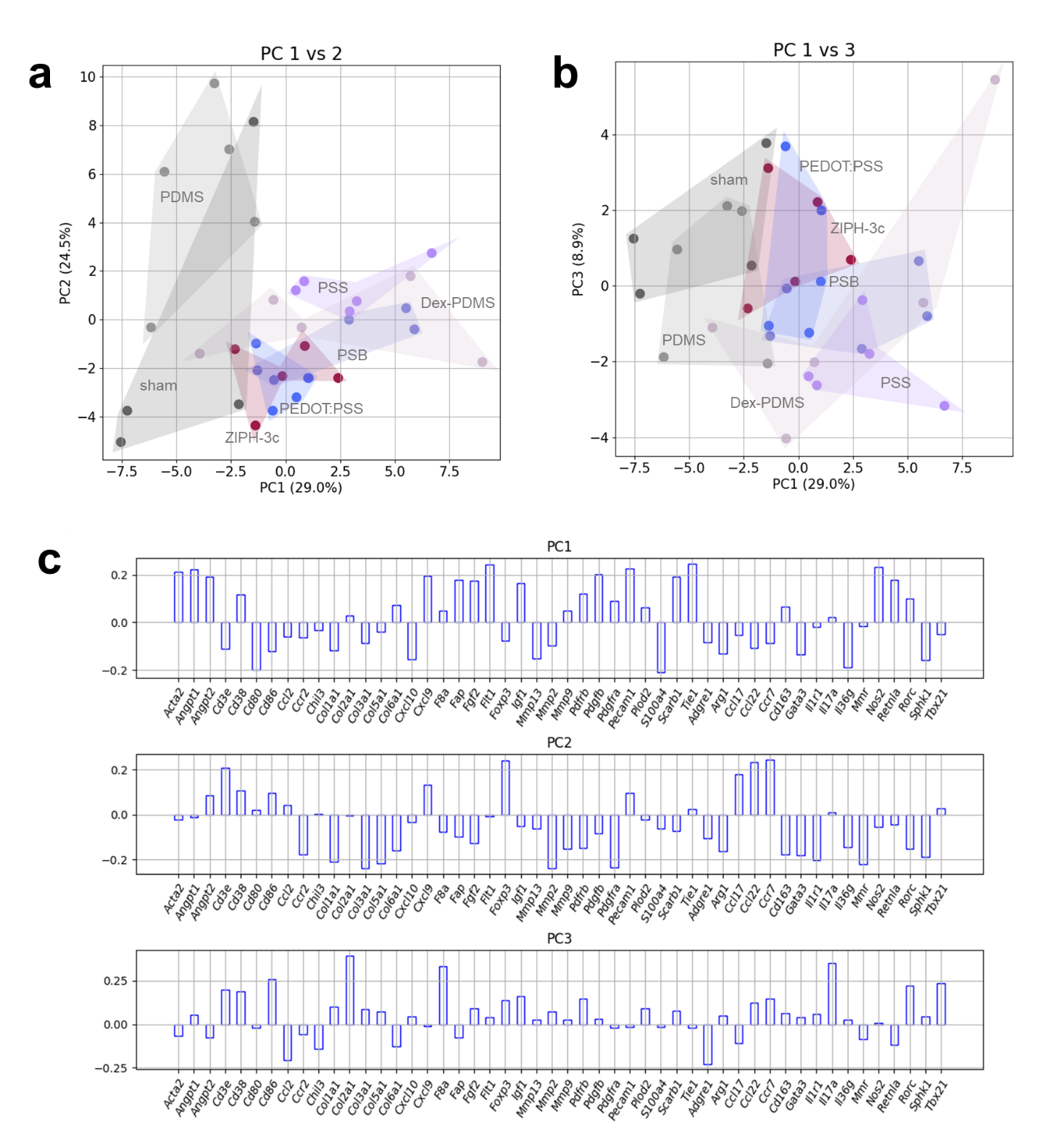

**Supplementary Figure 46.** Principle component analysis (PCA) of 50-genes, acquired via NanoString, 12-weeks post-implantation. **a,** Each sample plotted on PC1-PC2 vector space. **b,** Each sample plotted on PC1-PC3 vector space. **c,** Genes and their associated weights for each PC.

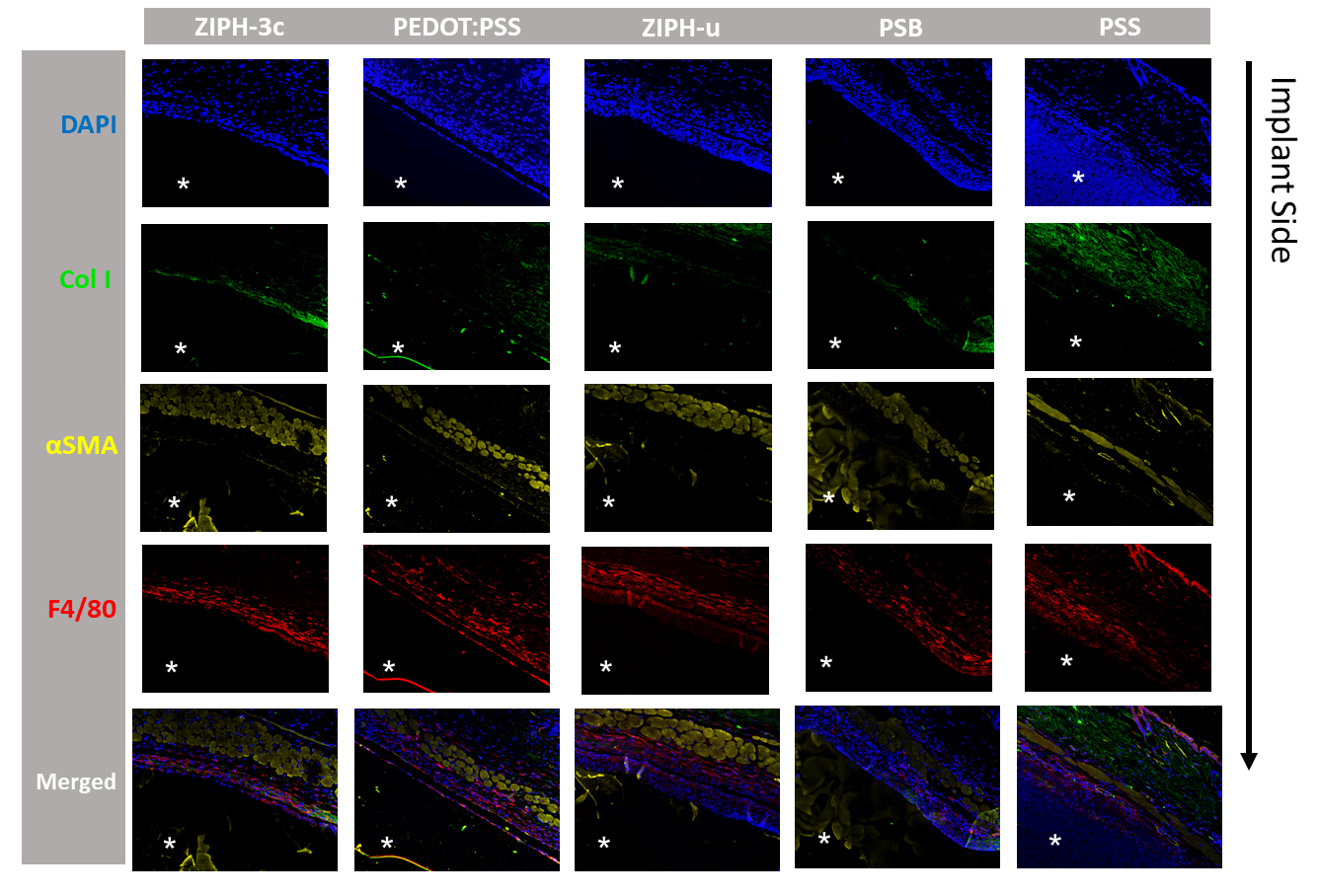
**Supplementary Figure 47.** Immunofluorescence microscopy images of tissue sections stained for markers associated with fibrosis 1-week post-implantation. Implant locations are marked by asterisks.

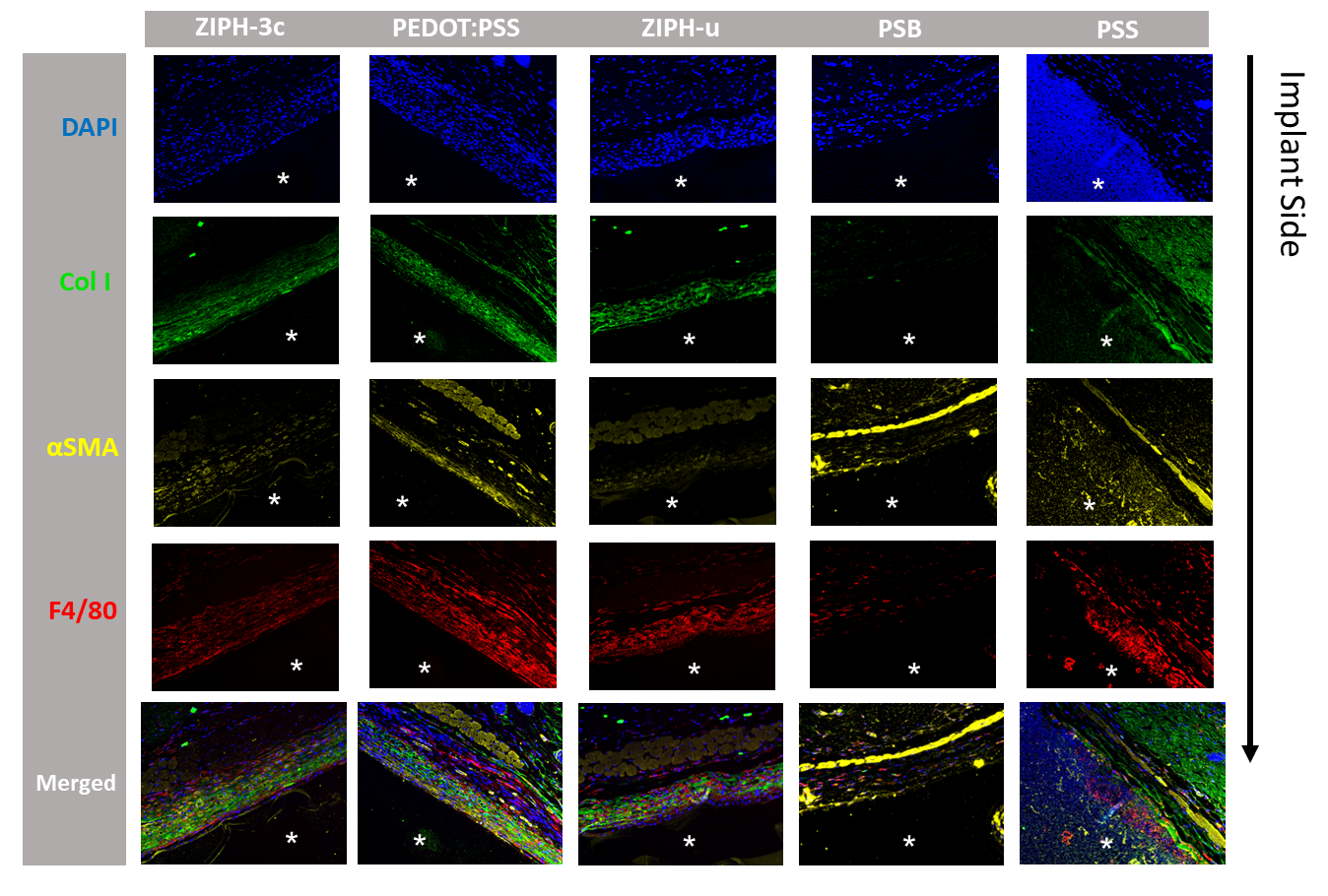

**Supplementary Figure 48.** Immunofluorescence microscopy images of tissue sections stained for markers associated with fibrosis 4-weeks post-implantation. Implant locations are marked by asterisks.

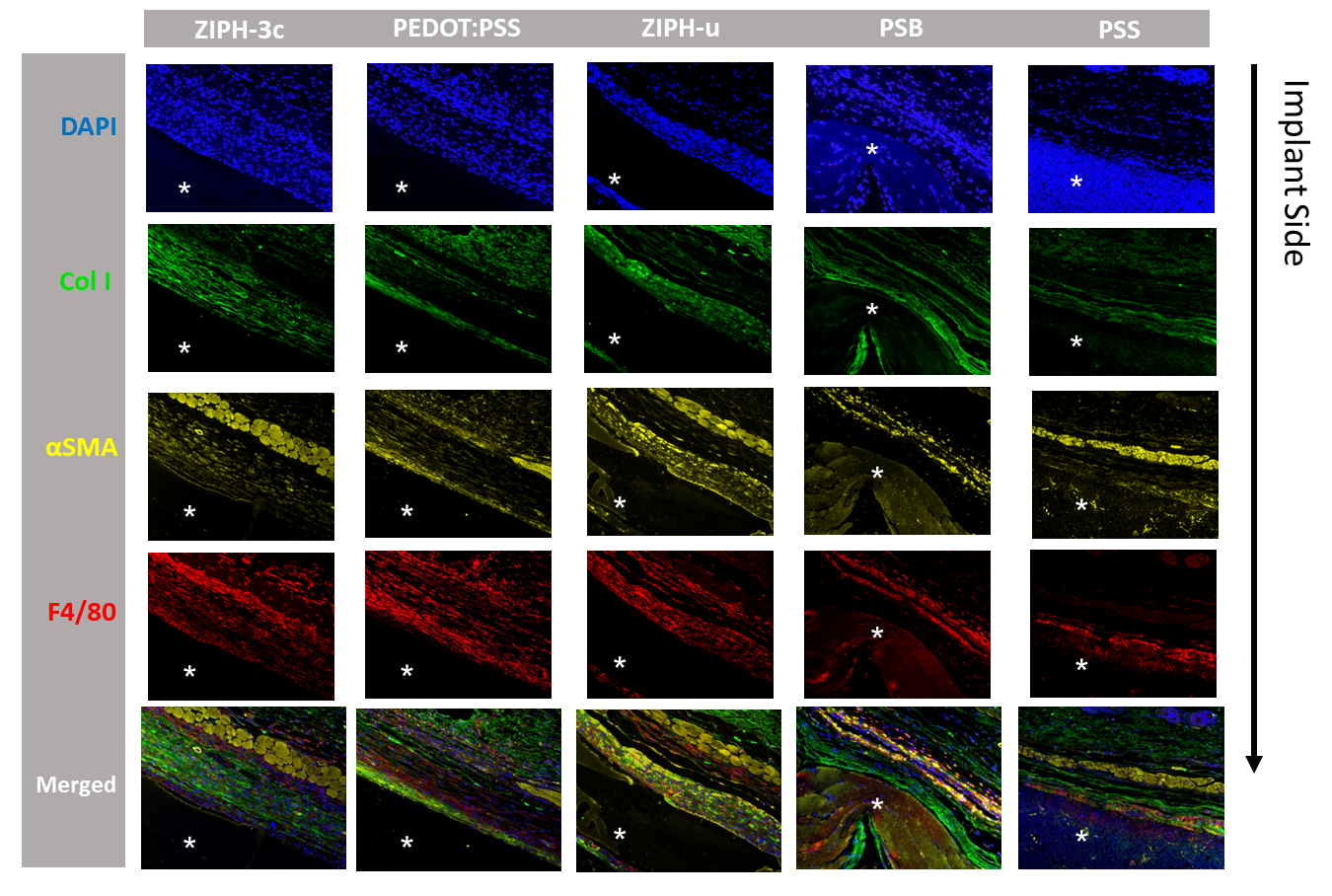
**Supplementary Figure 49.** Immunofluorescence microscopy images of tissue sections stained for markers associated with fibrosis 12-weeks post-implantation. Implant locations are marked by asterisks.

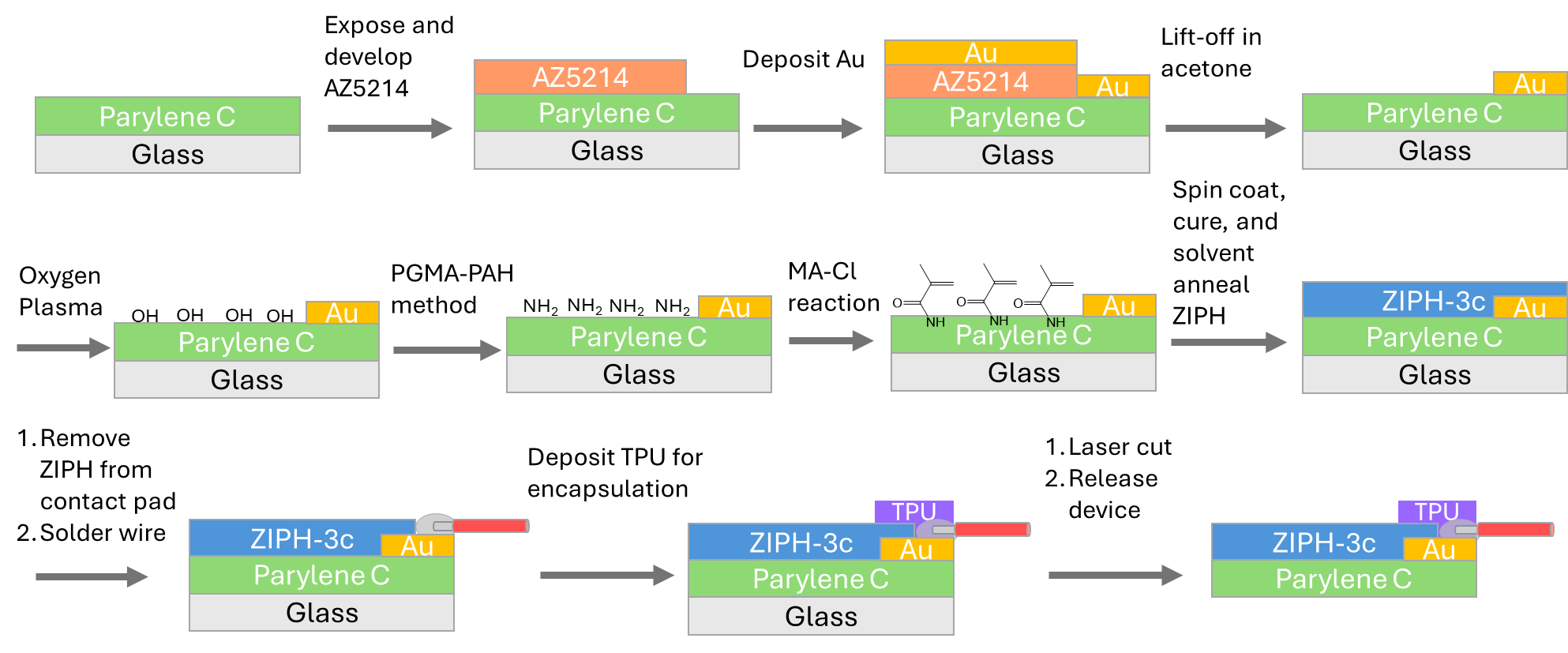

**Supplementary Figure 50.** Fabrication procedures for ZIPH-3c electrodes. PGMA, poly(glycidyl methacrylate); PAH, polyallylamine; MA-Cl, methacryloyl chloride.

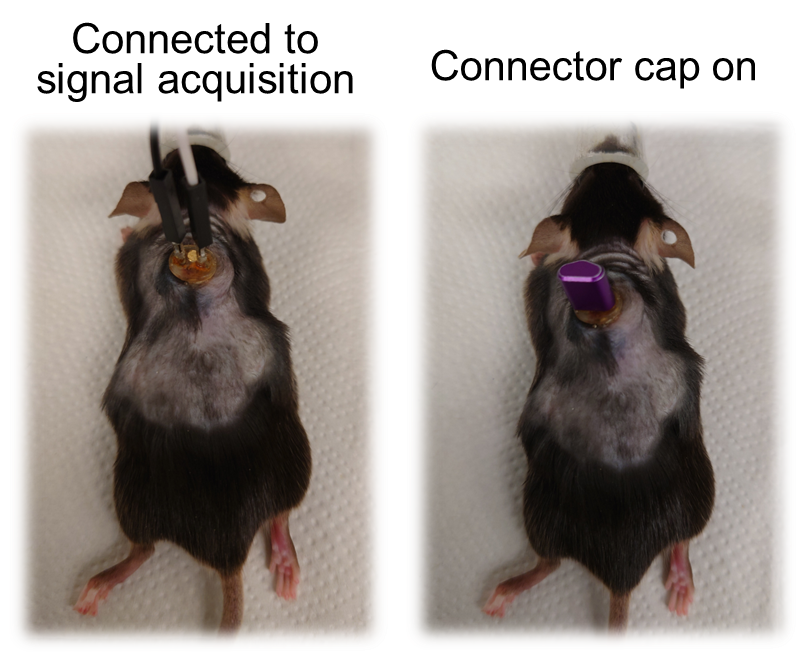

**Supplementary Fig. 51.** Anesthetized mouse with electrodes connected to peripheral electronics (left) and disconnected mouse with a cap on to protect the connector ports (right).

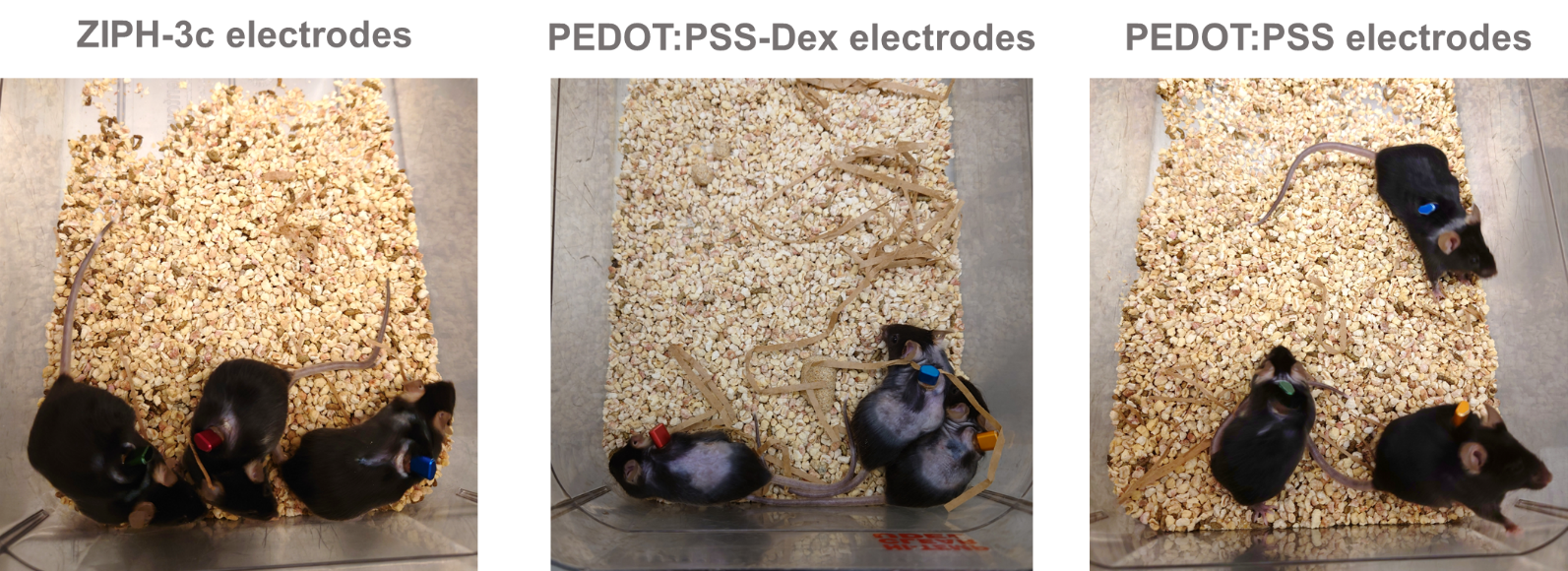

**Supplementary Figure 52.** Mice after 12-weeks post-implantation of ECG electrodes. Most of the fur grew back for mice implanted with ZIPH-3c and PEDOT:PSS electrodes, but this was not the case for mice implanted with PEDOT:PSS-Dex electrodes.

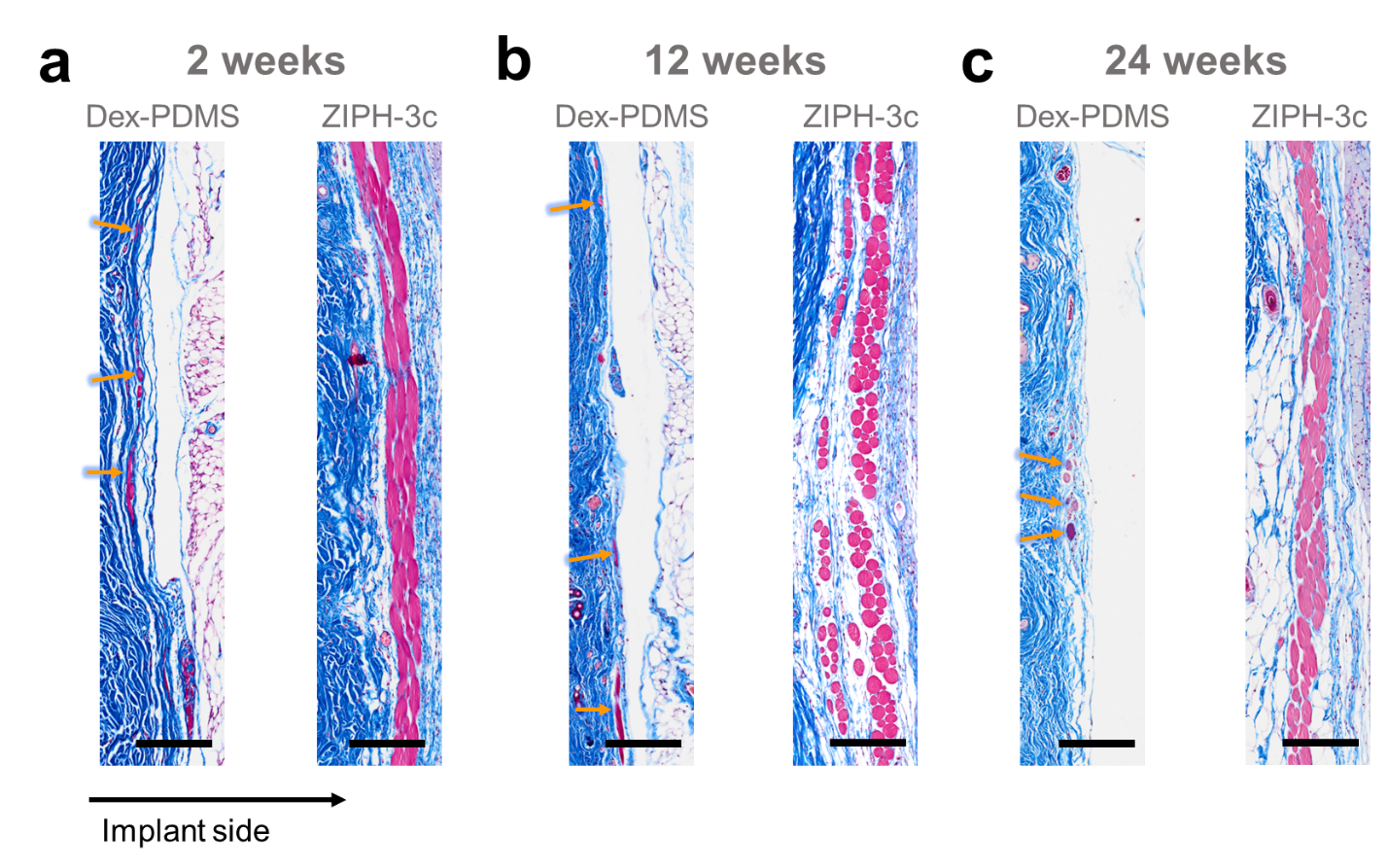

**Supplementary Figure 53.** Atrophy of the panniculus carnosus adjacent to the implant is observed for Dex-PDMS but not for ZIPH-3c. Images of the panniculus carnosus adjacent to the Dex-PDMS and ZIPH-3c implants at 2-week (**a**), 12-week (**b**), and 24-week (**c**) post-implantation. Arrows indicate the remains of atrophied panniculus carnosus. Scale bar, 200 μm.

| Material | Young’s Modulus (kPa) |
| --- | --- |
| PEDOT:PSS Hydrogel | 720 |
| ZIPH-3c | 9.9 |
| ZIPH-E | 1.4 |
| ZIPH-u | 8.7 |
| PSB | 9.0 |
| PSS | 9.7 |
| PDMS | 9.8 |
| Dex-PDMS | 10 |

**Supplementary Tables**

**Supplementary Table 1.** Young’s moduli of all materials used in *in vivo* experiments. Young’s moduli were calculated from tensile tests.

| **Double OR Single Network** | **Conductive Network** | **Secondary Network / Additive** | **Conductivity (S/cm)** | **Young’s Modulus** | **Biocompatibility (testing method)** | **Refs** |
| --- | --- | --- | --- | --- | --- | --- |
| Double | PEDOT:PSS | PSB / EBSA | 2.5 (wet); 80 (dry) | 9.9 kPa | 64% decrease in collagen density compared to pure PEDOT:PSS (MTS) | This work |
|  |  | PSB | 6 | 7.25 kPa | Less inflammation compared to gold (H&E) | ^83^ |
|  |  | Poly(acrylic acid) / DMSO | 247 | 25 kPa | Less fibrosis compared to stainless steel (H&E) | ^84^ |
|  |  | Poly(acrylic acid) / ionic liquid | 0.23 | 99 kPa | N/A | ^85^ |
|  |  | Poly(vinyl alcohol) | 10 | 460 kPa | Similar inflammation compared to gold (H&E) | ^86^ |
|  |  | Alginate | 0.087 | 1 MPa | N/A | ^87^ |
|  | Polyaniline | Poly(acrylic acid) | 0.05 | 50 kPa | N/A | ^88^ |
|  | Polypyrrole | Agarose | 0.7 | 27 kPa | N/A | ^89^ |
|  | Gold | Polyacrylamide, poly(acrylic acid), or poly(vinyl alcohol) | 520 | 2-5 MPa | Thick fibrotic capsule and high inflammation (MTS, H&E) | ^90^ |
| Single | PEDOT:PSS | Ionic liquid | 47 | 28 kPa | Less inflammation compared to PET (IF) | ^28^ |
|  |  | *D*-sorbitol | 74 | 2.6 MPa | Less inflammation compared to platinum (H&E) | ^91^ |
|  |  | DMSO | 20 | 2 MPa | N/A | ^92^ |

**Supplementary Table 2.** Comparison of conductivity, Young’s modulus, and biocompatibility of electrically conductive hydrogels from our work and previously reported works.
